# Biomechanical Compartmentalisation Influences Angiogenesis and Stem Cell Proliferation in the Regenerating Intestine via Vascular Piezo

**DOI:** 10.1101/2025.04.16.649133

**Authors:** Jade A. Phillips, Jessica Perochon, Cai T. Johnson, Matthew Walker, Colin Nixon, Mark Hughes, André Barros-Carvalho, Yachuan Yu, Louise Mitchell, Karen Blyth, Massimo Vassalli, Julia B. Cordero

## Abstract

The vasculature is a prominent component of developmental and adult tissue microenvironments. How tissue specific characteristics and environmental states influence vascular biology and function, remains largely understudied.

Using the fruit fly *Drosophila melanogaster* midgut and its associated vascular-like tracheal system, we discover reciprocal, mechanochemical interorgan communication between the adult intestine and its vascular microenvironment, which influences vascular and stem cell adaptations during intestinal regeneration. Mechanistically, apoptotic epithelial cells contribute to biomechanical remodelling of the regenerating midgut, which is associated with activation of Piezo in a subset of gut tracheal cells. Piezo drives activation of the mechanosensitive transcription factor Yorkie/YAP, leading to tracheal remodelling and intestinal stem cell proliferation. Furthermore, we identify a conserved role of mammalian vascular *Piezo1* influencing remodelling of the intestinal crypt vasculature, and intestinal regeneration upon DNA damage and inflammation in mice.

Our cross-species *in vivo* study reveals the presence of biomechanically distinct compartments in the regenerative intestine and identifies mechanosensory interorgan signalling, which contributes to intestinal regeneration through the vascular stem cell niche. Furthermore, our work highlights the importance of studying tissue- and context-specific vascular biology to understand compartmentalization, plasticity, and the complexity of tissue/vasculature interactions within a living organ.

## Introduction

The adult intestine is a remarkably plastic organ, capable of responding to a myriad of environmental stimuli through intrinsic cellular adaptations and complex interactions with a highly heterogeneous microenvironment ^1-3^. How, specialised interactions between the intestinal epithelium and its diverse microenvironment contribute to tissue plasticity and the functional versatility of the adult intestine, is poorly understood.

The vasculature interacts with target tissues through the exchange of oxygen, metabolites and angiocrine factors ^4-6^. Beyond supporting organ specific functions, the vascular system of highly self-renewing and plastic tissues, such as the intestine and the skin, must adjust to significant tissue remodelling processes, which range from discrete cellular adaptations to whole organ resizing, involving profound chemical and biophysical perturbations ^7-13^. While chemical crosstalk between tissues and their vasculature is abundantly studied, less is known about how changes in tissue physical state are sensed and translated into adaptive responses within the vascular microenvironment.

Interactions between the adult intestine and its vascular microenvironment are reflected in the crosstalk between the midgut and the rich tracheal network in the fruit fly *Drosophila melanogaster* ^7-10^. The *Drosophila* tracheal system is a mesh-like structure composed of oxygen-delivering tubes, functionally and molecularly alike the mammalian blood vasculature and respiratory system ^14, 15^. *Drosophila* terminal tracheal cells are related to vertebrate vascular tip cells ^16^ and extend prominent cytoplasmic projections to supply oxygen to their target tissues ^14, 17, 18^. Previous work from us and others showed a chemical crosstalk between the intestinal epithelium and the vasculature like tracheal tissue in *Drosophila* during pathogen-induced intestinal regeneration and tumour growth triggered by reactive oxygen species (ROS) dependent signalling ^7, 8^. However, chemical stress signals alone are insufficient to explain the broad range of tracheal/midgut interactions observed in the adult fly midgut ^7, 8, 10^ or adaptations of the mammalian vascular network in broad physiological settings ^11-13^. Here, we used fly and mouse model systems, combined with live imaging, quantitative biology and biophysical approaches, to investigate how regenerative changes in tissue physical state are sensed by the vascular microenvironment and translated into adaptive responses during intestinal regeneration.

Results from our *in vivo* studies provide a framework for understanding how tissue-derived mechanosensory signals contribute to vascular adaptation across physiological and pathological contexts.

## Results

### Changes in cellular and tissue mechanics in the regenerating intestine correlate with remodelling of associated tracheal/vascular tissue

Feeding flies with pathogenic bacteria causes a significant disruption to the gut’s normal function and structure, including stress signal activation, epithelial cell death and compromised organismal heath ^19-21^. The dense network of terminal tracheal cells (TTCs) populating the *Drosophila melanogaster* midgut and labelled with *GFP* driven under the Serum Response Factor promoter (*dSRFGal4^ts^>GFP*), remodels through ROS induced cross-tissue signalling during damage-induced midgut regeneration by oral pathogenic infection and oxidative stress ^7, 8^ (Fig. 1a, b and Supplementary Fig.1a-c). Active tracheal remodelling and regenerative ISC proliferation is similarly observed in midguts exposed to 48-hour treatment with dextran sodium sulphate (DSS), which induces intestinal damage and compensatory intestinal stem cell proliferation by disrupting the basal membrane of the intestinal epithelium and inducing extra cellular matrix remodelling ^22, 23^ (Supplementary Fig. 1d-f). Damage to the midgut epithelium upon 16-hour oral exposure with the entomopathogen *Pseudomonas entomophila* (*Pe*) generates high levels of ROS in the gut lumen, mainly via enterocytes (ECs), as a protective mechanism from the host against the pathogen ^24^ (Supplementary Fig. 1g, h). On the other hand, 48-hour DSS treatment does not affect basal levels of ROS in the midgut (Supplementary Fig. 1i, j). Of note, basal levels of ROS detected in midguts from the DSS treatment group and their matched vehicle control (Sucrose) were overall higher than those in Sucrose midguts from the *Pe* treatment group, possibly due to longer incubation times with the vehicle and damaging agent in the DSS treatment group. Importantly, treating flies with the antioxidant N-acetyl cysteine (NAC) alongside DSS, had no significant effect on DSS induced tracheal remodelling or ISC proliferation (Supplementary Fig. 1k-m). Such observations suggested that, remodelling of the adult gut associated trachea, is driven by multiple signals, and cannot be explained solely by the presence of oxidative stress. Understanding these additional mechanisms of gut-tracheal communication may provide insight into how diverse tissue-derived signals regulate vascular remodelling across physiological and pathological contexts^11-13, 25-27^.

**Figure 1.**
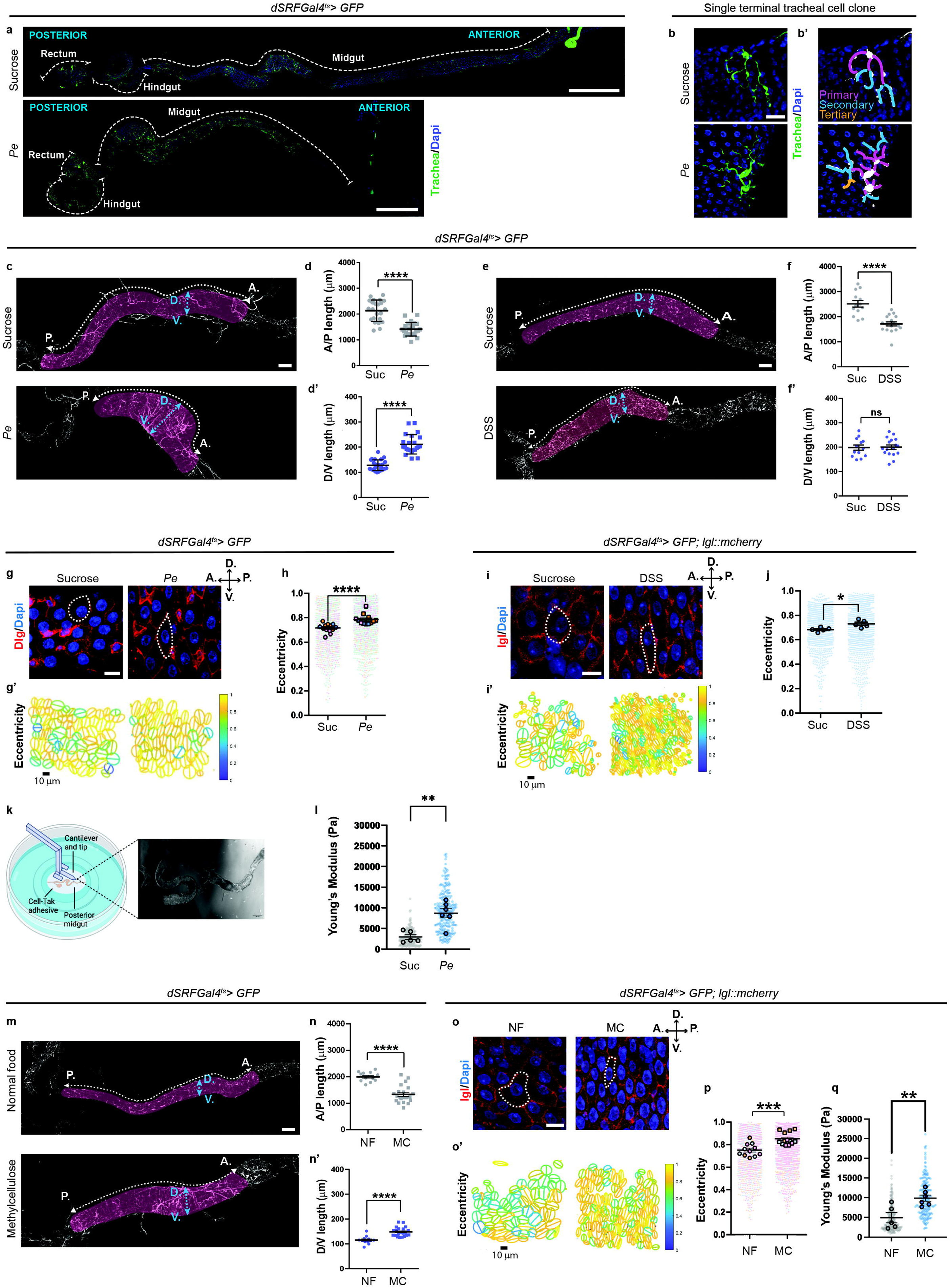
Morphological, geometrical and biomechanical changes in the regenerating adult *Drosophila* intestine correlate with tracheal remodelling. **a.** Adult midgut segments and associated trachea (green) upon sucrose or *Pe* feeding (n=1, 1). Scale bars, 500 μm. b. Examples of single tracheal cell clones from sucrose or *Pe* midguts (n=1, 1). Scale bars, 20 μm. b’. Segmentation of primary (purple), secondary (blue) and tertiary (orange) branches in cells from b (n=1, 1). c. Posterior midgut (shaded red) with anterior-posterior (A.P.) and dorsal-ventral (D.V) lengths indicated (arrows). Scale bar, 100 μm. d, d’. Quantification of A.P. (d) and D.V. (**d’**) lengths as in c (n=23, 23). **e**. Midguts upon sucrose or DSS feeding. Scale bar, 100 μm. **f, f**’. Quantification of A.P. (**f**) and D.V. (**f’**) lengths as in e (n=13,17). **g**. Intestinal epithelium upon sucrose or *Pe* feeding; anti-Dlg (apical located septate junctions; red). Scale bar, 10 μm. **g’**. Epithelial eccentricity as in f. Scale bar, 10 μm. **h**. Quantification of eccentricity as in g’; small circles indicate individual cells, large circles indicate per-gut averages, colours denote biological replicates (n=13, 12). **i**. Intestinal epithelium with sucrose feeding or DSS treatment. *lgl*::mcherry septate junction reporter (red). Scale bar, 10 μm. **i’**. Eccentricity as in n. Scale bar, 10 μm. **j**. Quantification as in i’ (n=5,5). **k.** Nanoindentation setup. *Created in BioRender. Cordero, J. (2026)* https://BioRender.com/z99l168. **l,** Stiffness quantification (R4 region) upon sucrose and *Pe* feeding (**l**) (n=5,6); small circles represent individual measurements, large circles represent per-gut averages. **m**. Midgut upon normal food (NF) or MC feeding. Scale bar, 100 μm. **n, n’**. Quantification of A.P. (**n**) and D.V. (**n’**) lengths (n=12,22). **o.** Intestinal epithelium following NF/MC feeding. Scale bar, 10 μm. **o’**. Eccentricity as in v. Scale bar, 10 μm. **p**. Quantification of eccentricity as in p’ (n=11,13). **q**. Stiffness quantification (R4 region) upon NF/MC feeding (n=5, 6). Data presented as mean ± SEM. n values indicate the numbers of guts analysed. DAPI labels nuclei (blue). <u>Statistics</u>: Two-tailed unpaired Student’s t-test.

Organ reshaping is a recognised, yet understudied, property of regenerating midguts ^7, 28^. Morphological analysis of control and damaged midguts revealed increased length of the dorsal-ventral (D/V) axis and/or shortening of the anterior-posterior (A/P) axis in posterior midguts during *Pe* or DSS treatment (Fig. 1c-f’). To understand cellular features associated with the observed organ shape changes, we used available tools for image segmentation and morphometric analysis ^29^ to quantify cell morphology parameters within the midgut epithelium. Measurements of epithelial cellular shape obtained from masks of segmented midgut enterocytes labelled with anti-Discs Large (Dlg) or a *lethal giant larvae* (*lgl*) reporter, which mark apically located septate junctions in the midgut epithelium ^30^ revealed increased presence of elongated or stretched, *versus* circular, enterocytes (ECs)⎯ characterised as eccentricity⎯ in *Pe* (Fig. 1g-h) and DSS treated midguts (Fig. 1i-j).

Cell shape emerges from the interplay between cellular programmes and the mechanical environment of tissues. In epithelial tissues, changes in force balance, tension anisotropy and tissue architecture are often reflected in cell morphology, packing and organisation ^31-36^. Accordingly, increased cellular eccentricity is commonly associated with altered biomechanical states, although it represents a geometric descriptor rather than a direct measurement of force or stress ^37-42^. To independently assess whether intestinal regeneration was accompanied by changes in tissue mechanical properties, we next performed cantilever-based nanoindentation of freshly dissected midguts (Fig. 1k). Measurements revealed increased tissue stiffness in midguts exposed to 16-hour *Pe* compared with their respective controls (Fig. 1l). Together, these observations indicate that damage-induced intestinal regeneration is accompanied by coordinated changes in epithelial cell geometry and tissue mechanical properties, consistent with the emergence of an altered biomechanical state within the regenerating intestine. Of note, comparable nanoindentation experiments upon 48-hour Sucrose and DSS treatements were unsuccessful due to the overall fragility of the guts, which made *ex-vivo* organ manipulations for nanoindentaion difficult, precluding reliable assessment of organ stiffness at that experimental stage.

To investigate whether perturbations to the mechanical environment of the intestine could reproduce features of the regenerating gut, we fed flies with methylcellulose (MC), an indigestible fibre previously shown to induce gut expansion and activation of mechanosensitive signalling within the intestinal epithelium^43^. MC feeding alone recapitulated several morphological changes of regenerating midguts, including shortening of the A/P length of the gut tube, lengthening of the D/V axis, increased EC eccentricity and midgut stiffness (Fig. 1m-q). Furthermore, midguts of MC fed animals showed significant increase in remodelling of associated terminal tracheal cells (TTCs) (Supplementary Fig. 1n, o). Consistent with previously published data, marginal levels of ISC were observed upon MC feeding ^43^ (Supplementary Fig. 1p). Together, these observations indicate that biomechanical perturbation of the intestine reproduces epithelial, tissue-level and tracheal phenotypes associated with regenerative conditions of the adult *Drosophila* midgut.

### Changes in enterocyte geometry during intestinal regeneration suggest compartmentalized mechanics

Prior studies have compartmentalised the adult intestine into histologically, molecularly and metabolically defined sub-regions ^44-46^ (Fig. 2a). We noticed that, despite the global nature of the gut epithelial damage used in our models of intestinal regeneration, phenotypes of tracheal remodelling were regionalised across the posterior midgut, with the most prominent increase in TTC branching observed in the R4 *versus* R5 segment of the posterior midgut from *Pe* and DSS fed animals (Fig. 2b-e). Interestingly, measures of A/P gut axis length or morphological changes in the visceral muscle across R4/R5 did not match the regional bias observed in tracheal remodelling (Supplementary Fig. 2a-f). On the other hand, EC aspect ratio⎯ a geometrical measure of cellular elongation equivalent to eccentricity⎯ was significantly increased in R4, upon *Pe* and DSS treatment, while remaining largely stable in the R5 segment (Fig. 2b-d, f). These observations suggested the existence of regional differences in the biomechanical state of the regenerating epithelium. To explore this possibility further, we adopted a previously validated Bayesian Force Inference approach that infers mechanical information from epithelial geometry. This methodology has been shown to infer relative differences in junction tensions, force balances and tissue-scale stress patterns in *Drosophila* epithelia ^47^. We therefore used it to compare the inferred mechanical state of the epithelium across the R4 and R5 regions from tile scan images of *Pe* or DSS treated and control midguts, using the R5 gut region as an internal control. Using this approach, we observed a mild but significant increase in the inferred cell-pressure parameter in the R4 region, relative to R5, following intestinal damage (Fig. 2g-i and Supplementary Fig. 2g, h). In contrast, this inferred parameter remained stable across compartments in homeostatic midguts (Fig. 2g-i and Supplementary Fig. 2g, h). Changes in epithelial force balance are often accompanied by changes in the apparent mechanical properties of tissues and cell collectives, reflecting the intimate coupling between tissue architecture, cellular tension and mechanical response^48^. We therefore asked whether the regional differences revealed by morphometric analysis and force inference were also associated with measurable changes in tissue biomechanics. Cantilever-based nanoindentation on intact gut tubes detected a significant increase in tissue stiffness specific to the R4 region of the posterior midgut upon *Pe* damage (Fig. 2j).

**Figure 2.**
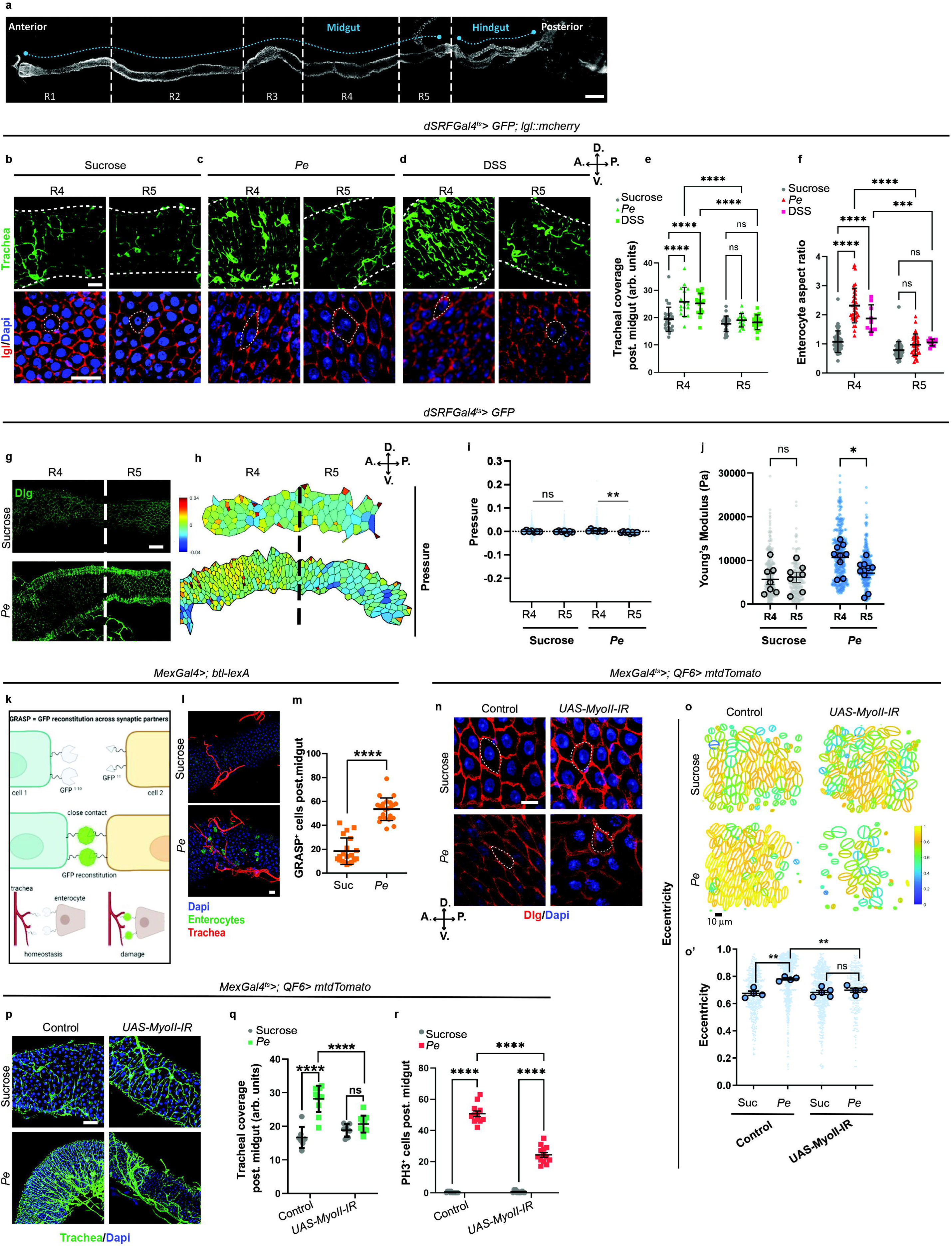
Enterocyte cell geometry and tissue biomechanical changes in the regenerating midgut. **a.** Adult midgut showing R1-R5 regions (n=1). Scale bar, 100 μm. **b-d**. Confocal images of trachea (green) and intestinal epithelium (lgl, red) in R4/R5 upon Sucrose (**b**), *Pe* (**c**) or DSS feeding (**d**). Scale bar, 20 μm. **e, f**. Quantification of tracheal coverage (**e**) and enterocyte aspect ratio (**f**) in R4/R5 as in b-d (n=29,16,18) (n=44,36,45; n= individual cell measured from guts as in e). **g**. Tile scans across R4/R5. Enterocytes septate junctions (lgl; green). Scale bar, 50 μm (n=1, 1; 2 biological replicates). **h**. Image of inferred relative cell pressure via Bayesian Force Inference derived from g. **i**. Quantification of relative pressure across R4/R5 regions under sucrose feeding or *Pe* infection. Small circles indicate individual cells, large circles indicate per-gut averages (n=7, 7, 8, 8). **j**. Stiffness quantification across R4/R5 regions via nanoindentation upon sucrose or *Pe* feeding. Small symbols indicate individual measurements. Large symbols indicate number of guts processed independently (n=7,7,9,9). **k**. GRASP system used to assess enterocyte/tracheal interactions. **l**. Confocal images with apical view of GRASP components’ expression in epithelium and trachea following sucrose or *Pe* feeding. Scale bar, 20 μm. **m**. Quantification of GRASP-positive cells per posterior midgut (n=20,23). **n**. Confocal images of enterocytes (Dlg, red) in control midguts or following enterocyte non-muscle Myosin II RNAi (*Mex^ts^>MyoII-IR*) upon sucrose or *Pe* feeding. Scale bar, 10 μm. **o**. Epithelial eccentricity in midguts as in a. Scale bar, 10 μm. **o’**. Quantification of eccentricity as in o; small circles indicate individual cells, large circles indicate per-gut averages (n=4,5,4,4). **p**. Confocal images of trachea (green) in wildtype midguts or *Mex^ts^> MyoII-IR* midguts under sucrose or *Pe* feeding. Scale bar, 20 μm. **q, r**. Quantification of tracheal remodelling (**q**) and ISC proliferation (**r**) in midguts as in p (n=7,11,7,10), (n=13,12,13,13). Data presented as mean ± SEM. Unless otherwise indicated, n=numbers of guts analysed. DAPI labels nuclei (blue). <u>Statistics</u>: **e, f, i, m**. Two-tailed unpaired Student’s t-test. **o, q, r.** Two-way ANOVA followed by Tukey’s multiple comparisons test. **j**, Two-way ANOVA followed by Sidak’s multiple comparison test.

Together, the regional increase in epithelial cell eccentricity, the force-inference analysis and the independent measurements of tissue stiffness suggest a previously unrecognised biomechanical compartmentalisation of the regenerating adult *Drosophila* midgut. These observations suggest that regenerative responses are associated with distinct regional biomechanical states, with the R4 compartment exhibiting the most pronounced changes.

### Enterocytes mediate biomechanical communication leading to tracheal remodelling within the regenerating midgut

Cellular projections of TTCs appear located between the visceral muscle and epithelial compartments of the adult midgut ^7^. We reasoned that, one way for trachea to respond to changes in cell geometry within the midgut epithelium, would be through physical contact with ECs. To test this, we used the GRASP (GFP reconstitution across synaptic partners) system ^49^ to visualise points of physical proximity between tracheal cells and ECs by expressing the 11^th^ strand of a split GFP in the main tracheal branches under a *btl-lexA* driver and strands 1-10 in ECs under the *Mex-Gal4* driver (Fig. 2k). While a constitutive red signal will allow visualisation of gut associated trachea (Fig. 2k, l), a GFP signal would only be detected upon direct contact between tracheal cells and ECs (Fig. 2k). While in homeostatic conditions we saw RFP but little GFP reconstitution, *Pe* infected midguts revealed increased tracheal branching and a significant increase in the number of GFP positive enterocytes within the epithelium (Fig. 2l, m). We observed no GFP signal in midguts independently expressing individual GRASP components (Fig. S2i). These results suggest the presence of physical interaction between ECs and gut associated trachea, which is induced during midgut regeneration. As such, gut-associated tracheal cells may be equipped to sense cellular mechanics of the midgut epithelium.

Cell morphology is largely under the control of two opposing mechanical forces, cell adhesion to adjacent cells and the ECM, and cortical tension, the force generated by the actomyosin cortex ^33, 50, 34^. Cortical MyosinII (MyoII) is an essential component of the cell cytoskeleton, which drives anisotropic cell tension and cell force generation ^51-56^. We used RNA interference to knock down the myosin light chain-c subunit of no-muscle MyoII from ECs of regenerating midguts, which restricted EC eccentricity, tracheal remodelling and ISC proliferation in regenerating midguts (Fig. 2n-r). Our results demonstrate a mechanical interplay between midgut enterocytes and gut associated tracheal cells, which contributes to adaptive tracheogenesis and ISC proliferation during midgut regeneration.

### Enterocyte apoptosis contributes to tissue mechanics and tracheal remodelling in the regenerating intestine

To address the origin of the biomechanical remodelling observed during midgut regeneration, we considered various cellular and physiological features, which are common to regenerating midguts exposed to *Pe* or DSS damage, including ISC proliferation (Supplementary Fig.1c, f), apoptotic cell death (Fig. 3a-b’), gut tube and visceral muscle resizing (Fig. 1c-f’ and Supplementary Fig. 2e, f), and gut peristalsis (Supplementary Fig. 2j). We noticed that only EC apoptosis recapitulated the R4/R5 regionalization we observed in intestinal mechanics (Fig. 3a-b’ and Supplementary Fig. 2a-f), with increased levels of apoptosis detected in R4 *versus* R5 upon *Pe* and DSS treatment (Fig. 3a-b’). Previously, we demonstrated that ISC proliferation was not a trigger for tracheal remodelling during midgut regeneration but rather lies downstream of it ^7^. We therefore hypothesised that EC cell death could contribute to the biomechanical changes observed in the regenerating midgut.

**Figure 3.**
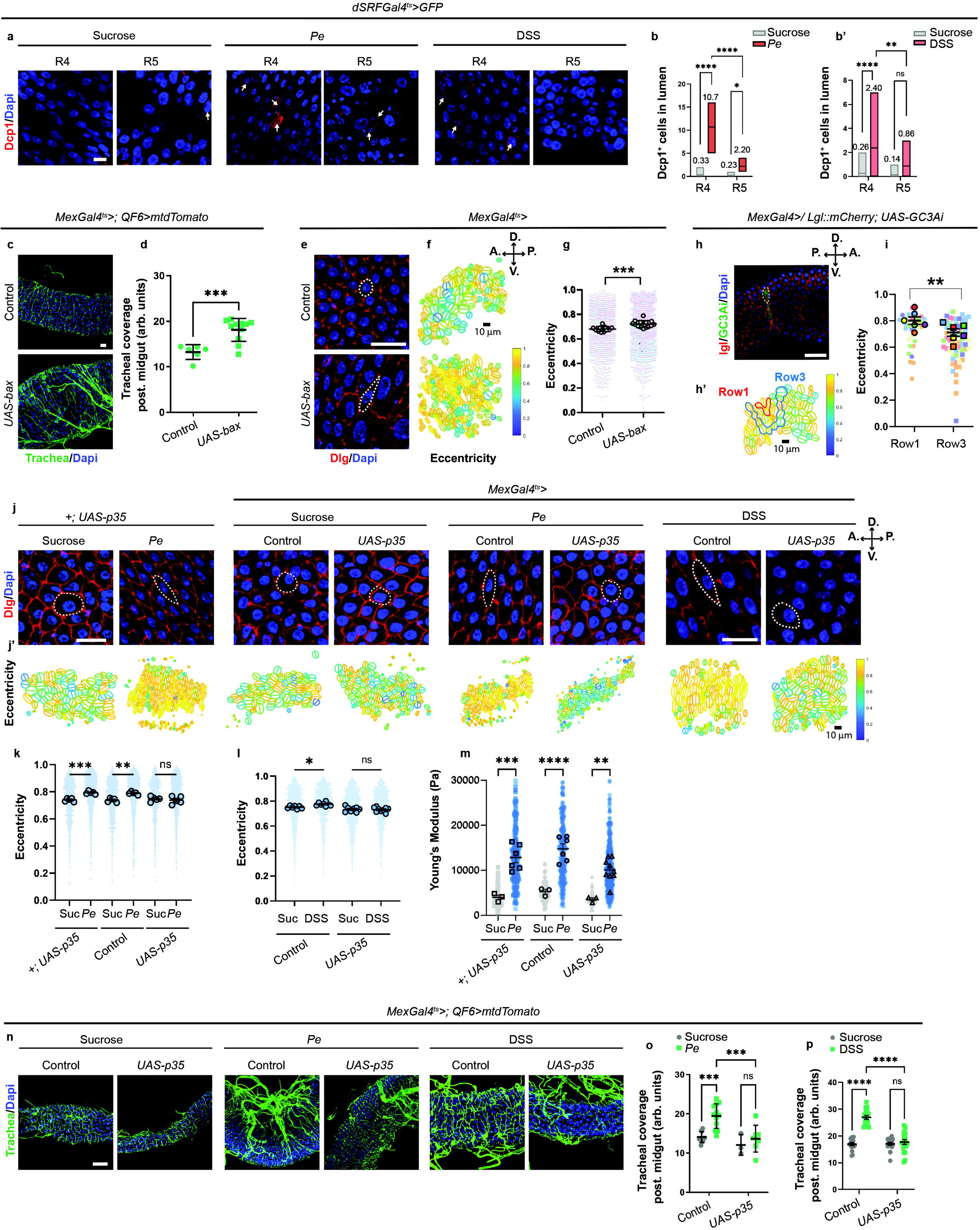
Enterocyte cell death contributes to cell geometry, tissue stiffness, and tracheal remodelling in the adult midgut. **a.** *Drosophila* caspase-1 (Dcp1) staining (white arrows) in R4/R5 regions under sucrose, *Pe* or DSS feeding. Scale bar, 20 μm. **b, b’**. Quantification of Dcp1+ cells in the posterior midgut lumen for conditions in a (n=18,10,17,10) (n=15, 15, 14, 15); box plots represent maxima, minima and mean. Mean value is indicated on top of boxes. **c**. Confocal images of midgut-associated trachea (red) in wildtype or enterocyte-specific *bax* expression (*Mex^ts^>bax*) midguts. Scale bar, 20 μm. **d**. Tracheal coverage quantification as in c (n=6,11). **e**. Enterocytes (Dlg, red) in the wildtype or *Mex^ts^>bax* midguts. Scale bar, 20 μm. **f**. Epithelial eccentricity as in e. **g**. Quantification of eccentricity as in f.; small circles indicate individual cells, large circles indicate per-gut averages, colours denote biological replicates (n=11,14). **h**. Midgut expressing apoptotic sensor GC3Ai (green) in enterocytes (*Mex-Gal4*) with cell boundary marker lgl (red), 2h post-infection; dashed line outlines an apoptotic cell. Scale bar, 50 μm. **h’.** Cellular eccentricity as in h. Row1 (red) and Row3 (blue) from the apoptotic cell highlighted. **i**. Quantification of eccentricity in Row1 and Row3 cells (n=6,6). **j**. Enterocyte septate junctions (Dlg; red) in wildtype or enterocyte-specific *p35* overexpression (*Mex^ts^>p35*) midguts under sucrose, *Pe* or DSS feeding. Scale bar, 20 μm. **j’.** Eccentricity as in j. **k, l**. Quantification of eccentricity, sucrose vs *Pe* (**k**) and Sucrose vs DSS (**l**) (n=5,5,5,5,5,5) (n=7,8,8,9). **m**. Stiffness (young’s modulus) via nanoindentation in midguts as in j.; small dots indicate individual measurements, large circles indicate per-gut averages (n=3,6,3,6,3,9). **n**. Trachea (red) in wildtype or *Mex^ts^>p35* midguts under sucrose, *Pe* or DSS feeding. Scale bar, 50 μm. **o, p**. Quantification of tracheal remodelling, sucrose vs *Pe* (**o**) and sucrose vs DSS (**p**) (n=9,12,3,7) (n=16,16,20,20). Data presented as mean ± SEM. n=numbers of guts analysed. DAPI labels nuclei (blue). <u>Statistics</u>: **b, d, g, i.** Two-tailed unpaired Student’s t-test. **k, l, m.** One-way ANOVA followed by Tukey’s multiple comparisons test. **o, p.** Two-way ANOVA followed by Tukey’s multiple comparisons test.

Consistent with our prior light microscopy-based results ^7^, induction of adult EC cell death by overexpressing the pro-apoptotic gene *bax* (*Mex^ts^>bax*) while fluorescently labelling trachea with *QF/QUAS* (*QF6>mtd-Tomato*) ^8^ showed that EC death in the absence of external damaging agents induces significant tracheal remodelling (Fig. 3c, d) and increases cellular eccentricity in the posterior midgut epithelium (Fig. 3e-g). A time course of oral *Pe* infection to monitor the earliest appearance of discrete cell death using the genetically encoded live apoptotic sensor *GC3Ai* ^57^ and *mCherry-lgl* revealed individual apoptotic ECs as early as 2hs after *Pe* feeding (Fig. 3h). Eliminating cells by apoptosis would locally result in a rosette (neighbouring cell stretching to compensate for the loss of an apoptotic cell) as reported previously ^58, 59^. This mechanism has been described as increasing neighbouring tissue tension ^60^. Consistently, image segmentation and morphometric analysis of midgut areas with individual apoptotic ECs revealed a localized pocket of epithelia with increased cellular eccentricity restricted to the area immediately surrounding an individual apoptotic cell (Fig. 3h-i). Furthermore, suppressing EC apoptosis by overexpressing the apoptotic inhibitor, p35 (*Mex^ts^>p35*) (Supplementary Fig. 3a-c) reduced the effect of midgut damage on cellular eccentricity (Fig 3j-l), gut stiffness (Fig. 3m) and tracheal remodelling (Fig. 3n-p) when compared to control genotypes. However, diminished EC apoptosis did not affect ROS levels in *Pe* infected midguts (Supplementary Fig. 3d, e), suggesting potentially independent roles of oxidative stress and EC cell death in the system.

Together, these findings identify EC apoptosis as a contributor to epithelial biomechanical changes and tracheal remodelling during regeneration of the *Drosophila* midgut.

### Compartmentalised biomechanical states in the regenerating mammalian intestine

Atomic force microscopy (AFM) assessment of snap-frozen cross-sections of the adult mouse small intestine showed increased stiffness in the apico-basal direction across the intestinal epithelium and its mesenchymal microenvironment ^61^ (Fig. 4a, b). Whole-body irradiation in mice involves the induction of intestinal cell apoptosis within the first 24hs (Supplementary Fig. 3f, g), followed by ISC proliferation and crypt regrowth 72hs post-injury ^62, 63^ (Fig. 4c-e and Supplementary Fig. 3f, h, h’). Results reported in our previous study indicated an increase in CD31+ endothelial cells surrounding regenerating crypts of the mouse small intestine ^7^. Immunostaining of E-CADHERIN junctions to segment the apical surface of crypt cells for morphometric analysis revealed increased cellular eccentricity of crypt cells from irradiated *versus* untreated mice (Fig. 4f-g). AFM of control and regenerating small intestines revealed that, while villi stiffness remained constant across treatments (Supplementary Fig. 3i), crypts undergoing regeneration following irradiation induced injury were stiffer than those from uninjured intestines (Fig. 4h). We noted that increased crypt stiffness was more prominent in regenerating intestines from female mice. Consistently, refined assessment of the blood vasculature through whole mount tissue preparations and immune-detection of PECAM-1, a marker of blood vascular endothelial cells, showed increased branching of blood vasculature, restricted to the crypt of the regenerating mouse small intestine (Fig. 4i, i’). Therefore, as in the *Drosophila* midgut, localised changes in epithelial geometry and tissue stiffness correlate with active remodelling of the vasculature surrounding regenerating crypts of the mouse small intestine.

**Figure 4.**
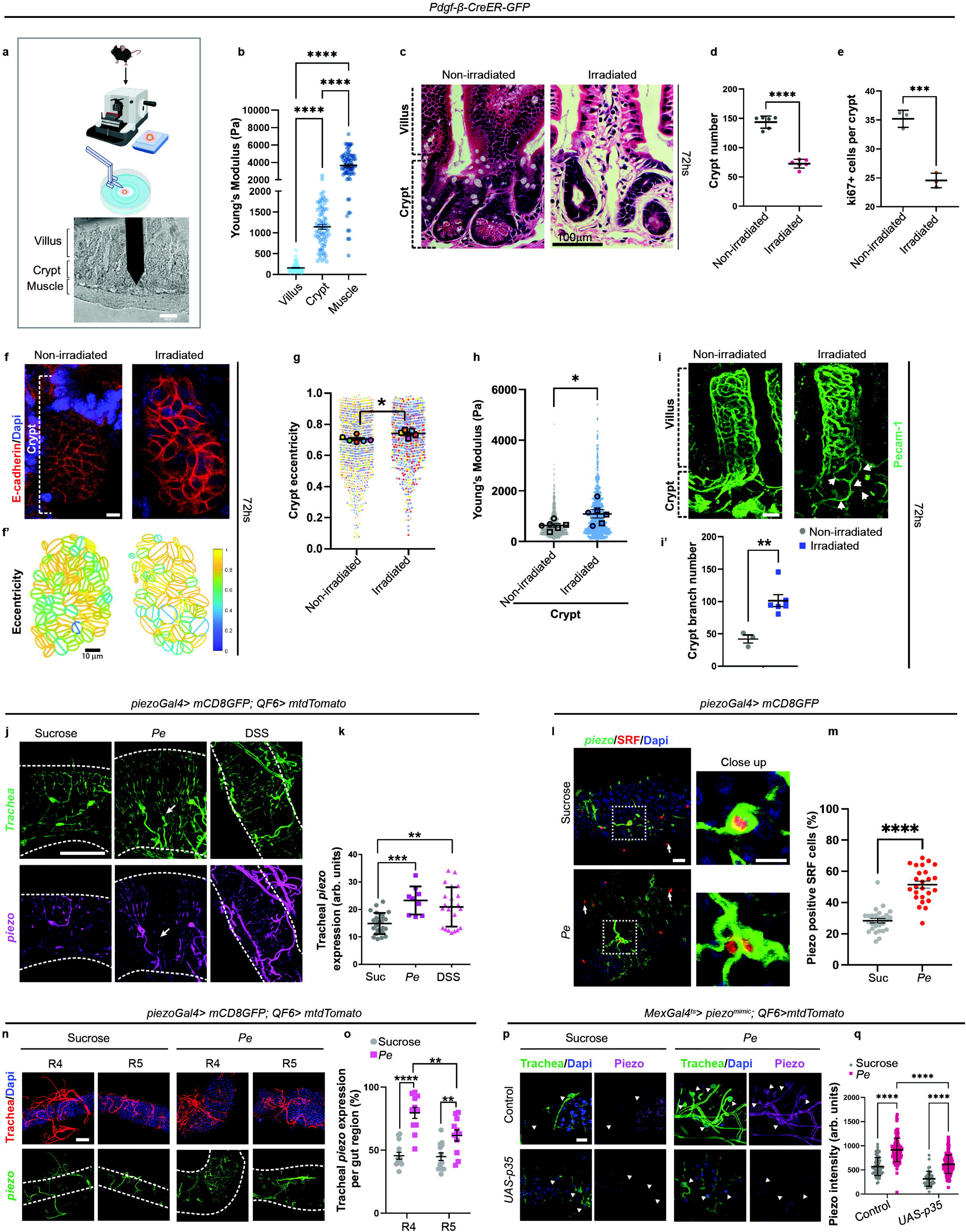
Compartmentalized cell geometry, mechanical changes and vascular remodelling in the regenerating adult mouse intestine. **a.** Nanoindentation setup. *Created in BioRender. Cordero, J. (2026)* https://BioRender.com/pr2zt7m. **b**. Stiffness (Young’s modulus) in small intestinal muscle, crypt and villus (n=1,1,1). **c**. H&E staining of small intestines from healthy and irradiated mice 72h post treatment. Scale bar, 100 mm. **d, e**. Quantification of (**d**) crypt numbers, and (**e**) Ki67^+ve^ cells per crypts (n=6,6) (n=3,3). **f**. Wholemount confocal images of intestinal crypts from control and irradiated mice; E-CADHERIN stains apical cell boundaries (red); DAPI (blue). Scale bar, 20 mm. **f’, g** Epithelial eccentricity; smaller circles indicate individual cell values and large circles indicate average cell eccentricity per gut (n=6,5). **h**. Stiffness in crypts of healthy and irradiated mice; Smaller dots represent individual cells, larger circles represent average values per gut (n=6,6). **i**. Wholemount immunostaining of crypt blood vasculature (PECAM-1; green) in homeostasis and post-irradiation. Arrows indicate expanded vasculature. Scale bar, 20 mm. **i’**. Quantification of crypt vasculature branches (n=3,6). **j**. Midgut trachea (green) *piezo* expression (magenta) upon Sucrose, Pe or DSS treatment. Arrows point to *piezo* negative trachea. Scale bar, 20 mm. **k**. Quantification of tracheal *piezo* expression (n=24,9,22). **l**. *piezo* expression (green) in midgut tracheal cell bodies (anti-SRF; red) upon sucrose vs *Pe* feeding. Scale bar, 50 mm. **m**. Percentage *piezo* expressing SRF cells in posterior midguts (n=27,25). **n**. Piezo expression (green) in the trachea (red) of R4/R5 regions following sucrose or *Pe* feeding. Scale bar, 50 mm. **o**. Quantification of R4/R5 tracheal *piezo* expression (n=13,11,13,11). **p**. Tracheal (green) Piezo expression (pink) in wildtype and (*Mex^ts^>p35*) midguts upon sucrose and *Pe* feeding. Arrowheads indicate tracheal cells. Scale bar, 20 mm. **q**. Quantification of tracheal expressed Piezo (n=70, 126, 41, 164); n=number of tracheal cells. Data represent the mean ± SEM. Unless otherwise noted, n= numbers of guts. <u>Statistics</u>: **d, e, g, h, i’, k, m.** Two-tailed unpaired Student’s t-test. **b**. One-way ANOVA. **o, q**. Two-way ANOVA followed by Tukey’s multiple comparisons test.

### Biomechanical changes within the *Drosophila* intestine are sensed by tracheal expressed mechanosensory channel Piezo

We next studied the mechanisms underpinning vascular remodelling downstream of intestinal mechanics. We hypothesised that biomechanical changes in the regenerating midgut epithelium were sensed through mechanosensory channels expressed in the gut vasculature.

Stimulation of vascular endothelial cells by intrinsic shear stress and stiffness drive angiogenesis and vascular remodelling ^64-66^. Mechanical signals in the vascular endothelium are largely transduced through TRP and Piezo family ion channels ^67, 68^. While the role of Piezo1 in vascular development is well documented ^69^, little is known about mechanisms of regulation and/or function of vascular Piezo1 in adult self-renewing epithelia. Recent studies in adult *Drosophila* and mouse intestine reported a role of stem/progenitor cell expressed *Piezo* in homeostatic stem cell maintenance and differentiation of the secretory lineage of intestinal stem cells ^43, 61^. While a single *piezo* gene exists in *Drosophila* ^70^, described functions of *Piezo* within the mammalian intestinal epithelium, are redundant between *Piezo1* and *Piezo2* ^61^.

Using an available *GAL4* reporter of endogenous *piezo* gene expression *(piezo-GAL4)* ^43^ we identified subpopulations of tracheal cells expressing *piezo* in the adult posterior midgut, representing approximately 25% of TTCs (Fig.4j-m). Strikingly, we observed a significant increase in tracheal *piezo* expression upon intestinal damage with orally administered *Pe* or DSS (Fig. 4j, k) and a two-fold increase in the overall proportion of *piezo* expressing TTCs in posterior midguts following *Pe* treatment (Fig. 4l, m and Supplementary Fig. 4a). In line with our findings of biomechanical regionalization in the posterior midgut (Fig. 2), *piezo* was most upregulated in R4 *versus* R5 (Fig. 4n, o). Tracheal *piezo* upregulation was confirmed with a reporter of Piezo protein expression (*piezo^mimic^*) (Supplementary Fig. 4b, c) ^71,^ _72._

Consistent with previous reports ^7, 8^, general tracheal remodelling of *Pe* treated midguts was reduced upon neutralization of ROS with NAC treatment (Fig. S4d-f). However, *piezo* upregulation within trachea of *Pe* treated midguts was not dependent on oxidative stress (Supplementary Fig. 4d, e). Nevertheless, we noted that *piezo* expressing TTCs showed partially reduced remodelling following NAC treatment (Supplementary Fig. 4d), suggesting that at least some of the mechanisms leading to tracheal remodelling downstream of *piezo* may be shared with those activated by oxidative stress ^7, 8^. Methylcellulose induced stretching of homeostatic intestines recapitulated *piezo* upregulation in TTCs (Supplementary Fig. 4g, h). Overall, these results suggest that mechanical changes within the regenerating *Drosophila* midgut epithelium induce non-autonomous upregulation of *piezo* in a subpopulation of gut associated trachea. Importantly, inhibition of EC apoptosis impaired Piezo upregulation in the trachea (Fig. 4p, q).

### Dynamic regulation of Piezo expression and activity during damage induced midgut regeneration

To further characterize *piezo* expression and activity in the midgut, we performed a time course experiment, which revealed a gradual increase in gut-associated tracheal *piezo* expression during a 16h period of *Pe* feeding (Fig. 5a, b). Time-dependent *piezo* expression correlated with gut tracheal remodelling dynamics and nanoindentation measurements of intestinal tube stiffness (Fig. 5c, d).

**Figure 5.**
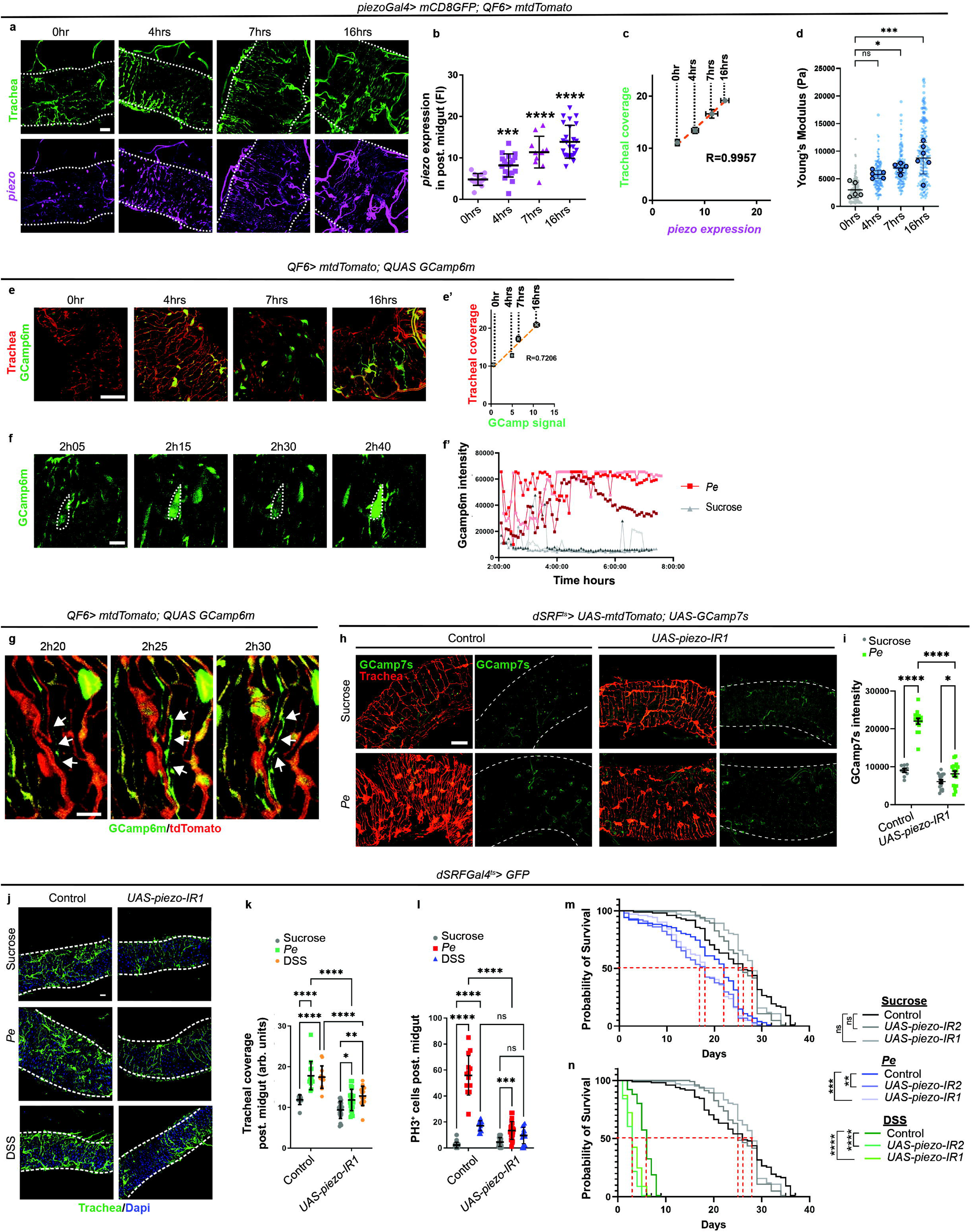
Dynamics of tracheal Piezo activity during *Drosophila* midgut damage. **a.** Confocal images of midgut trachea (green) and *piezo* expression (magenta) across 0-16h *Pe* infection. Scale bar, 20 mm. **b**. Quantification of tracheal *piezo* expression as in a (n=14,19,11,25). **c**. Tracheal coverage and *piezo* expression correlation across 0-16h *Pe* infection; Pearson’s correlation coefficient (r=0.9957) (n=14,19,11,25). **d**. Stiffness (Young’s modulus) in midguts across 0-16h *Pe* infection (n=5,5,5,6). **e**. Trachea (red) expressing GCamp6m calcium sensor (green) across 0-16h *Pe* infection. Scale bar, 20 mm. **e’**. Tracheal coverage and GCamp6m signal correlation across 0-16h *Pe* infection; Pearson’s correlation coefficient (r=0.7206) (n=19,20,16,20). **f**. Close up still views of tracheal GCamp6m from live imaging during *Pe* infection; cell bodies outlined. Scale bar, 20 mm. **f’**. Single tracheal cell GCamp6m intensity plotted across time course of *Pe* infection (red) or sucrose feeding (grey). Each line represents a single cell (n=3,3) n= tracheal cells. **g**. Close still views of tracheal (red) GCamp6m (green) from live imaging during *Pe* infection. Arrows indicate branches showing calcium pulse prior to a remodelling event. Scale bar, 20 mm. **h**. GCamp7s (green) in the trachea (red) following sucrose or *Pe* feeding in wildtype guts or upon tracheal *piezo* knockdown by RNAi (*piezo-IR1).* Scale bar, 20 mm. **i**. Quantification of tracheal GCamp7s intensity as in h (n=9,15,17,17). **j**. Confocal images of the midgut associated trachea (green) upon sucrose, *Pe* or DSS feeding in control animals or following tracheal *piezo* knockdown (*piezo-IR1)*. Scale bar, 20 mm. **k, l**. Quantification of tracheal coverage (**k**) and ISC proliferation (**l**) in midguts as in j (n=29,14, 13, 27, 23, 15) (n=29,14, 13, 27, 23, 15). **m, n**. Lifespan of wildtype and *piezo-IR* flies following sucrose (grey), *Pe* (blue) (**m**), or 6% DSS (green) (**n**) feeding (n=100,100,100). Red dashed lines indicate 50% survival per condition. Data represent the mean ± SEM. Unless otherwise noted n=number of guts or flies. <u>Statistics</u>: **m, n**. Mantel-Cox test. **d. One-way ANOVA** followed by Tukey’s multiple comparisons test. **b, i, k, l.** Two-way ANOVA followed by Tukey’s multiple comparisons test.

Signalling downstream of Piezo channel activation involves increased intracellular calcium (Ca^2+^) influx ^68, 69^. We therefore used the genetic Ca^2+^ sensor (*GCamp6m*) expressed in tracheal cells to measure tracheal Ca^2+^ during a time course of intestinal damage (Fig. 5e, f). Global and cell specific Ca^2+^ levels in TTCs increased over the time course of *Pe* infection (Fig. 5e-f’ and Movies 1, 2), with kinetics highly correlating to tracheal remodelling (Fig. 5e’). *In vivo* imaging of *GCamp6m* activity in individual TTCs revealed discrete Ca^2+^ pulses within extending and fusing cytoplasmic projections of TTCs as early as 2hs after the start of *Pe* feeding (Fig. 5g), which precedes a detectable increase in global tracheal remodelling (Fig. 5a, b). RNA interference to knock down *piezo* expression in adult TTCs (*dSRF^ts^>piezo-IR*) showed that increased Ca^2+^ signals within tracheal cells of the regenerating midgut were strongly dependent on *piezo* (Fig. 5h, i and Movies 2, 3).

In summary, these results suggest that temporal patterns of *piezo* gene upregulation and Piezo channel activation in gut associated trachea overlap with the dynamic TTC remodelling and gut biomechanical changes observed during midgut regeneration. Our *in vivo* tracking of Ca^2+^ influx in tracheal cells suggest that Piezo activation in gut associated trachea precedes measurable increase in tracheal remodelling.

### Tracheal Piezo contributes to TTC remodelling and regenerative ISC proliferation upon damage of the *Drosophila* midgut

Next, we used RNA interference gene knockdown to assess the functional role of tracheal *piezo* in TTC remodelling and ISC proliferation in regenerating midguts. To account for potential off target effects of the RNAis, we used two independent, priorly validated *piezo* RNAi lines ^43, 73-75^ that target different sections of the *piezo* gene and efficiently downregulate *piezo* mRNA (Supplementary Fig. 4i). Our results showed that, knocking down *piezo* in adult TTCs (*dSRF^ts^>piezo-IR*), diminished tracheal remodelling and ISC proliferation upon midgut injury by oral administration of *Pe* or DSS (Fig. 5j-l and Supplementary Fig. 4j, k). Of note, epithelial eccentricity was not affected in *dSRF^ts^>piezo-IR* midguts (Supplementary Fig. 4l, m), suggesting that mechanical changes in the midgut epithelium are upstream of tracheal *piezo* activation and function in midgut regeneration.

Intestinal damage compromises animal health and lifespan ^22^. Robust midgut regeneration is directly associated with the ability to maintain organismal survival in the face of intestinal epithelial damage ^76^. While showing no impact on homeostatic lifespan, tracheal *piezo* knockdown strongly reduced animal survival upon *Pe* and DSS feeding (Fig. 5m, n).

Finally, overexpression of tracheal *piezo* upon infection significantly potentiated tracheal remodelling and ISC proliferation during midgut regeneration and slowed down the recovery of tracheal and ISC homeostasis following removal of damage (Supplementary Fig. 4o-q). Altogether, these results suggest that tracheal *piezo* functionally contributes to TTCs remodelling, regenerative ISC proliferation and organismal survival upon *Drosophila* midgut damage.

### The blood vasculature and vascular *Piezo1* are functional components of the regenerative stem cell niche in the mammalian small intestine

The rich microenvironment of the mammalian gastrointestinal tract is composed of diverse cellular subtypes, including multiple mesenchymal cell populations, enteric neurons, and endothelial cells that together provide physical support, contractility, and multiple stem cell niche factors to the adult intestinal epithelium ^77-83^. While lymphatic vessels are known as functional components of the mammalian stem cell niche that supports intestinal stem cell function during tissue homeostasis and regeneration ^77, 84^, the role of the intestinal blood vasculature remains unknown.

Single molecule RNA in situ hybridisation detected co-expression of *Piezo1* and vascular *Pecam-1* in the mouse small intestinal crypt (Fig. 6a). We next used a tamoxifen inducible *Pdgf-*β*-Cre*-*GFP* ^85^ for visualisation of the intestinal blood vasculature from whole mount tissue preparations (Fig. 6b) ^86^ and to investigate the role of the blood vessels and vascular *Piezo1* in the adult mouse intestine from control *Pdgf-*β*-Cre*-*GFP* mice (*Piezo1^wt/wt^*) or *Pdgf-*β*-Cre*-*GFP* mice combined with an inducible *Piezo1* knockout allele (*Piezo1^fl/fl^*). In the first instance, adult mice were treated with orally gavaged tamoxifen for Cre induction and allowed to age for 5 days prior to intestinal damage by 10 Gy whole body irradiation (IR), followed by small intestine tissue sampling at 72 hours post IR (Supplementary Fig. 5a). Knocking out *Piezo1* from the adult vasculature had no apparent impact on overall animal health, general intestinal vasculature structure (Supplementary Fig. 5b-d) or self-renewal of the intestinal epithelium in homeostatic conditions (Supplementary Fig. 5e-i). On the other hand, vascular *Piezo1* knock out animals showed a significant impairment in crypt vascular remodelling (Fig. 6c, d), and a decrease in the total number of regenerating crypts and proliferating cells within intestinal crypts, following irradiation-induced damage (Fig. 6e-h).

**Figure 6.**
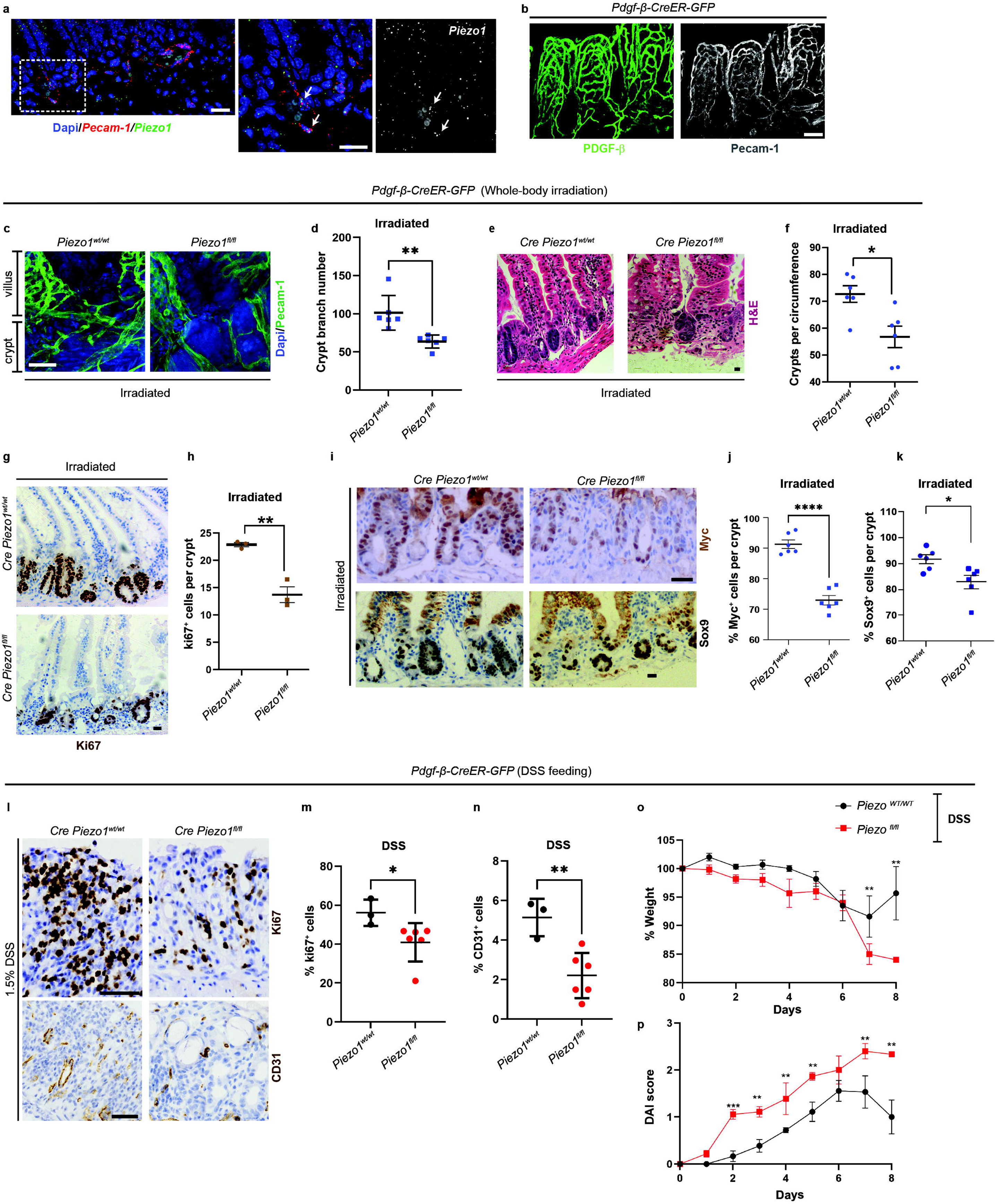
The role of vascular *Piezo1* is conserved in mouse models of intestinal regeneration. **a.** Confocal images of single molecule RNA in situ detection of *Pecam-1* (red) and *Piezo-1* (green; grey) in mouse small intestinal crypts. DAPI (blue). Right panel is a magnified view of the area highlighted by the dotted box in the left panel (n=1). Scale bar, 20 mm. **b**. Wholemount immunostaining of the crypt intestinal vasculature of *Pdgf-beta-CreER (*green) co-stained with an anti-PECAM-1 antibody (grey) (n=1). Scale bar, 50 mm. **c**. Wholemount immunostaining of the crypt intestinal vasculature of irradiated *Pdgf-beta-CreER Piezo1^wt/wt^* and *Piezo1^fl/fl^* mice. Scale bar, 10 mm. **d**. Quantification of crypt vascular branch number in intestines as in b (n=6,6). **e**. H&E staining of small intestinal samples from irradiated *Pdgf-beta-CreER Piezo1^wt/wt^* and *Piezo1^fl/fl^* mice. Scale bar, 20 mm. **f**. Quantification of the number of crypts per circumference as in e (n=6,6). **g**. Ki67 staining of staining of small intestinal samples from irradiated *Pdgf-beta-CreER Piezo1^wt/wt^* and *Piezo1^fl/fl^* mice. Scale bar, 20 mm. **h**. Quantification of the number of Ki67+ cells per crypt as in g (n=3,3). **i**. Immunostaining of Myc and Sox9 in intestinal crypts of *Pdgf-beta-CreER Piezo1^wt/wt^* and *Piezo1^fl/fl^* mice following irradiation. Scale bar, 20 mm. **j, k.** Quantification of Myc (**j**) and Sox9 (**k**) (n=6,6) (n=6,6). **l**. Ki67 and CD31 staining of colon samples from DSS treated *Pdgf-beta-CreER Piezo1^wt/wt^* and *Piezo1^fl/fl^* mice. Scale bar, 20 mm. **m, n**. Quantification of percentage of ki67+ (**m**) and CD31+ (**n**) cells in ulcerated regions of the colon (n=3,6) (n=3,6). **o**. Percentage weight loss of *Pdgf-beta-CreER Piezo1^wt/wt^* (black) and *Piezo1^fl/fl^* (red) mice following onset of DSS treatment (n=6,6). **p**. DAI score of *Pdgf-beta-CreER Piezo1^wt/wt^* (black) and *Piezo1^fl/fl^* (red) following onset of DSS treatment (n=6,6). Data represent the mean ± SEM. n=numbers of guts/mice analysed. <u>Statistics</u>: Two-tailed unpaired Student’s t-test.

Wnt signalling is a central regulator of intestinal health and disease in mammals ^87^. Wnt signalling activity as per levels of MYC and SOX9 ^88, 89^ in post-damage regenerating crypts was impaired in *Piezo1^fl/fl^* mice compared to *Piezo1^wt/wt^* counterparts (Fig. 6i-k).

To further interrogate the functional implications of vascular *Piezo1* in intestinal and organismal health, we tested its impact in the large intestine, following inflammatory damage induced by DSS treatment ^90, 91^ (Supplementary Fig. 6a, b). Our analysis of intestinal histology and overall animal health in *Pdgf-*β*-Cre*-*GFP* control and *Piezo1^fl/fl^* mice, which was carried out 3 days after the removal of DSS (Supplementary Fig. 6a, b), showed *Piezo1^fl/fl^* mice exhibiting significantly impaired cell proliferation (Fig. 6l, m) and vascularization (Fig. 6l, n) of the large intestine, accompanied by accelerated body weight loss (Fig. 6o) and higher disease associated index (DAI) scores, which are characterised by the presence of weight loss, diarrhoea and rectal bleeding (Fig. 6p). As in the case of the irradiation, functional effects of vascular Piezo1 were evident only upon DSS induced damage (Supplementary Fig. 6b-e)

Therefore, endothelial Piezo1 helps to promote angiogenesis and regenerative proliferation of the mouse intestine upon irradiation and DSS induced damage, and to protect organismal health from the deleterious effects of intestinal inflammation.

### Tracheal Piezo activates a Yorkie/YAP-TAZ dependent mechanosensory program for tracheal remodelling and ISC proliferation in the regenerating midgut

Intriguingly, our results showed that Piezo channel activity and gene expression are both upregulated in the trachea during midgut regeneration (Fig.5). We next used the chemical agonist of Piezo, Yoda1, in a concentration previously shown to effectively and specifically activate Piezo in *Drosophila* ^92^ and mice ^93^, upon ectopic or systemic application, respectively. Feeding animals with 1mM Yoda1 was sufficient to induce upregulation of *piezo* expression within the trachea (Fig. 7a, b), and recapitulated *piezo* gain of function phenotypes of increased tracheal remodelling (Supplementary Fig. 4o-q) and mild ISC proliferation (Fig. 7c, d) ^43^. Therefore, Piezo activation appears involved in *piezo* gene expression within adult *Drosophila* midgut trachea. We next investigated the mechanisms leading to *piezo* upregulation in the trachea and downstream mediators of the role of Piezo in midgut tracheal remodelling.

**Figure 7.**
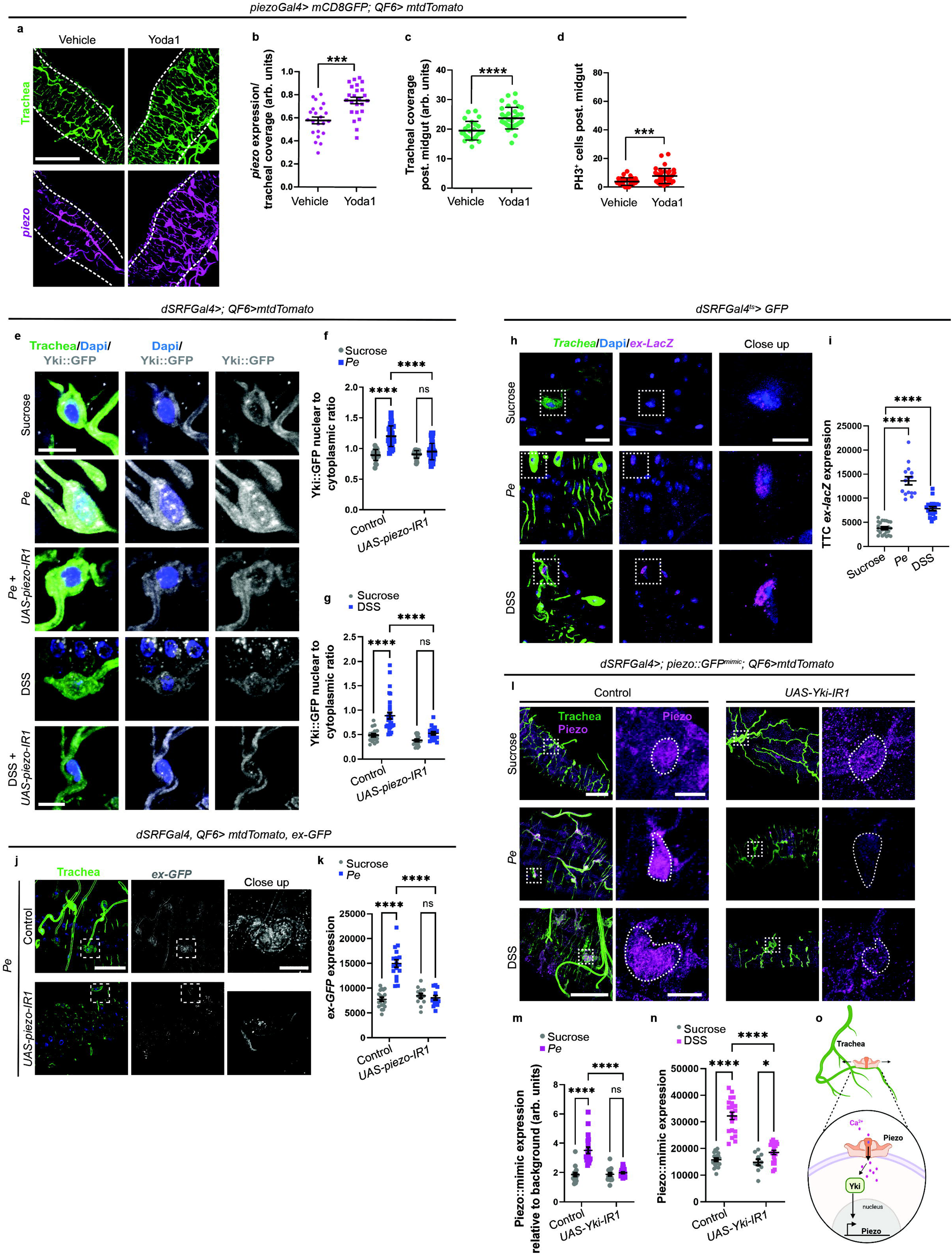
Piezo contributes to Yki activity and remodelling pathways in gut associated trachea. **a.** Midgut trachea (green) and *piezo* expression (magenta) following Yoda1 treatment. Scale bar, 100 mm. **b**. Quantification of tracheal *piezo* expression in midguts as in a (n=22,24); n= gut numbers. **c, d.** Quantification of tracheal coverage (**c**) and ISC proliferation (**d**) in midguts as in a (n=25,35) (n=31,35); n= gut numbers. **e**. Midgut trachea (green) and Yki::GFP (grey) upon sucrose, *Pe* or DSS feeding in control or following tracheal *piezo* knockdown (*piezo-IR1*). Scale bar, 10 mm. **f**, **g**. Quantification of tracheal Yki nuclear to cytoplasmic ratio in midguts as in e. (n=23,30,28,41) (n=19,31,20,18); n= tracheal cells. **h**. Expanded expression (*ex-lacZ*; pink) in midgut trachea (green) upon sucrose, *Pe* or DSS feeding. Scale bar, 10 mm. **i**. Quantification of tracheal *ex-lacZ* expression as in h (n=41,15,19). **j**. of Ex-GFP (grey) expression in midgut trachea (green) upon sucrose or *Pe* feeding in control midguts or following tracheal *piezo* knockdown (*piezo-IR1).* Scale bars, 20 mm and 10 mm. **k**. Quantification of tracheal Ex-GFP expression as in j (n=19,17,13,15). **l**. Confocal images of midgut trachea (green) and Piezo expression (pink) upon sucrose, *Pe* or DSS feeding in wildtype flies or upon tracheal *Yki* knockdown (*Yki-IR1*). Dashed lines highlight individual cell bodies. Scale bars, 50 mm and 10 mm. **m, n**. Quantification of tracheal Piezo expression in midguts as in l (n=17,20,20,20) (n=20,21,10,21). **o**. Schematic overview of pathways downstream of tracheal Piezo. *Created in BioRender. Cordero, J. (2026)* https://BioRender.com/ikh3fao. Data represent the mean ± SEM. Unless otherwise noted, n=number of guts<u>. Statistics</u>: **b, c, d, i**. Two-tailed unpaired Student’s t-test. **f, g, k, m, n**. Two-way ANOVA followed by Tukey’s multiple comparisons test.

The mechanosensitive transcription factors YAP/TAZ have been functionally associated to *Piezo1* in mammalian endothelial cells ^94-96^. Therefore, their *Drosophila* orthologue, Yki, seemed a suitable candidate to link Piezo activity with the regulation of its own gene expression in the trachea. As with its mammalian counterparts, Yki activation leads to its translocation from the cytoplasm to the nucleus to drive transcriptional target regulation ^97^. Using an endogenous Yki protein reporter line, we monitored Yki localisation within gut associated trachea in homeostatic and regenerating midguts. While Yki expression was largely restricted to the cytoplasm of TTCs in homeostatic midguts (Fig.7e; top panels), pathogenic infection or DSS feeding led to translocation of Yki into TTCs’ nuclei, which was prevented by *piezo* knockdown (Fig. 7e-g). The Hippo signalling target and Yki partner *expanded* (*ex*) ^98^ was also upregulated in TTCs from regenerating midguts (Fig. 7h, i). Importantly *ex* upregulation in the trachea was not only dependent on Yki (Supplementary Fig. 7a, b), but it was also reliant on Piezo expression (Fig. 7j, k and Supplementary Fig. 7c). Furthermore, activation of Piezo by Yoda1 induced nuclear translocation of Yki and upregulation of *ex* in TTCs (Supplementary Fig. 7d-g). Importantly, *yki* knockdown inhibited Piezo upregulation in TTCs of *Pe* and DSS treated midguts (Fig. 7l-n).

Collectively, these results support the existence of a feedback mechanism through which Yki and Piezo positively regulate each other in TTCs during midgut regeneration; Yki is activated in remodelling trachea from the regenerating midgut in a Piezo dependent manner, and activation of Yki in turn contributes to the upregulation of tracheal *piezo* expression (Fig. 7o).

Subsequently, we further investigated the functional role of tracheal Yki in TTC remodelling and regenerative ISC proliferation in the adult midgut, and its connection with the role of Piezo in the system. Firstly, we knocked down *yki* from the adult trachea using two independent RNAi lines, which resulted in a significant decrease in TTC remodelling and ISC proliferation in *Pe* and DSS treated midguts (Fig. 8a-c and Supplementary Fig. 7h, i). Conversely, overexpression of a constitutively active Yki (*Yki^CA^*) in adult TTCs ameliorated the negative impact of *piezo* knockdown on tracheal remodelling and ISC proliferation in damaged midguts (Fig. 8d-f). Furthermore, tracheal *Yki^CA^* overexpression in wild type midguts potentiated tracheal remodelling during the damage phase of intestinal regeneration (Supplementary Fig. 8a-b’), and delayed homeostasis recovery of TTC remodelling and ISC proliferation ^7^ 48hs after damage removal (Supplementary Fig. 8b’-c’).

**Figure 8.**
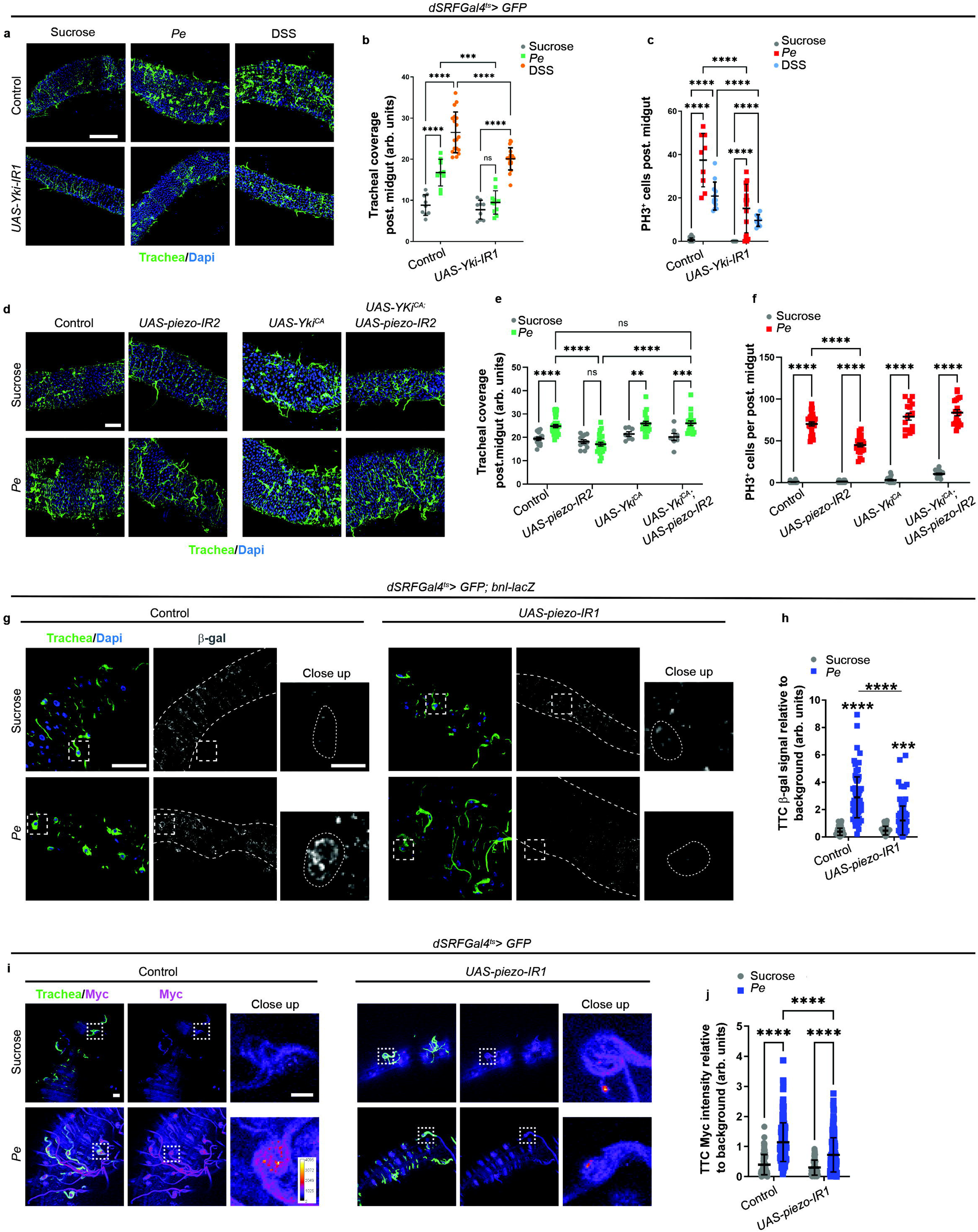
Yki influences tracheal remodelling downstream of Piezo. **a.** Midgut trachea (green) upon sucrose, *Pe* or DSS feeding in control midguts or following tracheal *yki* knockdown (*yki-IR1*). Scale bar, 20 mm. **b, c**. Quantification of tracheal coverage (**b**) and ISC proliferation (**c**) in midguts as in a (n=9,9,18,9,11,18) (n=21,9,12,10,16,8). **d**. Confocal images of midgut associated trachea (green) without (control) or with overexpression of constitutively active *Yki* (*Yki-CA*) and *piezo* knockdown upon *Pe* infection. Scale bar, 50 mm. **e, f.** Quantification of tracheal coverage (**e**) and ISC proliferation (**f**) in midguts as in d (n=15,28,12,24,7,20,7,18) (n=24,27,21,22,16,16,15,18). **g**. Confocal images of midgut associated trachea (green) and anti-Bnl expression (grey) upon sucrose or *Pe* feeding in control midguts or following tracheal *piezo* knockdown (*piezo-IR1*). Scale bar, 50 mm, 10 mm. **h**. Quantification of Bnl expression per TTC in midguts as in g (n=89,95,44,111); n=tracheal cells. **i**. Representative confocal images of midgut associated trachea (green) and anti-Myc expression (pink) upon sucrose or *Pe* feeding in control midguts or following tracheal *piezo* knockdown (*piezo-IR1*). Scale bar, 10 mm, 10 mm. **j**. Quantification of Myc expression per TTC in midguts as in k (n=106,196,61,225); n=tracheal cells. Data represent the mean ± SEM. Unless otherwise noted, n= numbers of guts analysed. Dashed lined squares or circles highlight individual tracheal cells. <u>Statistics</u>: Two-way ANOVA followed by Tukey’s multiple comparisons test.

Oxidative stress induces a tracheal programme, including upregulation of Fibroblast growth factor-like *branchless* (*bnl*), its receptor breathless (*btl*) and the transcription factor Myc in TTCs ^7, 8^. Interestingly, while *piezo* induced Bnl and Myc upregulation following oral exposure with *Pe* (Fig. 8g-j) and DSS (Supplementary Fig. 8d-g), it did not affect upregulation of *btl* (Supplementary Fig. 8h, i). These results suggest that mechanical and oxidative stress signals partially overlap in the molecular programs they regulate in TTCs during midgut regeneration. Consistent with the functional interrelationship between Piezo and Yki, tracheal *yki* knockdown impaired Bnl and Myc upregulation in TTCs from regenerating midguts (Fig. 9a-d and Supplementary Fig. 9a-d), albeit with non-significant effects on Myc upregulation upon DSS treatment (Supplementary Fig. 9c, d). Finally, blocking EC apoptosis in *Pe* treated midguts strongly impaired Myc upregulation in TTCs (Supplementary Fig. 9e, f), reaffirming the positive contribution of epithelial cell death towards activation of tracheal molecular programs during midgut regeneration. Overall, these results reveal a mechano-chemical model of gut/tracheal interaction during adult midgut regeneration where Piezo activation within gut associated trachea is translated into a cell intrinsic molecular program through the activation of Yki in terminal tracheal cells, which includes upregulation of tracheal *piezo* itself as well as Bnl/FGF and Myc, leading to regenerative tracheal remodelling and ISC proliferation (Fig. 9e).

**Figure 9.**
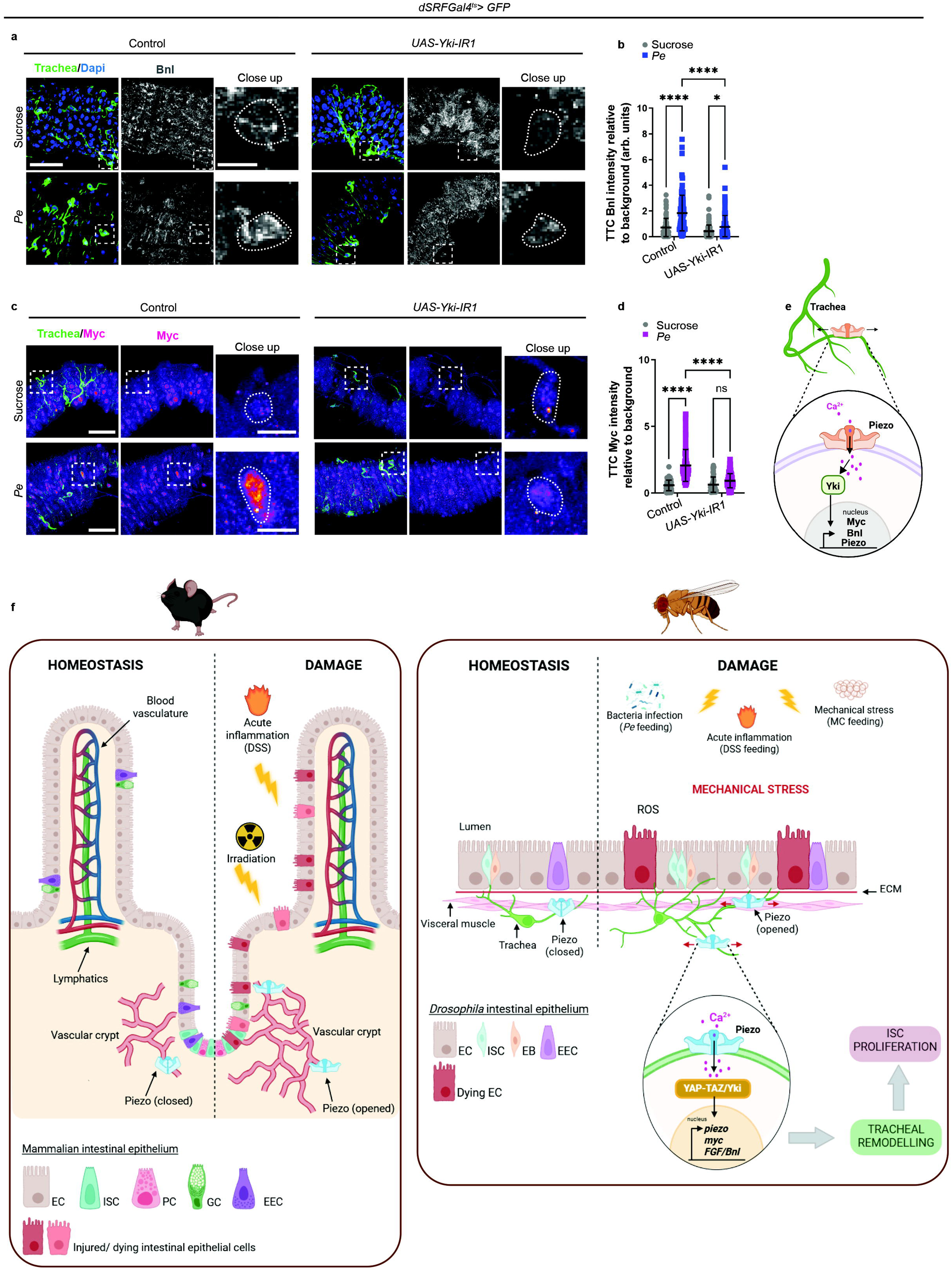
Yki activation is upstream of tracheal remodelling pathways. **a.** Representative confocal images of midgut trachea (green) and anti-Bnl expression (grey) upon sucrose or *Pe* feeding in control midguts or following tracheal *yki* knockdown (*yki-IR1*). Scale bar, 50 mm and 10 mm. **b**. Quantification of Bnl expression per TTC in midguts as in a (n=78,111,116,129); n=tracheal cells. **c**. Representative confocal images of midgut associated trachea (green) and anti-Myc expression (pink) upon sucrose or *Pe* feeding in control midguts or following tracheal *yki* knockdown (*yki-IR1*). Scale bar, 50 mm and 10 mm. **d**. Quantification of Myc expression per TTC in midguts as in c (n=66,103,78,122); n=tracheal cells. Dashed lined squares or circles highlight individual tracheal cells. **e**. Schematic overview of pathways downstream of Piezo and Yki activation in gut trachea. *Created in BioRender. Cordero, J. (2026)* https://BioRender.com/ikh3fao. **f**. Schematic representation of our working model for vascular and tracheal Piezo in intestinal regeneration. *Created in BioRender. Cordero, J. (2026)* https://BioRender.com/dx9ou7e. Data represent the mean ± SEM. <u>Statistics</u>: Two-way ANOVA followed by Tukey’s multiple comparisons test.

Altogether, our cross-species study reports a previously unrecognised pro-angiogenic mechanical crosstalk between the intestinal epithelium and the tracheal/vascular tissue. Apoptotic intestinal epithelial cells contribute to changes in cellular geometry and tissue reshaping, and mechanical stress within the regenerating intestinal epithelium, which induce activation of a subset of trachea/vascular cells expressing the mechanosensory channel Piezo. Piezo activation in the vascular microenvironment is part of the machinery driving remodelling of the vascular niche, regenerative stem cell proliferation and organismal health upon intestinal damage and inflammation (Fig. 9f).

## Discussion

The vasculature is an intrinsically plastic tissue with central metabolic and signalling roles that support developing and adult organs. A few of the many diverse contexts of vascular remodelling in physiology and disease include pregnancy and foetal growth, wound healing, bone fracture repair, tumour growth and metastasis ^13, 27, 99-103^. A seemingly common theme amongst the mentioned examples is the presence of vascular adaptation to environmental and/or host organ changes through largely unknown mechanism. Here, we report a cross-species mechanobiological crosstalk between the intestinal epithelium and its tracheal/vascular system, which drives reciprocal adaptation of the intestine and vascular microenvironment during intestinal regeneration.

### A diversity of extrinsic signals leading to common phenomenology of post-developmental vascular remodelling

Hypoxia and oxidative stress modulate vascular/tracheal remodelling across species ^104-107^. Nevertheless, adaptive vascularization of adult organs is observed in the context of a wide range of chemical, physiological and environmental cues ^7-13, 108^. Our previous observations showed a conserved phenomenology of tracheal/vasculature changes upon diverse intestinal insults ^7^. Here, we show that diverse forms of intestinal damage converge on a common programme of epithelial biomechanical remodelling, characterised by changes in tissue geometry, mechanical properties and mechanosensory signalling. As such, our findings may explain mechanisms of vascular remodelling observed in broad physiological and pathological contexts.

Our results revealed Piezo expression in 25%-50% of tracheal cells in the posterior midgut, which do not appear to depend on FGFR/Btl for their function and are largely refractory to the action of the antioxidant NAC. These results suggest that TTCs in the adult intestine may be highly heterogeneous. For instance, Piezo and Btl expressing trachea could represent two distinct tracheal sub-populations. Tracheal cell diversity may explain the partial impact of *piezo* knockdown on regenerative ISC proliferation (Fig. 5l) as well as the ability of the tracheal/vascular microenvironment to respond to different cues and support intestinal function accordingly.

Despite the heterogeneous nature of *piezo* expression in midgut trachea, gene knock down from TTCs results in rather global effect on tracheal tissue remodelling. A similar phenomenon is observed upon *btl* knockdown ^7, 8^. How do context specific cues sensed by a sub-set of tracheal cells translate into global vascular/tracheal responses? One possible answer to this open question may lie in the existence of paracrine interaction amongst distinct tracheal cell populations. While definite proof of such a phenomenon remains to be obtained, our live imaging-based visualization of intracellular Ca^2+^ signals in gut-associated trachea revealed the presence of branch fusion events between distinct tracheal cells and transfer of intracellular calcium, during TTC remodelling (Fig. 5g).

### Cellular apoptosis as a contributor to intestinal biomechanics and gut associated tracheal remodelling during intestinal regeneration

We show that enterocyte morphology and tissue mechanics change within the intestine across different challenges, including chemical, pathogenic damage and experimentally induced gut expansion (Fig. 1). Whilst the intestinal visceral muscle and the extracellular matrix may likely contribute to the biomechanical properties of the gut that are relayed to the trachea, intestinal epithelial changes appear to be important events underpinning the altered cellular geometry and tissue mechanical properties observed in regenerating midguts. We identified physical interaction between trachea and intestinal EC, and EC apoptosis as a cue contributing to mechanical changes in regenerating midguts as well as tracheal remodelling. Previous studies have characterized a biomechanical changes generated upon cell death in developmental and homeostatic contexts ^58, 59^, in a mechanism described as increasing neighbouring tissue tension ^60^. Furthermore, in a phenomenology reminiscent of our observations on individual apoptotic cells (Fig. 3h), cells closest to apoptotic cells experience tensile stretching in a model of homeostatic compensatory proliferation *in vitro* ^109^. Our findings suggest that proposed roles of apoptosis in tissue biomechanics may also apply to a complex, multilayer epithelial tissue such as intestine. However, apoptosis is likely a context dependent rather than universal driver of biomechanical organ/vascular interactions.

### Multicellular role of Piezo mechanosensing in the adult intestine

Pioneering work in *Drosophila* discovered a role of Piezo within stem/progenitor cells in the intestine in differentiation of the secretory lineage of ISCs ^43^ and extracellular matrix repair upon intestinal damage ^110^. Subsequent work in the mammalian intestine reported redundant roles of intestinal stem cell *Piezo1* and *Piezo2* in homeostatic stem cell maintenance and differentiation of secretory cell lineages in the mouse intestine ^61^.

Using *Drosophila* and mouse models, our results demonstrate a non-redundant role of vascular/tracheal *Piezo1* in intestinal regeneration, contributing to angiogenesis, ISC proliferation and organismal health, upon intestinal damage. *Piezo* expression in other gut associated tissues includes enteric neurons in both *Drosophila* and mammals, as well as in mammalian intestinal smooth muscle, with important mechanosensory functions leading to the regulation of gut peristalsis, inflammation and feeding ^56, 73-75, 111, 112^. Given the central contribution of enteric neurons and the visceral muscle to the intestinal stem cell niche ^113, 114^, and the observed role of tracheal Piezo in regenerative ISC proliferation, it would be very important to functionally interrogate the contribution of additional microenvironmental sources of Piezo to homeostasis and regeneration of the midgut epithelium.

## Methods

### Full genotypes and Resource Table

A list of full genotypes of animals used in this study can be found in Supplementary Table 1. A list of all resources used in this study can be found in Supplementary Table 2.

### Fly husbandry

For temperature sensitive experiments, using the *GAL4/UAS* or *LexA/LexAop; GAL80^ts^* system, flies were crossed, the F_1_ progeny were reared, and experimental adults were aged for 3-5 days after eclosion at 18°C. Then, animals were moved to 29°C for 5 days to allow transgenes activation. Experiments that did not contain temperature-sensitive transgenes were performed with animals kept at 25°C and experimental adults aged for 5 days prior to any procedure. All experimental animals were transferred to fresh food every 2 days, except for survival experiment where food was changed every day. Only posterior midguts from adult mated females were analysed in this study.

### *Drosophila* intestinal damage models

#### Bacterial infection

*Pseudomonas entomophila* (*Pe*) – females were fed on filter papers containing 400 µl of 5% sucrose solution only or sucrose containing *Pe* OD_600_=100 for 16hs or otherwise indicated.

#### DSS treatment

(Sigma-Aldrich, cat no. 42867) – females were fed on filter papers containing 400 µl of 5% sucrose solution only or sucrose containing 3% DSS for 2 days with fresh medium applied daily.

#### Methylcellulose feeding

(Sigma-Aldrich, cat no. 274429) – For our mainstream protocol, females were fed on normal food as a control or on 5% sucrose containing 10% methylcellulose for 72hs. Flies were transferred onto new food every two days. Prior to dissection, control flies were transferred overnight to 400 µl of 5% sucrose on filter papers to clear the gut. Flies exhibiting guts with over a 30% increase in diameter were selected as exhibiting the ‘full gut’ methylcellulose phenotype. Using this protocol, only about 10% of the flies experienced the over full phenotype. We therefore adapted the protocol by gluing the annus of the flies to prevent excretion of the MC. In this case, MC was fed for only 16hs, and this adaptation highly increased the efficiency of animals with MC in the gut, with 60-80% of flies fed the polymer exhibiting the overfull phenotype.

### *Drosophila* chemical treatments

#### NAC treatment

(Sigma-Aldrich, cat no. A7250) – Flies were placed on filter paper discs soaked with 400 µl of 5% sucrose solution containing 20mM NAC for 24hs. Then, flies were fed with either 5% sucrose + NAC or 5% sucrose + NAC + *Pe* (OD_600_=100) for 16hs or 5% sucrose + NAC + 3% DSS for 48hs ^7^.

#### DHE treatment

(Invitrogen Molecular Probes, cat no. D23107) – Tissues were dissected in Schneiders medium and then incubated in 30mM DHE solution for 5-7mins at room temperature and in a dark chamber. Then, midguts were washed three times with Schneiders medium for 5mins, mounted in DAPI containing media followed by immediate imaging and image capture.

#### Yoda1 treatment

Yoda1 (Tocris, cat no. 5586) was suspended in DMSO and diluted to 1 mM in 5% sucrose. Vehicle containing an equivalent volume of DMSO in 5% sucrose was used as control. Flies were placed in vials containing filter paper discs soaked with either solution for 16h at 29°C prior to dissection.

### *Drosophila* survival experiments

All experiments were done on 5 days old adult flies. Flies were exposed to bacteria (*Pe* OD_600_=100) for 24hs, or 6% DSS treatment for the entire course of the experiment. The control flies were exposed to 5% sucrose solution. Flies were kept at 29°C and monitored for survival post-treatment which was recorded daily. *Pe* treated flies were flipped into fresh food vials every day and DSS treated flies were kept on fresh 6% DSS solution every day. The survival curves were calculated from 5 vials of 20 flies per treatment group.

### *Drosophila* immunohistochemistry

Guts from adult female flies were dissected in PBS and immediately fixed in 4% formaldehyde (FA, EM grade, Polysciences) diluted in PBS for 1h at room temperature (RT) and then washed three times in PBS-T (PBS + 0.2% Triton X100). Samples were incubated with the primary antibody diluted in PBT (PBS-T + 2 % BSA) overnight at 4°C. Guts were washed three times in PBS-T and then incubated with secondary antibody in PBT for 2hs at RT. After incubation, guts were washed three times for 20mins in PBS-T and mounted on glass slides with 13mm x 0.12mm spacers (sigma) in Vectashield mounting medium with DAPI (Vector Laboratories). For anti-Myc, anti-β-gal, anti-Dcp1, *piezo::mimic* and anti-Bnl staining, fixation was performed in 4% formaldehyde for 1h, followed by 5min methanol fixation added to the sample gradually and then 10min in 100% methanol. Samples were blocked in 7% goat serum for 1h before primary antibody incubation. The following primary antibodies were used: chicken anti-GFP (1:200, Abcam, cat. no. ab13970), rabbit anti-DsRed (1:1000, Clontech, cat no. 632496), mouse anti-Dlg (1:50, DSHB 4F3E2, AB 528203), mouse anti-PH3 (1:100; Cell Signaling, cat. No. 9706), rabbit anti-cleaved Dcp1 (1:100; Cell Signaling, cat No.9578s), rabbit anti-β-gal (1:1,000; MP Biochemicals, cat no. 559761), guinea pig anti-Myc (1:100, gift from G. Morata), rat anti-Bnl (1:50, M. Krasnow), mouse anti-SRF (1:50, gift from M. Krasnow). Secondary antibodies: Alexa Fluor anti-chicken 488 (1:200), anti-rabbit 594 (1:100) and anti-mouse 647 (1:100) (Invitrogen) were used. Guts were mounted in Vectashield anti-fade mounting medium for fluorescence with DAPI (Vector Laboratories, Inc.) to visualize nuclei.

### Image acquisition

#### Confocal Microscopy

(Zeiss LSM 780/880) – Each image represents half-volume of the posterior midgut (area between hindgut and the copper cell region) and was acquired with a ×20, ×40 or ×63 lens on a Zeiss LSM 780 or 880 confocal microscope using identical acquisition conditions for all samples from a given experiment. The images represent the maximal intensity projection of Z-stacks and were processed using ImageJ (Version 1.54f) and Carl Zeiss Zen 3.0 to adjust the brightness and contrast^7^.

#### Nanoindentation

Flies were anesthetised on ice and midguts were freshly dissected on the day of measurement in Adult Hemolymph-like culture media (2mM CaCl2, 5mM KCl, 5mM HEPES, 8.2mM MgCl2, 108mM NaCl, 4mM NaHCO3, 1mM NaH2PO4, 5mM Trehalose and 10 mM Sucrose) and then transferred onto Poly l lysine (Gibco, A3890401) pretreated glass bottom petri dish (thickness 1,5) with Hemolymph-like solution for tissue preservation. A Chiaro nanoindentation device (Optics11), integrated above an inverted Axiovert 200M microscope (Zeiss), was used to probe gut mechanics using an adapted procedure previously described ^115^. A cantilever (stiffness (*K*)= 0.52-0.56 Nm^-1^) with a spherical tip (radius= 3.0-3.5 μm) was used to probe the midguts *ex vivo*. For the methylcellulose midgut nanoindentation, a cantilever (stiffness (*K*)= 0.42 Nm^-1^) with a spherical tip (radius= 12 μm) was used. The threshold and find surface values were set using an initial indentation. A matrix scan was configured along a 25-point line scan at intervals of 5µm along R4 or otherwise specified midgut regions. Scans were repeated twice in two parallel lines per region of interest for every midgut. The midguts were approached by the probe at 5 μm/s and indented with displacement parameters as defined in the displacement control mode.

#### *Ex vivo* live imaging

Flies were placed in vials containing filter paper discs with either 400 µl *Pe* or sucrose for indicated time prior to imaging. Coverslips were glued into imaging mounts. Flies were dissected in cold imaging medium (Schneider Insect medium (Sigma, 50146), 5% Fly Extract (VDRC), 1:1000 penicillin-streptomycin (Invitrogen, 15140), 10% FBS (Sigma, F4135) and placed on the coverslip (84). 10 µl Imaging Agarose (Low gelling temperature agarose 6% (Sigma, A4018), Schneider Insect medium (Thermo, 21720024) was placed on top of each gut to hold it to the surface of the coverslip and modified Schneider’s medium was added for imaging. Plates were sealed with lids containing damp paper to maintain humidity. Z stacks of the R4 region of the posterior midgut were acquired on a ZEISS 780.

#### *In vivo* live imaging

Flies were starved for two hours and then placed on *Pe* or sucrose solution for two hours prior to imaging. Flies were glued on their dorsal side and the A1 segment of abdominal cuticle removed. The annus of the flies was glued to prevent excretion of bacteria. A small portion of the R4 region was gently exposed with forceps and covered with 2% low gelling temperature agarose (Sigma, A4018) in Schneider’s Insect Medium (Sigma 50146). The dorsal side of the fly was then covered with Imaging medium (Schneider Insect Medium, 5 percent Fly Extract (VDRC), 0.1 percent Penicillin-Streptomycin (Invitrogen, 15140), 10 percent FBS (Sigma F4135). A coverslip was gently placed and secured over the dorsal side of the fly before imaging on an inverted Zeiss 880 confocal microscope. Acquisitions were taken at 5-minute intervals. Protocol adapted from ^116^.

#### Tracheal clone generation

Clones were generated by mating parental lines (*hsFLP; +; FRT82B, Gal80* and *dSRF-Gal4, UAS-GFP; ConLacZ, FRT82B*) for 3 days before transferring into fresh vials. The developing F_1_ offspring were then heat shocked 3 x 1h at day 9 post cross initiation at 37 °C. Adults were collected and aged for 7 days before dissecting midguts and confocal imaging on a ZEISS 780.

### Quantification Methods

#### A/P and D/V length and sarcomere length

Gut dorsal-ventral (DV) and anterior-posterior length (AP) were quantified using ImageJ (Version 1.54f) software. Length was measured using the freehand tool in the A/P axis of the posterior midgut. The D/V axis was measured from the widest point of the R4 region. For analysis of sarcomere length, phalloidin was used to detect the visceral muscle around the gut. Confocal Z stacks were acquired by a Zeiss 780. A Z projection was generated, and the freehand tool was used to measure length between two sarcomeres. Around 5-10 measurements were taken per image, and an average was taken per gut. Analysis was carried out in ImageJ (Version 1.54f).

#### ISC proliferation

Proliferating cells were detected by immunostaining against PH3. Olympus BX51fluorescence microscope was used to manually, on visual inspection, count the number of cells positive for PH3 staining per posterior midgut (R4 and R5 region).

#### Tracheal coverage

Tracheal tile scans were Z projected in ImageJ (Version 1.54f) and binarized from thresholding. A skeleton was then produced of the R4 and R5 regions of the gut and mean grey value was used to measure coverage of tracheal branches.

#### ROS fluorescence

For quantification of total ROS levels, pixel intensities of Z stacks from R4 (sum slices) were used. The mean of the summed DHE intensities averaged from each tissue was used for statistical analysis.

#### Gut contraction

Flies were starved for two hours and then placed on DSS, *Pe* or sucrose solution for two hours prior to imaging. Guts were then dissected in Schneider’s *Drosophila* medium (1x Sigma Aldrich 50146) at room temperature. Guts were then transferred to an imaging mount in a solution of room temperature Schneider’s medium and immediately imaged on a Leica M205 FA. Guts were imaged at 5 second intervals for a period of 5 minutes for each gut. The number of gut contractions was counted by visual inspection per 5-minute video.

#### Eccentricity

Images of the fly posterior midgut (R4) epithelium or mouse small intestine crypts were projected using the local z projector plugin in ImageJ (Version 1.54f) to produce a curvature corrected Z projection and heightmap. Z projected images were then segmented in Cellpose. Cellpose segmentations were converted to 4-bit connectivity images using an ImageJ plugin morpholibJ to create a cell mask ^29^. Masks and heightmaps from each sample were then analysed in Matlab using Deproj to analyse cellular eccentricity ^117^. 5 randomly selected intestinal crypts per mouse were imaged and segmented for eccentricity scoring.

#### Bayesian Force Inference of epithelium

Z projections and segmentation masks of R4/R5 tile scans were generated from Z-stacks, as above, using ‘local-z-projector’ and ‘cellpose’, respectively. Segmentation masks were then run through Tissue Analyser to produce coordinate data files of cell edges and vertices within the segmentation mask ^118^. Coordinate files were then processed in the Bayesian Force Inference programme in MATLAB to generate pressure data ^12^. Bayesian inference was performed using one gaussian prior of tension solved using QR decomposition via Suitesparse in MATLAB. The Bayesian Force Inference programme did not use Markov chain Monte Carlo but instead solved the bayesian chain inverse problem via optimisation using regularised linear algebra with Suitesparse.

#### Stiffness data analysis

(Nanoindentation and AFM)-Acquired force-displacement curves (*F-z*) were loaded in the graphical user interface (GUI) Nanoprepare ^119^, which was used to clean the data by deselecting any curves with failed contact or inappropriate noise. Collating this cleaned data set, formal data analysis was carried out on the forward segment of the curve in the GUI SoftMech (https://doi.org/10.5281/zenodo.19467492). The curves were then filtered to smooth any background noise generated by movement through the solution before contact. For AFM data from mouse cryosections, a Savitzky-Golay filter, was used to smooth the force indentation curves ^120^. For the midgut nanoindentation data a linear detrend filter was applied to the force-indentation curves. The segment denoting force applied (forward segment) to the sample was then isolated by applying a contact point identifying algorithm. Fit constant +line (nanite) ^121^ and Goodness of Fit (GoF) Sphere ^122^ algorithms were applied to the AFM and nanoindentation datasets respectively. To the force-indentation (*F*–δ) curves, the Hertz model (assuming Poisson ratio=0.5) was applied to the max indentation of δ = 0.1*R* to quantify the Young’s modulus (Pa) of the samples.

#### GRASP signal

GRASP positive cells within the R4 and R5 regions of the midgut were detected by immunostaining for GFP. Zeiss 780 confocal microscope was used to manually count the number of GRASP+ cells per posterior midgut.

#### Dcp1 signal

Z stacks of the R4 and R5 regions of the intestine were acquired using a Zeiss 780. Dcp1 positive cells were counted in the lumen of the posterior midgut manually, by visual inspection, to determine the number of apoptotic cells. Images were analysed using ImageJ (Version 1.54f).

#### *lacZ* reporters

Antibodies against β-galactosidase were used to detect *btl-lacZ*, *bnl-lacZ* and *ex-lacZ* reporters. Images of R4 were taken using confocal microscopy and staining was quantified using ImageJ. For *btl* and *bnl* quantifications, the background staining signal was subtracted from the total signal of β-galactosidase detected in the TTCs, ISCs/EBs or ECs. This value was then divided by the background signal to normalise the data. For ex-lacZ fluorescence intensity was quantified directly.

#### Myc and Bnl staining

Midguts stained with antibodies to detect these proteins included a methanol fixation step between the paraformaldehyde fixation and PBST wash steps of the standard protocol^63^. Images of R4 were acquired using confocal microscopy and staining was quantified using ImageJ (Version 1.54f). For each quantified gut, the background staining signal was subtracted from the total antibody signal in DAPI+ cells. This value was then divided by the background signal to normalize the data. In the case Supplementary Figs. 8g and 9d, total staining signal intensity was measured and ploted in the graphs.

#### Tracheal expression of *piezo* gene and Piezo protein trap

Quantifications were generated from a single z-stack of the R4 and R5 region for each midgut by confocal microscope and analysis was performed using ImageJ (Version 1.54f). A maximum intensity projection was produced for all images and channels were split. A skeleton was created in the channel corresponding to the gut trachea and threshold adjusted accordingly to ensure good selection of all tracheal branches. A region of interest was created from the trachea and applied to the Piezo protein trap expressing channel and fluorescence intensity quantified. For *piezo* gene expression, a skeleton was produced of piezo expressing trachea and total trachea, and coverage was calculated using mean grey value. A percentage of piezo positive trachea was then calculated by dividing piezo positive trachea by total tracheal coverage.

#### dSRF/Piezo quantification

To quantify the proportion of Piezo positive dSRF cells confocal microscope Z stacks of posterior midguts were obtained using a Zeiss 780 and analysed in ImageJ (Version 1.54f). A maximum intensity projection was produced for all images and channels were merged to show overlay of both d*SRF-mCherry* and *piezo*-GFP signal. Total dSRF positive tracheal cells were counted manually in the R4 and R5 regions and then the percentage of these dSRF positive cells which were also positive for *piezo-*GFP, were quantified.

#### Tracheal GCamp signal

Quantifications were generated from a single z-stack of the R4 region for each midgut by confocal microscopy and analysis was performed using ImageJ (Version 1.54f). A maximum intensity projection was produced for all images and channels were split. A skeleton was created in the channel corresponding to the gut trachea (red) and threshold adjusted accordingly to ensure good selection of tracheal branches. A region of interest was created from the active GCamp expressing trachea (green) and applied to the general trachea expressing channel (red) and fluorescence intensity quantified. For single cell GCamp dynamics, fluorescence intensity of individual tracheal body cells was quantified at each time point of the imaging time course.

#### Yki::GFP reporter

Quantifications were generated from a single Z stack of the R4 region of each midgut by confocal microscope and analysis was performed using ImageJ (Version 1.54f). A single Z plane was used to quantify each body cell where the nucleus was visible. Intensity of GFP was measured in the cytoplasm and the nucleus of each body cell and a ratio of nuclear to cytoplasmic intensity was generated to quantify nuclear translocation.

#### Ex::GFP reporter

Quantifications were generated from a single Z stack of the R4 region of each midgut by confocal microscope and analysis was performed using ImageJ (Version 1.54f). A single Z plane was used to quantify each body cell. Intensity of GFP was measured in the cytoplasm of each body cell. And average body cell intensity was generated per gut from 5-10 body cells.

## RT-qPCRs

*Piezo-Gal4* flies were crossed with either W1118, *Uas-piezoRNAi #105132* or *UAS*-*piezoRNAi #2796*. F1 offspring were then collected at the third instar larva stage and homogenised in 500 μL of Trizol (Thermo Fischer Scientific) using a mortar and a pestle. 200 μL of Chloroform was added and mixed well with the Trizol by vortexing before centrifugation on a bench top centrifuge. The aqueous phase was collected, and RNA was precipitated with iso-propanol. RNA was treated with DNase (Thermo Fischer Scientific) and quantified using a NanoDrop Spectrophotometer. Approximately 2 μg of RNA was converted to cDNA using a high-capacity cDNA reverse transcription kit (Thermo Fisher Scientific). RT-qPCR was performed using Perfecta SYBR green fast mix (Quantabio) as per manufacturer’s instructions on an Applied Biosystems QuantStudio 3 fast real-time PCR system and each reaction was carried out in three technical replicates per biological replicate. mRNA expression levels of target genes were normalized to the internal control *rpl32* using standard curves. Results shown represent average *piezo*/*rpl32* ratios of mRNA levels.

### Mouse models

All mouse experiments were performed in accordance with the UK Home Office and ARRIVE guidelines and were reviewed by the Animal Welfare and Ethical Review Board of the University of Glasgow. Experiments were performed under PPL PP7226959 and PIL <u>I33158383</u>. Transgenic mouse strains were backcrossed with C57Bl6J for 10 generations prior to experimental work. Mice were induced by 3mg Tamoxifen by oral gavage on two consecutive days. For regeneration experiments, mice were exposed to 10Gy whole body irradiation 5 days after Tamoxifen administration and then sampled 72 hours post radiation treatment. Unirradiated, age matched, tamoxifen treated mice were used for assessment of intestinal homeostasis. Males and female mice were used in the study. Intestines were flushed with cold PBS and sections taken for wholemount immunostaining and cryosections which were snap frozen in OCT. Ten x 1cm sections from the duodenum and jejunum were wrapped in surgical tape to make parcels and fixed in 4 percent paraformaldehyde (PFA) overnight for Haematoxylin and Eosin sections. For DSS treatment, female mice were given 1.5% DSS in drinking water for 5 consecutive days, 7 days after Tamoxifen treatment. Mice were sampled at day 8 after DSS treatment began or at humane endpoint prior to day 8. DAI score was calculated by scoring weight loss, stool consistency and rectal bleeding on a score of 0-4 and DAI was calculated as (weight loss score + stool consistency + rectal bleeding)/3. Weight loss score was categorised as 0=0% weight loss, 2=5-10% weight loss, 4=>15% weight loss. Stool consistency was categorised as 0=normal stool, 2=loose stool, 4=diarrhea. Rectal bleeding was categorised as 0=no bleeding, 2=slight bleeding, 4=gross bleeding. Colons were flushed with cold PBS, cut open lengthwise prior to fixation in 4% PFA overnight. Colons were then rolled into swiss roll preparations for immunohistochemistry processing.

### Immunohistochemistry of mouse tissue samples

All haematoxylin and eosin (H&E), immunohistochemistry (IHC), Alcian Blue (AB) and dual *in-situ* hybridization immunofluorescence (ISH-IF) staining was performed on 4µm formalin fixed paraffin embedded sections (FFPE) which had previously been ovened at 60 C for 2 hours. FFPE sections for c-Myc (ab32072, Abcam), CD31 (ab28364, Abcam), chromogranin A (ab15160, Abcam) and Sox 9 (AB5535, Millipore) detection were stained using an Agilent autostainer Link48. H&E staining was performed on a Leica autostainer (ST5020). Staining for AB was performed manually on FFPE sections that were dewaxed and rehydrated through xylene and a graded ethanol series before washing in water. Rehydrated slides were stained for 10 minutes in Alcian Blue solution before rinsing in tap water and then counterstained with Eosin for 30 seconds. <u>Irradiation experiments</u>: quantification of Myc and Sox9 expression, were done by manually counting the number of positively stained cells per small intestinal crypt and calculating this value as a percentage of total crypt cells. For Ki67, Alcian Blue and Chromogranin A, the total number of staining positive cells per small intestinal crypt was manually counted. A minimum of 5 crypts or crypt/villi were counted from each gut section present in a parcel and an average value was calculated for each mouse/small intestinal parcel. Total number of crypts were manually counted by visual inspection of H&E stained small intestinal parcels. Each data point represents the sum of the number of crypts present in all the gut sections of each small intestinal parcel/mouse. <u>For DSS experiments</u>: CD31 and Ki67 in ulcerated regions of the tissue were selected in QUPath and the ‘detect positive cells’ feature was used to calculate the percentage of positive cells in the ulcers for each staining. Images were acquired using a Grundium Ocus Slide Scanner and analysed using QUPath (Version 0.5.1).

### RNAscope of mouse tissue samples

Dual *ISH-IF* detection for *PECAM-1* (316728-C2) and *Piezo* 1 (500518-C1), 3-plex negative probe (320878) and 3-plex positive probe (320885) (all Bio-Techne) mRNA was performed using Multiplex fluorescent kit (322800; Bio-Techne) strictly according to the manufacturer’s instructions. The staining was visualised with Opal dyes (FP1487001KT, FP1488001KT, Akoya Biosciences). To complete staining, sections were placed in a 60°C oven for 30 minutes till dry, then sections were cover slipped using Ecomount (320409, Bio-Techne).

### Whole mount Immunostaining

Samples were prepared using a protocol adapted from ^86^. Sections of small intestine were flushed with cold PBS, cut open and pinned longitudinally on silicone plates. Sections were then fixed overnight in 4 percent PFA at 4 degrees on a shaker. Samples were then washed in PBS before washing in 10 percent sucrose in PBS and then 20 percent sucrose with 10 percent glycerol in PBS. Samples were then dipped in 0.1 percent sodium azide for sample preservation. Some sections were stored in sodium azide for up to 3 months at 4 degrees for further immunostaining. Samples were then washed in PBS before being placed in blocking buffer (PBS 5 percent donkey serum, 0.5 percent BSA, 0.3 percent Triton, 0.1 percent sodium azide) for 2 hours before incubation with primary antibody for 48 hours at 4 degrees.

Primary antibodies were diluted to the following concentrations in blocking buffer: chicken anti-GFP (1:400, Abcam), rat anti-PECAM-1 (1:400, Cell Signalling), rabbit anti-E-CADHERIN (1:500, Cell Signalling). Samples were then washed in blocking buffer before incubation in secondary antibody for 16 hours at 4 degrees. All secondary antibodies were diluted in blocking buffer to a concentration of 1:500. Samples were washed in wash buffer (PBS 0.3 percent Triton) before post fixation in 1 percent PFA at 4 degrees overnight. Samples were washed in PBS before mounting. Small strips 1-2 villi thick were cut from intestine sections and mounted in Vectashield with DAPI. Images were acquired on a Zeiss 880. Vascular coverage from 5 randomly selected crypt/villi units per mouse were quantified using the Mexican Hat filter and Skeletonise plugins on Z projections of villi in ImageJ (Version 1.54f). Epithelial quantifications were performed the same way as in the *Drosophila* model using LocalZprojector, cellpose and Deproj.

### Atomic Force Microscopy (AFM)

Small intestines from *Pdgf-*β*-CreER-GFP* mice were dissected and flushed to remove faecal content with ice-cold 1X PBS. The first 10 cm of the jejunum was discarded as this region tends to experience less irradiative damage. The downstream jejunum section was then cut into segments of 1cm, filled with optimal cutting temperature compound (OCT), aligned in sets of 3 in a cryomold and embedded in OCT. The tissue was fresh-frozen in a EtOH (100%) bath on dry-ice and stored at -80°C. 30μm sections were prepared at -20°C using a -15°C object temperature and transferred on to 0.28mg/ml Cell-Tak (Corning, 354240) treated (16 hours at 37°C, followed by 1 wash with water) 22mm x 22mm coverslips (thickness No 1, 5). Cover slips with adhered tissue sections were then glued to a 30mm dish and stored at -80°C. This protocol was adapted from ^61^. AFM experiments on the mouse small intestine cryosections were carried out under blinding of experimental conditions using a Drive AFM (Nanosurf) integrated above a Axio Observer fluorescent microscope (Zeiss) within 2 weeks from cryo-sectioning. Arrow TL (Nanoworld) cantilevers, stiffness (*K*) = 0.3 Nm^-1^ with a spherical bead (10μm in diameter) glued on the tip were used to probe small intestine cryosections. The samples were thawed in PBS supplemented with protease inhibitor cocktail (Roche 11873580001) and Pen/Strep (Gibco, 15140122). The nominal elastic constant and resonance frequency was determined using non-contact thermal calibration method^123^. The force setpoint and distance to record were set at 10Nn and 20μm, respectively. The cryosections were approached at 5 μm/s and measured in 4ml of PBS supplemented with protease inhibitor cocktail (Roche 11873580001) and Pen/Strep (Gibco, 15140122). For mechanical measurements maps were created using a spectroscopy grid (25 points in a 5μm x 5μm square) and muscle, crypt and villi areas from irradiated and control Pdgf-β-CreER-GFP mice.

## Statistics and Reproducibility

Except otherwise noted, n values represent data from 2 or 3 independent biological replicates. Each biological replicate represents a set of independent samples/experimental units (n) processed within the same day and belonging to animals derived from cultures with controlled feeding conditions, developmental timing and population density. Each biological replicate was processed in different days, and derives from independent F1s, which have been reared in independent batches of food.

In the case of qRT-PCR in Supplementary Figure 4i, ‘n’ refers to the number of biological replicates, which is each made of pulled whole larval tissue samples. Experiments in Figure 3m, 4d and Supplementary Figure 3i represent 6 biological replicates. Micrographs in Figures 1a, b and 2a are graphical exemplars of typical tissue anatomy and cellular morphologies that don’t directly correspond to any quantifications in this study but are well recognised and have been priorly characterized in other studies. Data in Figs. 1j, n-n’; 2 o-o’; 3d, k, l, o; 4b; 7f and Supplementary Figs. 2h, c, d; 4h; 7b, e, h, i; 9b represent single biological replicates with multiple sets of independent tissue samples (n) and that have been replicated in independent contexts throughout the manuscript or in previous studies. Supplementary videos 1-3 are representative of 3 independent movies each.

Data were analysed using one-way or two-way ANOVA with Sidak’s or Turkey’s multiple comparisons or T-test, where P>0.05 was statistically significant. For survival experiments, Kaplan-Meier survival curves were applied, and Mantel-Cox test was used. Results were analysed using Prism10 for Windows Version 10.6.1 (892). Individual sample data points are shown with group means presented as mean + SEM, where *P=0.05, ** P = 0.01, *** P=0.001 and **** P= 0.0001. Data is presented as mean + SD in Supplementary Figure 4i. Further information on sample size, statistical tests used and *P* values for each experiment are indicated in figure legends and corresponding source data files.

## Data availability

All raw data generated in this study has been deposited in institutional public repositories under accession code http://dx.doi.org/10.5525/gla.researchdata.2286 Numeric data generated in this study are provided in the Source Data file.

## Acknowledgements

We would like to thank Nic Tapon (The Francis Crick Institute), Norbert Perrimon (Harvard), Irene Miguel-Aliaga (The Francis Crick Institute), Shigeo Hayashi (Rinken Centre, Kobe, Japan), Chrysoula Pitsouli (University of Cyprus), Thomas Kornberg and Wanpeng Wang (Kornberg Lab-UCSF), Barry Denholm (University of Edinburgh), Barry Thompson (EMBL Australia), Gines Morata (Autonomous University of Madrid), Lucy O’Brien (Stanford University), Susumu Hirabayashi (Imperial College London), Magali Suzanne (CBI Toulouse), Yanlan Mao (UCL) and Andrew Davidson (University of Glasgow) for generously sharing flies and reagents, and Marcus Fruttiger (UCL) for sharing the *Pdgf-beta-Cre-GFP* mice. We thank the Bloomington Drosophila Stock Centre; the Vienna Drosophila Resource Center and the *Drosophila* Studies Hybridoma Bank for fly stocks and antibodies. We would like to thank the Core Services and Advanced Technologies at the CRUK Scotland Institute, which is core funded by Cancer Research UK (A31287). With particular thanks to Leo Carlin, Claire Mitchell, and Peter Thomason from the Beatson Advanced Imaging Resource for imaging support, Derek Miller and Evarest Onwubiko from the Biological Service Unit for assistance with animal whole body perfusions for whole mount intestinal sample preparations, and Dimitris Athineos and Dale Watt for assistance with the purchase and back crossing of the *Piezo1^fl^* mice. We are grateful to Magali Suzanne for critical comments and advice on the manuscript, and Jeremiah Bernier-Latmani and Tatiana Petrova for their support and advice during whole mount intestinal vascular tissue preparation and imaging. The manuscript was critically reviewed by Catherine Winchester (CRUK Scotland Institute-RRID:SCR_027384). We thank all members of the Cordero laboratory for general advice on the project.

## Authors contributions

J.A.P. Designed, performed and analysed most experiments, including the work on tissue segmentation, morphometric analysis, and GRASP and mouse experiments.

J.P. Designed, performed and analysed experiments, including the study of signalling mechanisms downstream of Piezo. Contributed to study conceptualization, supervision and funding acquisition.

C.T.J. Designed, performed and analysed all experiments involving the use of Chiaro nanoindenter, AFM and the time course of Piezo expression and activation.

M.W. Provided training and assistance with nanoindentation data acquisition and analysis.

N. and M.H. Processed samples for mammalian histology.

A.B.C. Contributed to early efforts in live imaging.

Y.Y. Provided general technical assistance.

L.M and K.B. Provided support with the supervision and planning of the mouse work.

M.V. Planning, supervision and analysis of nanoindentation and AFM work and funding acquisition.

J.B.C. Study conceptualization, supervision, data analysis and funding acquisition.

J.A.P., J.P., C.T.J, and J.B.C., Wrote the manuscript with input from the rest of the authors.

---

**Movie 1: Intracellular Calcium in gut associated trachea of homeostatic midguts.** Time-lapse *in vivo* live imaging of a control *Drosophila* posterior midgut from Sucrose fed mated female fly. The general tracheal network was labelled with *Q6>mtdTomato* (red) and the calcium reporter *GCamp6s* (green) is expressed under the control of the tracheal driver *dSRF-gal4*. Fly was starved and then fed sucrose for 2 hours prior to imaging for 3 hours.

**Movie 2: Increased intracellular Calcium in gut associated trachea of infected midguts.** Time-lapse *in vivo* live imaging of a control *Drosophila* posterior midgut from *Pe* fed mated female fly. The general tracheal network was labelled with *Q6>mtdTomato* (red) and the calcium reporter *GCamp6s* (green) is expressed under the control of the tracheal driver *dSRF-gal4*. Fly was starved and then fed *Pe* for 2 hours prior to imaging for 3 hours.

**Movie 3: *Piezo* dependent increased intracellular Calcium in gut associated trachea of infected midguts.** Time-lapse *in vivo* live imaging of a *Drosophila* posterior midgut from *Pe* fed mated female fly. The general tracheal network was labelled with *Q6>mtdTomato* (red) and the calcium reporter *GCamp6s* (green) and a *piezo-RNAi* (*piezo-IR1*) are expressed under the control of the tracheal driver *dSRF-gal4*. Fly was starved and then fed *Pe* for 2 hours prior to imaging for 3 hours.

## References

1. Hageman JH, Heinz MC, Kretzschmar K, Van Der Vaart J, Clevers H, Snippert HJG. Intestinal Regeneration: Regulation by the Microenvironment. Developmental Cell. 2020;54(4):435–46.

2. Sato T, Van Es JH, Snippert HJ, Stange DE, Vries RG, Van Den Born M, et al. Paneth cells constitute the niche for Lgr5 stem cells in intestinal crypts. Nature. 2011;469(7330):415–8.

3. Li H, Jasper H. Gastrointestinal stem cells in health and disease: from flies to humans. Disease Models & Mechanisms. 2016;9(5):487–99.

4. Ding B-S, Cao Z, Lis R, Nolan DJ, Guo P, Simons M, et al. Divergent angiocrine signals from vascular niche balance liver regeneration and fibrosis. Nature. 2014;505(7481):97–102.

5. Rafii S, Butler JM, Ding B-S. Angiocrine functions of organ-specific endothelial cells. Nature. 2016;529(7586):316–25.

6. Rivas-Carrillo SD, Kanamune J, Iwanaga Y, Uemoto S, Daneri-Navarro A, Rivas-Carrillo JD. Endothelial Cells Promote Pancreatic Stem Cell Activation During Islet Regeneration in Mice. Transplantation Proceedings. 2011 2011/11/01/;43(9):3209–11.

7. Perochon J, Yu Y, Aughey GN, Medina AB, Southall TD, Cordero JB. Dynamic adult tracheal plasticity drives stem cell adaptation to changes in intestinal homeostasis in Drosophila. Nature Cell Biology. 2021;23(5):485–96.

8. Tamamouna V, Rahman MM, Petersson M, Charalambous I, Kux K, Mainor H, et al. Remodelling of oxygen-transporting tracheoles drives intestinal regeneration and tumorigenesis in Drosophila. Nature Cell Biology. 2021;23(5):497–510.

9. Gerit, Jacobson J, Karl, Hudry B, Christo, Dormann D, et al. Neuronal Control of Metabolism through Nutrient-Dependent Modulation of Tracheal Branching. Cell. 2014;156(1-2):69–83.

10. Blackie L, Gaspar P, Mosleh S, Lushchak O, Kong L, Jin Y, et al. The sex of organ geometry. Nature. 2024;630(8016):392–400.

11. Kam CY, Singh ID, Gonzalez DG, Matte-Martone C, Solá P, Solanas G, et al. Mechanisms of skin vascular maturation and maintenance captured by longitudinal imaging of live mice. Cell. 2023 2023/05/25/;186(11):2345–60.e16.

12. Kong W, Loison O, Chavadimane Shivakumar P, Chan EH, Saadaoui M, Collinet C, et al. Experimental validation of force inference in epithelia from cell to tissue scale. Scientific Reports. 2019;9(1).

13. Ichijo R, Kabata M, Kidoya H, Muramatsu F, Ishibashi R, Abe K, et al. Vasculature-driven stem cell population coordinates tissue scaling in dynamic organs. Science Advances. 2021;7(7):eabd2575.

14. Ghabrial A, Luschnig S, Metzstein MM, Krasnow MA. Branching Morphogenesis of the Drosophila Tracheal System. Annual Review of Cell and Developmental Biology. 2003;19(Volume 19, 2003):623–47.

15. Centanin L, Gorr TA, Wappner P. Tracheal remodelling in response to hypoxia. Journal of Insect Physiology. 2010 2010/05/01/;56(5):447–54.

16. Eilken HM, Adams RH. Dynamics of endothelial cell behavior in sprouting angiogenesis. Current Opinion in Cell Biology. 2010 2010/10/01/;22(5):617–25.

17. Affolter M, Bellusci S, Itoh N, Shilo B, Thiery J-P, Werb Z. Tube or Not Tube. Developmental Cell. 2003;4(1):11–8.

18. Hayashi S, Kondo T. Development and Function of the *Drosophila* Tracheal System. Genetics. 2018;209(2):367–80.

19. Buchon N, Broderick NA, Chakrabarti S, Lemaitre B. Invasive and indigenous microbiota impact intestinal stem cell activity through multiple pathways in Drosophila. Genes Dev. 2009 Oct 1;23(19):2333–44. PubMed PMID: 19797770. PMCID: PMC2758745. eng.

20. Buchon N, Broderick NA, Poidevin M, Pradervand S, Lemaitre B. Drosophila Intestinal Response to Bacterial Infection: Activation of Host Defense and Stem Cell Proliferation. Cell Host & Microbe. 2009 2009/02/19/;5(2):200–11.

21. Jiang H, Patel PH, Kohlmaier A, Grenley MO, McEwen DG, Edgar BA. Cytokine/Jak/Stat Signaling Mediates Regeneration and Homeostasis in the Drosophila Midgut. Cell. 2009 2009/06/26/;137(7):1343–55.

22. Amcheslavsky A, Jiang J, Ip YT. Tissue Damage-Induced Intestinal Stem Cell Division in Drosophila. Cell Stem Cell. 2009;4(1):49–61.

23. Howard AM, LaFever KS, Fenix AM, Scurrah CR, Lau KS, Burnette DT, et al. DSS-induced damage to basement membranes is repaired by matrix replacement and crosslinking. Journal of Cell Science. 2019;132(7).

24. Morris O, Jasper H. Reactive Oxygen Species in intestinal stem cell metabolism, fate and function. Free Radical Biology and Medicine. 2021 2021/04/01/;166:140–6.

25. Farnsworth RH, Lackmann M, Achen MG, Stacker SA. Vascular remodeling in cancer. Oncogene. 2014;33(27):3496–505.

26. Osol G, Moore LG. Maternal Uterine Vascular Remodeling During Pregnancy. Microcirculation. 2014 2014/01/01;21(1):38–47.

27. Spradley FT, Ge Y, Granger JP, Chade AR. Utero-placental vascular remodeling during late gestation in Sprague-Dawley rats. Pregnancy Hypertension. 2020 2020/04/01/;20:36–43.

28. Buchon N, Broderick NA, Kuraishi T, Lemaitre B. DrosophilaEGFR pathway coordinates stem cell proliferation and gut remodeling following infection. BMC Biology. 2010;8(1):152.

29. Stringer C, Wang T, Michaelos M, Pachitariu M. Cellpose: a generalist algorithm for cellular segmentation. Nature Methods. 2021;18(1):100–6.

30. Chen J, St Johnston D. Epithelial Cell Polarity During Drosophila Midgut Development. Front Cell Dev Biol. 2022;10:886773. PubMed PMID: 35846367. PMCID: PMC9281564. Epub 20220630. eng.

31. Bischofs IB, Klein F, Lehnert D, Bastmeyer M, Schwarz US. Filamentous Network Mechanics and Active Contractility Determine Cell and Tissue Shape. Biophysical Journal. 2008 2008/10/01/;95(7):3488–96.

32. Grosser S, Lippoldt J, Oswald L, Merkel M, Sussman DM, Renner F, et al. Cell and Nucleus Shape as an Indicator of Tissue Fluidity in Carcinoma. Physical Review X. 2021 02/17/;11(1):011033.

33. Lecuit T, Lenne P-F. Cell surface mechanics and the control of cell shape, tissue patterns and morphogenesis. Nature Reviews Molecular Cell Biology. 2007 2007/08/01;8(8):633–44.

34. Paluch E, Heisenberg C-P. Biology and Physics of Cell Shape Changes in Development. Current Biology. 2009 2009/09/15/;19(17):R790–R9.

35. Rape AD, Guo WH, Wang YL. The regulation of traction force in relation to cell shape and focal adhesions. Biomaterials. 2011 Mar;32(8):2043–51. PubMed PMID: 21163521. PMCID: PMC3029020. Epub 20101215. eng.

36. Zemel A, Rehfeldt F, Brown AE, Discher DE, Safran SA. Cell shape, spreading symmetry and the polarization of stress-fibers in cells. J Phys Condens Matter. 2010 May 19;22(19):194110. PubMed PMID: 20458358. PMCID: PMC2865697. eng.

37. Curcio EJ, Lubkin SR. Physical models of notochord cell packing reveal how tension ratios determine morphometry. Cells & Development. 2023 2023/03/01/;173:203825.

38. Kong W, Loison O, Chavadimane Shivakumar P, Chan EH, Saadaoui M, Collinet C, et al. Experimental validation of force inference in epithelia from cell to tissue scale. Sci Rep. 2019 Oct 10;9(1):14647. PubMed PMID: 31601854. PMCID: PMC6787039. Epub 20191010. eng.

39. Monier B, Gettings M, Gay G, Mangeat T, Schott S, Guarner A, et al. Apico-basal forces exerted by apoptotic cells drive epithelium folding. Nature. 2015 2015/02/01;518(7538):245–8.

40. Nestor-Bergmann A, Goddard G, Woolner S, Jensen OE. Relating cell shape and mechanical stress in a spatially disordered epithelium using a vertex-based model. Mathematical Medicine and Biology: A Journal of the IMA. 2018;35(Supplement_1):i1–i27.

41. Pomp W, Schakenraad K, Balcıoğlu HE, van Hoorn H, Danen EHJ, Merks RMH, et al. Cytoskeletal Anisotropy Controls Geometry and Forces of Adherent Cells. Physical Review Letters. 2018 10/26/;121(17):178101.

42. Sorrell EL, Lubkin SR. Bubble packing, eccentricity, and notochord development. Cells & Development. 2022 2022/03/01/;169:203753.

43. He L, Si G, Huang J, Samuel ADT, Perrimon N. Mechanical regulation of stem-cell differentiation by the stretch-activated Piezo channel. Nature. 2018;555(7694):103–6.

44. Buchon N, Osman D, David FPA, Fang HY, Boquete J-P, Deplancke B, et al. Morphological and molecular characterization of adult midgut compartmentalization in Drosophila. Cell Rep. 2013 2013/05/30/;3(5):1725–38. eng.

45. Marianes A, Spradling AC. Physiological and stem cell compartmentalization within the Drosophila midgut. Elife. 2013 Aug 27;2:e00886. PubMed PMID: 23991285. PMCID: PMC3755342. Epub 2013/08/31.

46. Zwick RK, Kasparek P, Palikuqi B, Viragova S, Weichselbaum L, McGinnis CS, et al. Epithelial zonation along the mouse and human small intestine defines five discrete metabolic domains. Nat Cell Biol. 2024 Feb;26(2):250–62. PubMed PMID: 38321203. PMCID: PMC11654995. Epub 20240206. eng.

47. Kong W, Loison O, Chavadimane Shivakumar P, Chan EH, Saadaoui M, Collinet C, et al. Experimental validation of force inference in epithelia from cell to tissue scale. Scientific Reports. 2019 2019/10/10;9(1):14647.

48. Brückner BR, Janshoff A. Elastic properties of epithelial cells probed by atomic force microscopy. Biochimica et Biophysica Acta (BBA)-Molecular Cell Research. 2015 2015/11/01/;1853(11, Part B):3075–82.

49. Feinberg EH, Vanhoven MK, Bendesky A, Wang G, Fetter RD, Shen K, et al. GFP Reconstitution Across Synaptic Partners (GRASP) Defines Cell Contacts and Synapses in Living Nervous Systems. Neuron. 2008;57(3):353–63.

50. Oates AC, Gorfinkiel N, González-Gaitán M, Heisenberg C-P. Quantitative approaches in developmental biology. Nature Reviews Genetics. 2009 2009/08/01;10(8):517-30.

51. Aguilar-Cuenca R, Juanes-García A, Vicente-Manzanares M. Myosin II in mechanotransduction: master and commander of cell migration, morphogenesis, and cancer. Cell Mol Life Sci. 2014 Feb;71(3):479–92. PubMed PMID: 23934154. PMCID: PMC11113847. Epub 20130811. eng.

52. Franke JD, Montague RA, Kiehart DP. Nonmuscle Myosin II Generates Forces that Transmit Tension and Drive Contraction in Multiple Tissues during Dorsal Closure. Current Biology. 2005 2005/12/24/;15(24):2208–21.

53. Garrido-Casado M, Asensio-Juárez G, Talayero VC, Vicente-Manzanares M. Engines of change: Nonmuscle myosin II in mechanobiology. Current Opinion in Cell Biology. 2024 2024/04/01/;87:102344.

54. Islam ST, Tang Y, Boliver H, Bi D, Fowler VM. Nonmuscle myosin IIA (NMIIA) regulates anisotropic cell tension to maintain the hexagonal packing of mouse lens meridional row cells. Mol Biol Cell. 2025 Oct 1;36(10):ar124. PubMed PMID: 40833808. PMCID: PMC12483326. Epub 20250820. eng.

55. Weißenbruch K, Grewe J, Hippler M, Fladung M, Tremmel M, Stricker K, et al. Distinct roles of nonmuscle myosin II isoforms for establishing tension and elasticity during cell morphodynamics. eLife. 2021 2021/08/10;10:e71888.

56. Xie Z, Rose L, Feng J, Zhao Y, Lu Y, Kane H, et al. Enteric neuronal Piezo1 maintains mechanical and immunological homeostasis by sensing force. Cell. 2025 May 1;188(9):2417–32.e19. PubMed PMID: 40132579. PMCID: PMC12048284. Epub 20250324. eng.

57. Schott S, Ambrosini A, Barbaste A, Benassayag C, Gracia M, Proag A, et al. A fluorescent toolkit for spatiotemporal tracking of apoptotic cells in living *Drosophila* tissues. Development. 2017;144(20):3840–6.

58. Rosenblatt J, Raff MC, Cramer LP. An epithelial cell destined for apoptosis signals its neighbors to extrude it by an actin-and myosin-dependent mechanism. Curr Biol. 2001 Nov 27;11(23):1847–57. PubMed PMID: 11728307. eng.

59. Toyama Y, Peralta XG, Wells AR, Kiehart DP, Edwards GS. Apoptotic Force and Tissue Dynamics During *Drosophila* Embryogenesis. Science. 2008;321(5896):1683–6.

60. Monier B, Gettings M, Gay G, Mangeat T, Schott S, Guarner A, et al. Apico-basal forces exerted by apoptotic cells drive epithelium folding. Nature. 2015 Feb 12;518(7538):245–8. PubMed PMID: 25607361. Epub 20150121. eng.

61. Baghdadi MB, Houtekamer RM, Perrin L, Rao-Bhatia A, Whelen M, Decker L, et al. PIEZO-dependent mechanosensing is essential for intestinal stem cell fate decision and maintenance. Science. 2024;386(6725):eadj7615.

62. Kim C-K, Yang VW, Bialkowska AB. The Role of Intestinal Stem Cells in Epithelial Regeneration Following Radiation-Induced Gut Injury. Current Stem Cell Reports. 2017 2017/12/01;3(4):320–32.

63. Johansson J, Naszai M, Hodder MC, Pickering KA, Miller BW, Ridgway RA, et al. RAL GTPases Drive Intestinal Stem Cell Function and Regeneration through Internalization of WNT Signalosomes. Cell Stem Cell. 2019;24(4):592–607.e7.

64. Fischer KS, Henn D, Zhao ET, Sivaraj D, Litmanovich B, Hahn WW, et al. Elevated Shear Stress Modulates Heterogenous Cellular Subpopulations to Induce Vascular Remodeling. Tissue Engineering Part A. 2024 2024/12/01;30(23-24):752–65.

65. Sun Y, Liu J, Xu Z, Lin X, Zhang X, Li L, et al. Matrix stiffness regulates myocardial differentiation of human umbilical cord mesenchymal stem cells. Aging. 2021;13(2):2231–50.

66. Zanotelli MR, Reinhart-King CA. Mechanical Forces in Tumor Angiogenesis. Springer International Publishing; 2018. p. 91–112.

67. Negri S, Faris P, Berra-Romani R, Guerra G, Moccia F. Endothelial Transient Receptor Potential Channels and Vascular Remodeling: Extracellular Ca2 + Entry for Angiogenesis, Arteriogenesis and Vasculogenesis. Frontiers in Physiology. 2020;10.

68. Coste B, Mathur J, Schmidt M, Earley TJ, Ranade S, Petrus MJ, et al. Piezo1 and Piezo2 Are Essential Components of Distinct Mechanically Activated Cation Channels. Science. 2010;330(6000):55–60.

69. Ranade SS, Qiu Z, Woo S-H, Hur SS, Murthy SE, Cahalan SM, et al. Piezo1, a mechanically activated ion channel, is required for vascular development in mice. Proceedings of the National Academy of Sciences. 2014;111(28):10347–52.

70. Kim SE, Coste B, Chadha A, Cook B, Patapoutian A. The role of Drosophila Piezo in mechanical nociception. Nature. 2012;483(7388):209–12.

71. Zechini L, Amato C, Scopelliti A, Wood W. Piezo acts as a molecular brake on wound closure to ensure effective inflammation and maintenance of epithelial integrity. Current Biology. 2022 2022/08/22/;32(16):3584–92.e4.

72. Nagarkar-Jaiswal S, Lee P-T, Campbell ME, Chen K, Anguiano-Zarate S, Cantu Gutierrez M, et al. A library of MiMICs allows tagging of genes and reversible, spatial and temporal knockdown of proteins in Drosophila. eLife. 2015 2015/03/31;4:e05338.

73. Oh Y, Lai JS, Min S, Huang HW, Liberles SD, Ryoo HD, et al. Periphery signals generated by Piezo-mediated stomach stretch and Neuromedin-mediated glucose load regulate the Drosophila brain nutrient sensor. Neuron. 2021 Jun 16;109(12):1979–95.e6. PubMed PMID: 34015253. PMCID: PMC8611812. Epub 20210519. eng.

74. Song Y, Li D, Farrelly O, Miles L, Li F, Kim SE, et al. The Mechanosensitive Ion Channel Piezo Inhibits Axon Regeneration. Neuron. 2019 Apr 17;102(2):373–89.e6. PubMed PMID: 30819546. PMCID: PMC6487666. Epub 20190225. eng.

75. Wang P, Jia Y, Liu T, Jan YN, Zhang W. Visceral Mechano-sensing Neurons Control Drosophila Feeding by Using Piezo as a Sensor. Neuron. 2020 Nov 25;108(4):640–50.e4. PubMed PMID: 32910893. PMCID: PMC8386590. Epub 20200909. eng.

76. Chakrabarti S, Liehl P, Buchon N, Lemaitre B. Infection-Induced Host Translational Blockage Inhibits Immune Responses and Epithelial Renewal in the Drosophila Gut. Cell Host & Microbe. 2012 2012/07/19/;12(1):60–70.

77. Niec RE, Chu T, Schernthanner M, Gur-Cohen S, Hidalgo L, Pasolli HA, et al. Lymphatics act as a signaling hub to regulate intestinal stem cell activity. Cell Stem Cell. 2022 2022/07/07/;29(7):1067–82.e18.

78. Wang P, Kljavin N, Nguyen TTT, Storm EE, Marsh B, Jiang J, et al. Adrenergic nerves regulate intestinal regeneration through IL-22 signaling from type 3 innate lymphoid cells. Cell Stem Cell. 2023 2023/09/07/;30(9):1166–78.e8.

79. Bernier-Latmani J, Petrova TV. Intestinal lymphatic vasculature: structure, mechanisms and functions. Nature Reviews Gastroenterology & Hepatology. 2017 2017/09/01;14(9):510–26.

80. Bernier-Latmani J, González-Loyola A, Petrova TV. Mechanisms and functions of intestinal vascular specialization. Journal of Experimental Medicine. 2024;221(1).

81. Stzepourginski I, Nigro G, Jacob J-M, Dulauroy S, Sansonetti PJ, Eberl G, et al. CD34 ^+^ mesenchymal cells are a major component of the intestinal stem cells niche at homeostasis and after injury. Proceedings of the National Academy of Sciences. 2017;114(4):E506–E13.

82. McCarthy N, Manieri E, Storm EE, Saadatpour A, Luoma AM, Kapoor VN, et al. Distinct Mesenchymal Cell Populations Generate the Essential Intestinal BMP Signaling Gradient. Cell Stem Cell. 2020;26(3):391–402.e5.

83. Kim J-E, Fei L, Yin W-C, Coquenlorge S, Rao-Bhatia A, Zhang X, et al. Single cell and genetic analyses reveal conserved populations and signaling mechanisms of gastrointestinal stromal niches. Nature Communications. 2020;11(1).

84. Palikuqi B, Rispal J, Reyes EA, Vaka D, Boffelli D, Klein O. Lymphangiocrine signals are required for proper intestinal repair after cytotoxic injury. Cell Stem Cell. 2022;29(8):1262–72.e5.

85. Claxton S, Kostourou V, Jadeja S, Chambon P, Hodivala-Dilke K, Fruttiger M. Efficient, inducible Cre-recombinase activation in vascular endothelium. genesis. 2008;46(2):74–80.

86. Bernier-Latmani J, Petrova TV. High-resolution 3D analysis of mouse small-intestinal stroma. Nature Protocols. 2016;11(9):1617–29.

87. Nusse R, Clevers H. Wnt/β-Catenin Signaling, Disease, and Emerging Therapeutic Modalities. Cell. 2017 Jun 1;169(6):985–99. PubMed PMID: 28575679. eng.

88. Blache P, Van De Wetering M, Duluc I, Domon C, Berta P, Freund J-NL, et al. SOX9 is an intestine crypt transcription factor, is regulated by the Wnt pathway, and represses the *CDX2* and *MUC2* genes. The Journal of Cell Biology. 2004;166(1):37–47.

89. Liu L, Rao JN, Zou T, Xiao L, Smith A, Zhuang R, et al. Activation of Wnt3a signaling stimulates intestinal epithelial repair by promoting c-Myc-regulated gene expression. American Journal of Physiology-Cell Physiology. 2012;302(1):C277–C85. PubMed PMID: 21975427.

90. Czarnewski P, Parigi SM, Sorini C, Diaz OE, Das S, Gagliani N, et al. Conserved transcriptomic profile between mouse and human colitis allows unsupervised patient stratification. Nature Communications. 2019 2019/06/28;10(1):2892.

91. Melgar S, Karlsson L, Rehnström E, Karlsson A, Utkovic H, Jansson L, et al. Validation of murine dextran sulfate sodium-induced colitis using four therapeutic agents for human inflammatory bowel disease. Int Immunopharmacol. 2008 Jun;8(6):836–44. PubMed PMID: 18442787. Epub 20080307. eng.

92. Mim MS, Kumar N, Levis M, Unger MF, Miranda G, Gazzo D, et al. Piezo regulates epithelial topology and promotes precision in organ size control. Cell Rep. 2024;43(7).

93. Mikesell AR, Isaeva O, Moehring F, Sadler KE, Menzel AD, Stucky CL. Keratinocyte PIEZO1 modulates cutaneous mechanosensation. eLife. 2022 2022/09/02;11:e65987.

94. Yang Y, Wang D, Zhang C, Yang W, Li C, Gao Z, et al. Piezo1 mediates endothelial atherogenic inflammatory responses via regulation of YAP/TAZ activation. Human Cell. 2022;35(1):51–62.

95. Lan Y, Lu J, Zhang S, Jie C, Chen C, Xiao C, et al. Piezo1-Mediated Mechanotransduction Contributes to Disturbed Flow-Induced Atherosclerotic Endothelial Inflammation. Journal of the American Heart Association. 2024;13(21):e035558.

96. Mao J, Yang R, Yuan P, Wu F, Wei Y, Nie Y, et al. Different stimuli induce endothelial dysfunction and promote atherosclerosis through the Piezo1/YAP signaling axis. Archives of Biochemistry and Biophysics. 2023 2023/10/01/;747:109755.

97. Pan D. The hippo signaling pathway in development and cancer. Dev Cell. 2010 Oct 19;19(4):491–505. PubMed PMID: 20951342. PMCID: PMC3124840. eng.

98. Badouel C, Gardano L, Amin N, Garg A, Rosenfeld R, Le Bihan T, et al. The FERM-Domain Protein Expanded Regulates Hippo Pathway Activity via Direct Interactions with the Transcriptional Activator Yorkie. Developmental Cell. 2009 2009/03/17/;16(3):411–20.

99. Chen P, Zhang G, Jiang S, Ning Y, Deng B, Pan X, et al. Mechanosensitive Piezo1 in endothelial cells promotes angiogenesis to support bone fracture repair. Cell Calcium. 2021 2021/07/01/;97:102431.

100. Kang H, Hong Z, Zhong M, Klomp J, Bayless KJ, Mehta D, et al. Piezo1 mediates angiogenesis through activation of MT1-MMP signaling. American Journal of Physiology-Cell Physiology. 2019;316(1):C92–C103.

101. Peralta M, Dupas A, Larnicol A, Lefebvre O, Goswami R, Stemmelen T, et al. Endothelial calcium firing mediates the extravasation of metastatic tumor cells. iScience. 2025 2025/02/21/;28(2):111690.

102. Boeldt DS, Bird IM. Vascular adaptation in pregnancy and endothelial dysfunction in preeclampsia. Journal of Endocrinology. 2017;232(1):R27–R44.

103. Guelfi S, Hodivala-Dilke K, Bergers G. Targeting the tumour vasculature: from vessel destruction to promotion. Nature Reviews Cancer. 2024;24(10):655–75.

104. Fraisl P, Mazzone M, Schmidt T, Carmeliet P. Regulation of Angiogenesis by Oxygen and Metabolism. Developmental Cell. 2009;16(2):167–79.

105. Gervais L, Bardin AJ. Tracheal remodelling supports stem cells. Nature Cell Biology. 2021;23(6):580–2.

106. Fortuño A, José GS, Moreno MU, Díez J, Zalba G. Oxidative stress and vascular remodelling. Experimental Physiology. 2005;90(4):457–62.

107. Centanin L, Dekanty A, Romero N, Irisarri M, Gorr TA, Wappner P. Cell Autonomy of HIF Effects in Drosophila: Tracheal Cells Sense Hypoxia and Induce Terminal Branch Sprouting. Developmental Cell. 2008;14(4):547–58.

108. Mukherjee P, El-Abbadi MM, Kasperzyk JL, Ranes MK, Seyfried TN. Dietary restriction reduces angiogenesis and growth in an orthotopic mouse brain tumour model. British Journal of Cancer. 2002;86(10):1615–21.

109. Kawaue T, Yow I, Pan Y, Le AP, Lou Y, Loberas M, et al. Inhomogeneous mechanotransduction defines the spatial pattern of apoptosis-induced compensatory proliferation. Developmental Cell. 2023 2023/02/27/;58(4):267–77.e5.

110. Stricker AM, Hutson MS, Page-McCaw A. Piezo-dependent surveillance of matrix stiffness generates transient cells that repair the basement membrane. Dev Cell. 2025 Jul 21;60(14):1936–46.e4. PubMed PMID: 40081371. PMCID: PMC12286749. Epub 20250312. eng.

111. Bautista GM, Du Y, Matthews MJ, Flores AM, Kushnir NR, Sweeney NK, et al. Smooth muscle cell Piezo1 depletion results in impaired contractile properties in murine small bowel. Communications Biology. 2025 2025/03/17;8(1):448.

112. Min S, Oh Y, Verma P, Whitehead SC, Yapici N, Van Vactor D, et al. Control of feeding by Piezo-mediated gut mechanosensation in Drosophila. eLife. 2021 2021/02/18;10:e63049.

113. Medina A, Bellec K, Polcowñuk S, Cordero JB. Investigating local and systemic intestinal signalling in health and disease with Drosophila. Dis Model Mech. 2022 Mar 1;15(3). PubMed PMID: 35344037. PMCID: PMC8990086. Epub 20220328. eng.

114. Medina AB, Perochon J, Tian Y, Johnson CT, Holcombe J, Ramesh P, et al. Neuroendocrine control of intestinal regeneration through the vascular niche in Drosophila. Dev Cell. 2025 Nov 17;60(22):3085–101.e6. PubMed PMID: 40695286. Epub 20250721. eng.

115. Ciccone G, Azevedo Gonzalez Oliva M, Antonovaite N, Lüchtefeld I, Salmeron-Sanchez M, Vassalli M. Experimental and Data Analysis Workflow for Soft Matter Nanoindentation. JoVE. 2022 2022/01/18(179):e63401.

116. Martin JL, Sanders EN, Moreno-Roman P, Jaramillo Koyama LA, Balachandra S, Du X, et al. Long-term live imaging of the Drosophila adult midgut reveals real-time dynamics of division, differentiation and loss. eLife. 2018 2018/11/14;7:e36248.

117. Herbert S, Valon L, Mancini L, Dray N, Caldarelli P, Gros J, et al. LocalZProjector and DeProj: a toolbox for local 2D projection and accurate morphometrics of large 3D microscopy images. BMC Biology. 2021;19(1).

118. Aigouy B, Umetsu D, Eaton S. Segmentation and Quantitative Analysis of Epithelial Tissues. Springer New York; 2016. p. 227–39.

119. Ciccone G, Azevedo Gonzalez Oliva M, Antonovaite N, Lüchtefeld I, Salmeron-Sanchez M, Vassalli M. Experimental and Data Analysis Workflow for Soft Matter Nanoindentation. Journal of Visualized Experiments. 2022 (179).

120. Virtanen P, Gommers R, Oliphant TE, Haberland M, Reddy T, Cournapeau D, et al. SciPy 1.0: fundamental algorithms for scientific computing in Python. Nature Methods. 2020 2020/03/01;17(3):261–72.

121. Müller P, Abuhattum S, Möllmert S, Ulbricht E, Taubenberger AV, Guck J. nanite: using machine learning to assess the quality of atomic force microscopy-enabled nano-indentation data. BMC Bioinformatics. 2019 2019/09/10;20(1):465.

122. Gavara N. Combined strategies for optimal detection of the contact point in AFM force-indentation curves obtained on thin samples and adherent cells. Scientific Reports. 2016 2016/02/19;6(1):21267.

123. Higgins MJ, Proksch R, Sader JE, Polcik M, Mc Endoo S, Cleveland JP, et al. Noninvasive determination of optical lever sensitivity in atomic force microscopy. Review of Scientific Instruments. 2006;77(1).

