## Supplementary Figures and Tables for "Biomechanical Compartmentalisation Influences Angiogenesis and Stem Cell Proliferation in the Regenerating Intestine via Vascular Piezo"

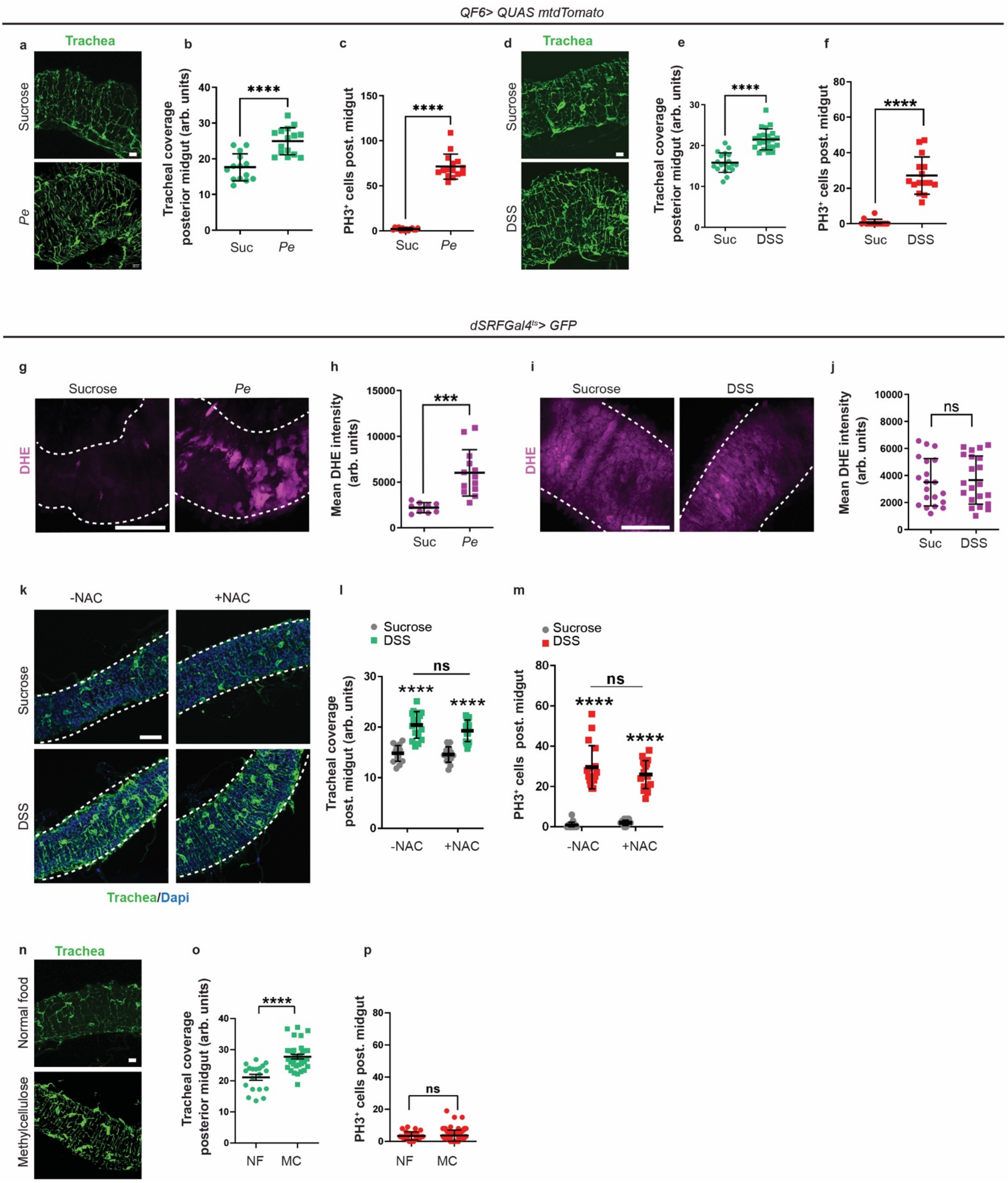
QF6> QUAS mtdTomato
a
Trachea
Sucrose
Pe
b
Tracheal coverage posterior midgut (arb. units)
****
Suc
Pe
c
PH3+ cells post. midgut
****
Suc
Pe
d
Trachea
Sucrose
DSS
e
Tracheal coverage posterior midgut (arb. units)
****
Suc
DSS
f
PH3+ cells post. midgut
****
Suc
DSS
dSRFGal4ts> GFP
g
Sucrose
Pe
DHE
h
Mean DHE intensity (arb. units)
***
Suc
Pe
i
Sucrose
DSS
DHE
j
Mean DHE intensity (arb. units)
ns
Suc
DSS
k
-NAC
+NAC
Sucrose
DSS
Trachea/Dapi
l
Sucrose
DSS
Tracheal coverage post. midgut (arb. units)
ns
****
****
-NAC
+NAC
m
Sucrose
DSS
PH3+ cells post. midgut
ns
****
****
-NAC
+NAC
n
Trachea
Normal food
Methylcellulose
o
Tracheal coverage posterior midgut (arb. units)
****
NF
MC
p
PH3+ cells post. midgut
ns
NF
MC

**Supplementary Figure 1. Biomechanical changes in the intestine correlate with tracheal remodelling. a.** Confocal images of the adult *Drosophila* midgut associated trachea (green) upon sucrose and *Pe* feeding. Scale bars, 20 mm. **b, c**. Quantification of tracheal remodelling (**b**) (n=14,15) and ISC proliferation (**c**) (n=14, 15) as in a. **d.** Midgut associated trachea (green) in the context of sucrose and DSS feeding. Scale bars, 20 mm. **e, f**. Quantification of tracheal remodelling (**e**) (n=19,22) and ISC proliferation (n=13, 15) (**f**) as in d. **g**. Intestinal reactive oxygen species detection (DHE; pink) in sucrose and *Pe* fed flies. Scale bar, 50 mm. **h**. Quantification of DHE intensity in midguts as in g (n=9,13). **i**. DHE staining (red) in sucrose and DSS fed flies. Scale bars, 50 mm. **j**. Quantification of gut DHE intensity in midguts as in I (n=20,21). **k**. Adult *Drosophila* midgut associated trachea (green) upon sucrose and DSS feeding with the antioxidant N-acetyl-cysteine (NAC+) or sucrose (NAC-). Scale bars, 50 mm. **l, m**. Quantification of tracheal remodelling (**l**) and ISC proliferation (**m**) in midguts as in k (n=17,17,16,17) (n=17,17,16,17). **n**. *Drosophila* midgut trachea following normal food (NF) and methylcellulose (MC) feeding. Scale bars, 20 mm. **o, p**. Quantification of tracheal remodelling (**o**) and ISC proliferation (**p**) in midguts as in n (n=19,23) (n=28,99). Data represent the mean ± SEM. n=numbers of guts analysed. <u>Statistics</u>: **b, c, e, f, h, j, o, p**. Two-tailed unpaired Student's t-test. **l, m**. Two-way ANOVA followed by Tukey's multiple comparisons test.

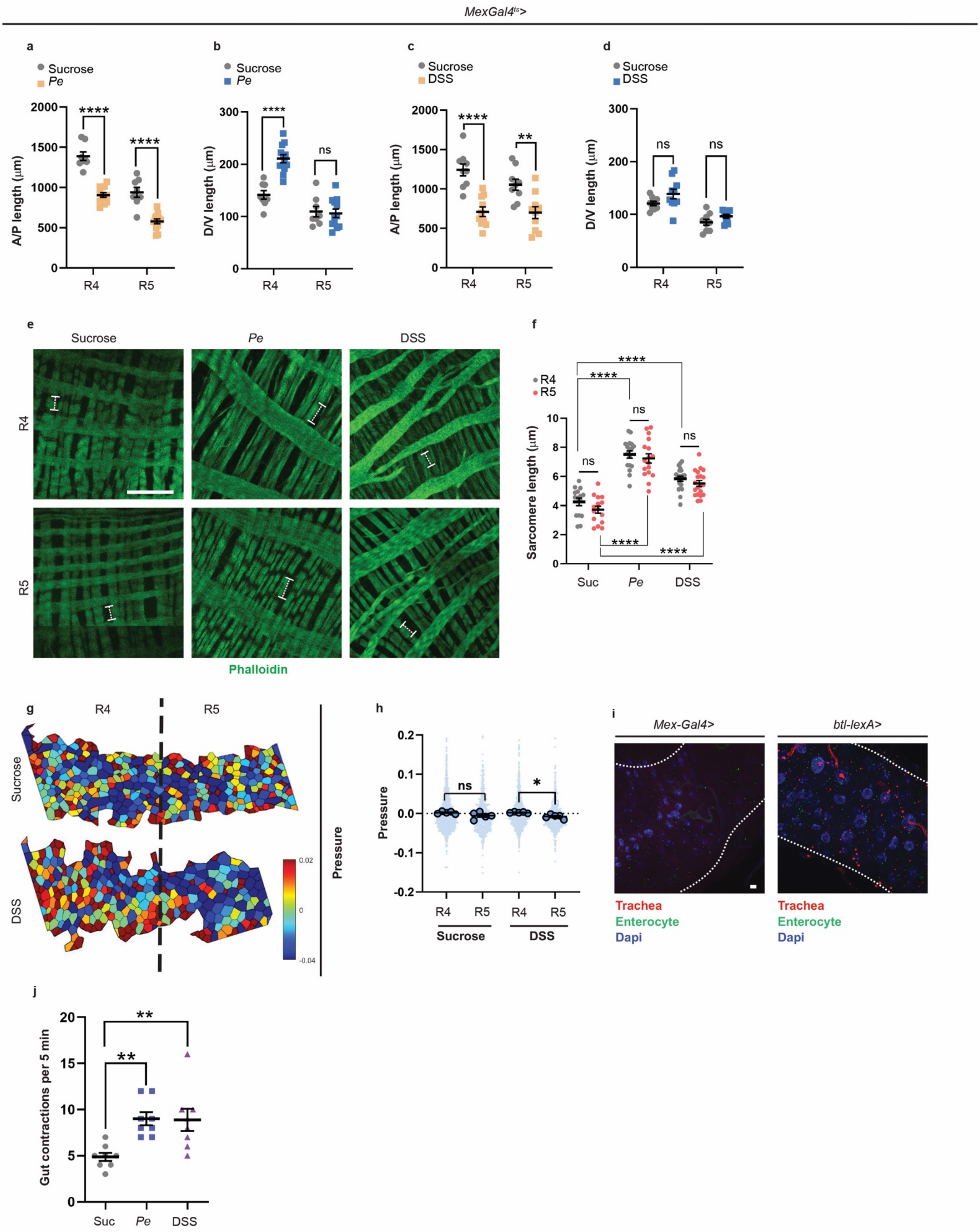
MexGal4ts>
a
Sucrose
Pe
A/P length (μm)
R4
R5
b
Sucrose
Pe
D/V length (μm)
c
Sucrose
DSS
A/P length (μm)
d
Sucrose
DSS
D/V length (μm)
e
Sucrose
Pe
DSS
R4
R5
Phalloidin
f
R4
R5
Sarcomere length (μm)
Suc
Pe
DSS
g
R4
R5
Sucrose
DSS
Pressure
h
Pressure
Sucrose
DSS
i
Mex-Gal4>
btl-lexA>
Trachea
Enterocyte
Dapi
j
Gut contractions per 5 min
Suc
Pe
DSS

**Supplementary Figure 2. Morphological changes are observed in the epithelial and muscle compartment of the regenerating midgut. a, b.** Quantification of anterior-posterior (A.P.) (**a**) and dorsal-ventral (D.V.) (**b**) length of the R4 and R5 compartments of the posterior midgut following sucrose or *Pe* treatment (n=8,12,8,12) (n=8,12,8,12). **c, d.** Quantification of anterior-posterior (A.P.) (**c**) and dorsal-ventral (D.V.) (**d**) length of the R4 and R5 compartments of the posterior midgut following sucrose or DSS treatment (n=9,10,9,10) (n=9,10,9,10). **e**. Confocal images of phalloidin staining (green) to detect visceral muscle in the R4 and R5 regions of the posterior midgut following sucrose, *Pe* or DSS treatment. Scale bar, 20 mm. **f**. Quantification of sarcomere length in conditions in e (n=15,15,17,17,20,20). **g**. Representative image of inferred relative cell pressure calculated using Bayesian Force Inference from the posterior midgut of flies treated with sucrose or DSS. **h**. Quantification of relative pressure in midguts as in g. Smaller circles represent individual cells and larger circles in represent average of cells per gut (n=5,5,5,5). **i**. Controls for GRASP system without *btl-lexA* driver or *Mex-Gal4* driver system. Scale bar, 50 mm. **j**. Quantification of the number of gut contractions during a five-minute imaging period of ex vivo cultured guts from flies fed with sucrose, *Pe* or DSS (n=8,8,8). Data represent the mean ± SEM. n=numbers of guts analysed. <u>Statistics</u>: Two-way ANOVA followed by Tukey’s multiple comparisons test.

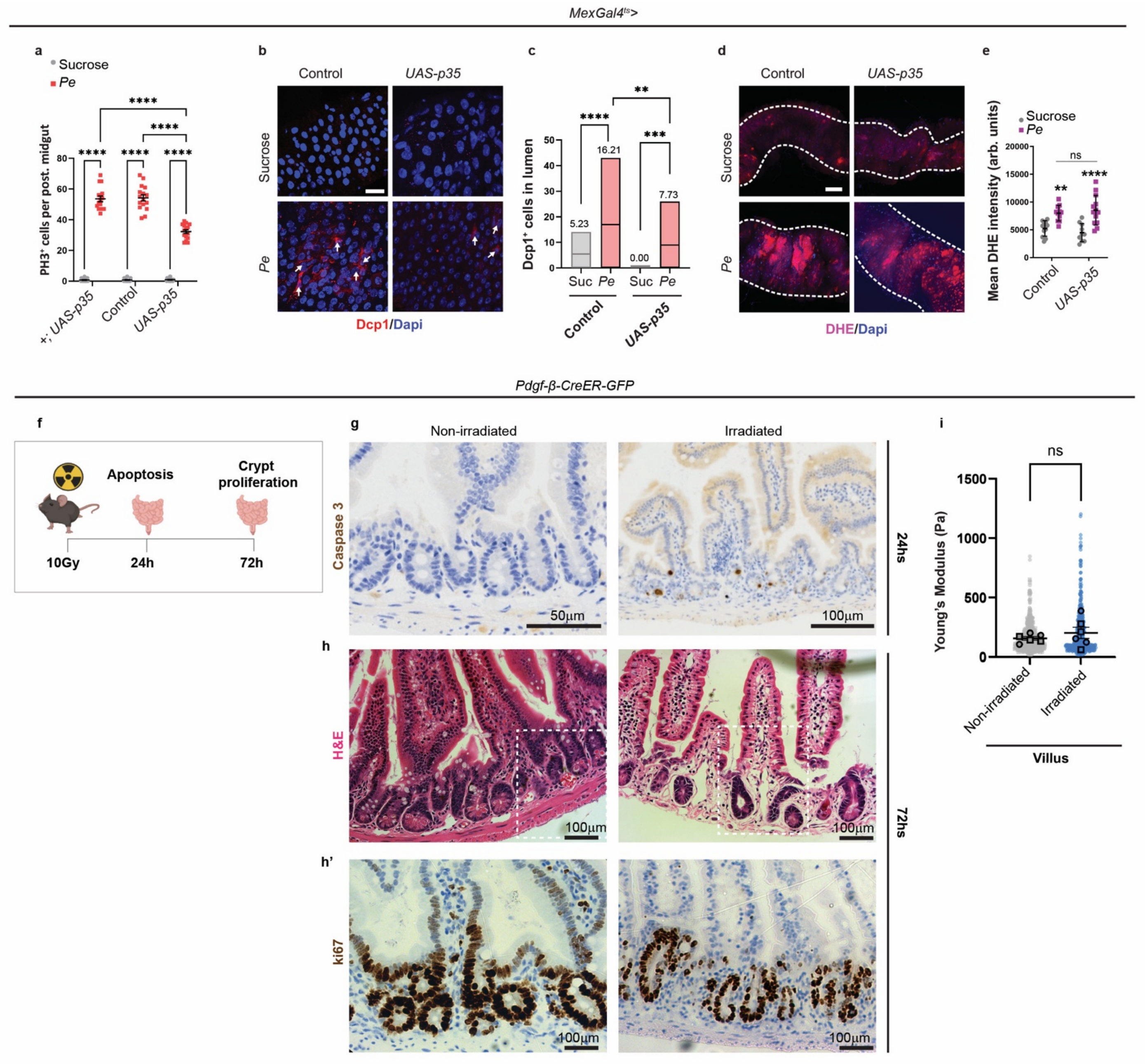

MexGal4ts>
a
Sucrose
Pe
PH3+ cells per post. midgut
+; UAS-p35
Control
UAS-p35
b
Control
UAS-p35
Sucrose
Pe
Dcp1/Dapi
c
Dcp1+ cells in lumen
5.23
16.21
0.00
7.73
Suc
Pe
Control
UAS-p35
d
Control
UAS-p35
Sucrose
Pe
DHE/Dapi
e
Sucrose
Pe
Mean DHE intensity (arb. units)
ns
Control
UAS-p35
Pdgf-β-CreER-GFP
f
Apoptosis
Crypt proliferation
10Gy
24h
72h
g
Non-irradiated
Irradiated
Caspase 3
50µm
100µm
24hs
h
H&E
100µm
h'
ki67
100µm
72hs
i
ns
Young's Modulus (Pa)
Non-irradiated
Irradiated
Villus

**Supplementary Figure 3. Inhibiting cell death does not influence gut ROS.** **a.** Quantification of ISC proliferation upon p35 overexpression in enterocytes (*Mex*$^{ts}$*>p35*) and in genetic background controls (n=18,16,18,16,16,15). **b**. *Drosophila* caspase-1 (Dcp1; arrows) staining in the R4 and R5 regions of sucrose or *Pe* treated flies in control or *Mex*$^{ts}$*>p35* midguts. Scale bar, 20 mm. **c**. Quantification of the number of Dcp1+ cells in the posterior midgut lumen as in b (n=16,21,26,25); box plots represent maxima, minima and mean. Mean value is indicated on top of boxes. **d**. Intestinal DHE staining (pink) in sucrose and *Pe* fed flies from control or *Mex*$^{ts}$*>p35* midguts. Scale bar, 50 mm. **e**. Quantification of DHE intensity in midguts as in d (n=11,11,10,13). **f**. Representative schematic of whole-body gamma irradiation (IR) time course. *Created in BioRender. Cordero, J. (2026) https://BioRender.com/c7qij6w*. **g**. Representative images of apoptotic marker, cleaved caspase-3 in healthy and irradiated mice, 24h post IR. Scale bars, 50 and 100 mm respectively. **h**. H&E staining of the small intestine from healthy and irradiated mice 72h post IR. Scale bar, 100 mm. **h'**. Ki67 staining of small intestine from healthy and irradiated mice 72h post IR. Scale bar, 100 mm. **i**. Quantification of stiffness (Young's modulus) in the villus region of healthy vs irradiated mice 72h post IR (n=6,6). Smaller dots represent individual cells, and larger circles represent average stiffness per gut. n= numbers of guts/mice. Data represent the mean ± SEM. Statistics: Two-way ANOVA followed by Tukey's multiple comparisons test.

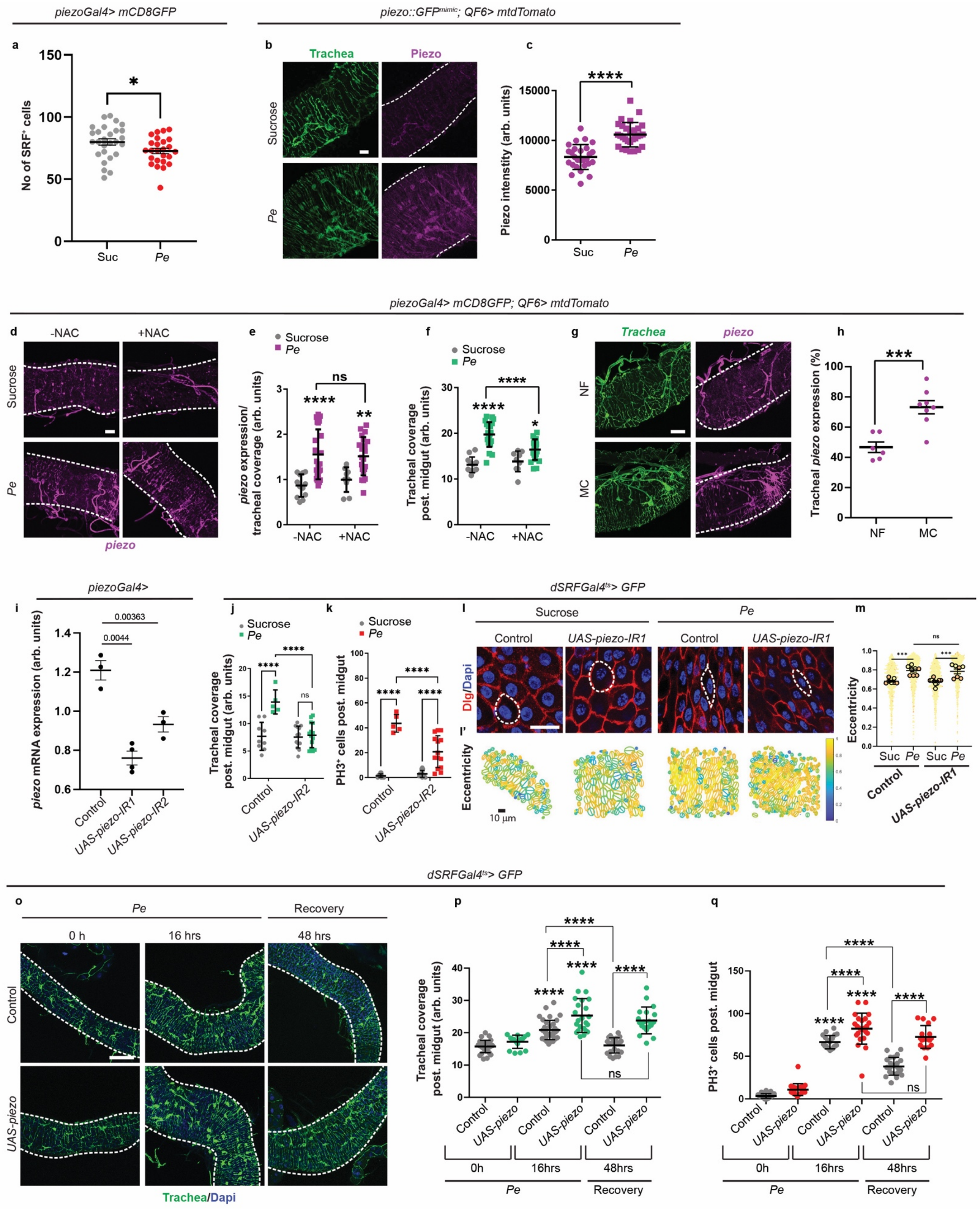
piezoGal4> mCD8GFP
piezo::GFPmimic; QF6> mtdTomato
a
No of SRF+ cells
Suc
Pe
b
Trachea
Piezo
Sucrose
Pe
c
Piezo intensity (arb. units)
Suc
Pe
piezoGal4> mCD8GFP; QF6> mtdTomato
d
-NAC
+NAC
Sucrose
Pe
piezo
e
Sucrose
Pe
piezo expression/ tracheal coverage (arb. units)
-NAC
+NAC
ns
f
Sucrose
Pe
Tracheal coverage post. midgut (arb. units)
-NAC
+NAC
g
Trachea
piezo
NF
MC
h
Tracheal piezo expression (%)
NF
MC
piezoGal4>
i
piezo mRNA expression (arb. units)
0.00363
0.0044
Control
UAS-piezo-IR1
UAS-piezo-IR2
dSRFGal4ts> GFP
j
Sucrose
Pe
Tracheal coverage post. midgut (arb. units)
ns
Control
UAS-piezo-IR2
k
Sucrose
Pe
PH3+ cells post. midgut
Control
UAS-piezo-IR2
l
Sucrose
Pe
Control
UAS-piezo-IR1
Control
UAS-piezo-IR1
Dlg/Dapi
l'
Eccentricity
10 µm
m
Eccentricity
ns
Suc
Pe
Suc
Pe
Control
UAS-piezo-IR1
dSRFGal4ts> GFP
o
Pe
Recovery
0 h
16 hrs
48 hrs
Control
UAS-piezo
Trachea/Dapi
p
Tracheal coverage post. midgut (arb. units)
ns
Control
UAS-piezo
Control
UAS-piezo
Control
UAS-piezo
0h
16hrs
48hrs
Pe
Recovery
q
PH3+ cells post. midgut
ns
Control
UAS-piezo
Control
UAS-piezo
Control
UAS-piezo
0h
16hrs
48hrs
Pe
Recovery

**Supplementary Figure 4. Mechanical perturbations in the midgut are sensed by tracheal Piezo. a.** Number of SRF+ tracheal cell nuclei in sucrose and *Pe* treated midguts (n=27,25). **b**. Midgut trachea (green) and Piezo expression (magenta) in sucrose and *Pe* . Scale bar, 20 mm. **c**. Quantification of tracheal Piezo as in a (n=27,29)**. d.** *piezo* expression (pink) following sucrose and *Pe* infection with (NAC+) or sucrose (NAC-). Scale bar, 20 mm. **e, f**. Quantification of tracheal *piezo* (**e**) and coverage (**f**) as in d (n=12,23,9,21) (n=12,23,9,21). **g**. Midgut trachea (green) and *piezo* expression (magenta) after normal (NF) or methylcellulose (MC) food. Scale bar, 50 mm. **h**. Quantification of tracheal piezo expression in midguts as in f (n= 6,8). **i**. Larval *piezo* mRNA levels following gene knockdown by RNAi. (n=3,4,3); n= biological replicates. Data represent mean ± SD. **j, k**. Quantification of tracheal remodelling (**j**) and ISC proliferation (**k**) upon sucrose and *Pe* infection in control and tracheal piezo knockdown (*piezo-IR*) midguts (n=9, 5, 13, 14) (n=9, 5, 13, 14). **l**. Midgut epithelium segmentation (anti-Dlg; red) upon sucrose and *Pe* infection in control and tracheal *piezo-IR*. Dashed lines outline single cells. Scale bar, 20 mm. **l'**. Representative images of epithelial eccentricity in midguts as in l. Scale bar, 10 mm. **m**. Quantification of epithelial eccentricity as in l. Smaller circles indicate individual cell values and large circles the average cell eccentricity value per gut. Each colour indicates a different biological replicate (n=7,8,9,7). **o**. Control or *piezo* overexpressing midgut trachea (green) upon 0 and 16hs *Pe* infection or 48hs after *Pe* removal (recovery). Scale bar, 20 mm. **p, q**. Quantification of tracheal coverage (**p**) (n=29,16,36,23,23,19) and ISC proliferation (**q**) (n=25,22,20,23,22,17) in midguts as in o. Data represent the mean ± SEM. Unless otherwise stated, n=numbers of guts. Statistics: **a, c, h**. Two-tailed unpaired Student's t-test. **e, f, j, k, m, p, q.** Two-way ANOVA followed by Tukey's multiple comparisons test. **i.** Two-way ANOVA followed by Dunnett's multiple comparisons test; p values as indicated in figure.

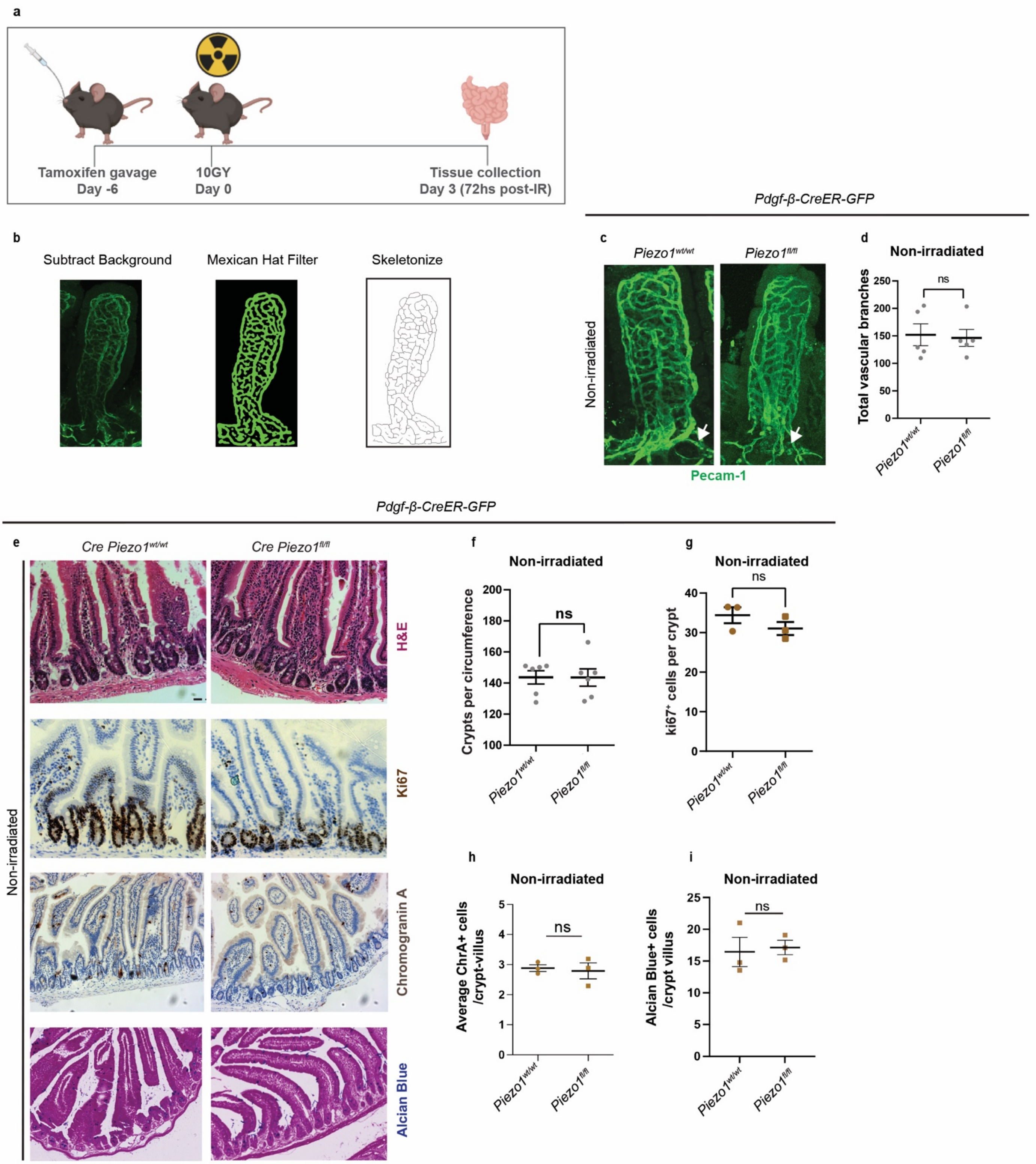
a
Tamoxifen gavage
Day -6
10GY
Day 0
Tissue collection
Day 3 (72hs post-IR)
Pdgf-β-CreER-GFP
b
Subtract Background
Mexican Hat Filter
Skeletonize
c
Piezo1wt/wt
Piezo1fl/fl
Non-irradiated
Pecam-1
d
Non-irradiated
Total vascular branches
ns
Pdgf-β-CreER-GFP
e
Cre Piezo1wt/wt
Cre Piezo1fl/fl
H&E
Ki67
Chromogranin A
Alcian Blue
Non-irradiated
f
Non-irradiated
Crypts per circumference
ns
g
Non-irradiated
ki67+ cells per crypt
ns
h
Non-irradiated
Average ChrA+ cells /crypt-villus
ns
i
Non-irradiated
Alcian Blue+ cells /crypt villus
ns

**Supplementary Figure 5. The role of vascular *Piezo1* is conserved in an irradiation induced damage model in the mammalian small intestine. a.** Experimental design for gene induction and whole-body gamma irradiation in mice. *Created in BioRender. Cordero, J. (2026) https://BioRender.com/7jxf72k*. **b**. Method used to segment and quantify vascular branching in whole mount immunostaining of the intestinal vasculature using anti-PECAM-1 antibody. Scale bar, 20 mm. **c**. Whole mount immunostaining of anti-PECAM-1 (green) in crypt villus images of healthy, *Pdgf-beta-CreER* $Piezo1^{wt/wt}$ and $Piezo1^{fl/fl}$ mice. **d**. Quantification of the number of vascular branches (arrows) in mice as in c (n=5,5). **e**. H&E, Ki67, chromogranin A and Alcian blue staining of small intestinal samples from healthy *Pdgf-beta-CreER* $Piezo1^{wt/wt}$ and $Piezo1^{fl/fl}$ mice. Scale bar, 20 mm. **f, g, h, i**. Quantification of number of crypts per circumference (**f**), percentage Ki67+ cells per crypt (**g**) and the number of chromogranin A (**h**) and Alcian blue (**i**) positive cells per crypt villus axis (n=6,6) (n=3,3) (n=3,3) (n=3,3). Data represent the mean ± SEM. n=numbers of guts/mice analysed. Statistics: Two-tailed unpaired Student's t-test.

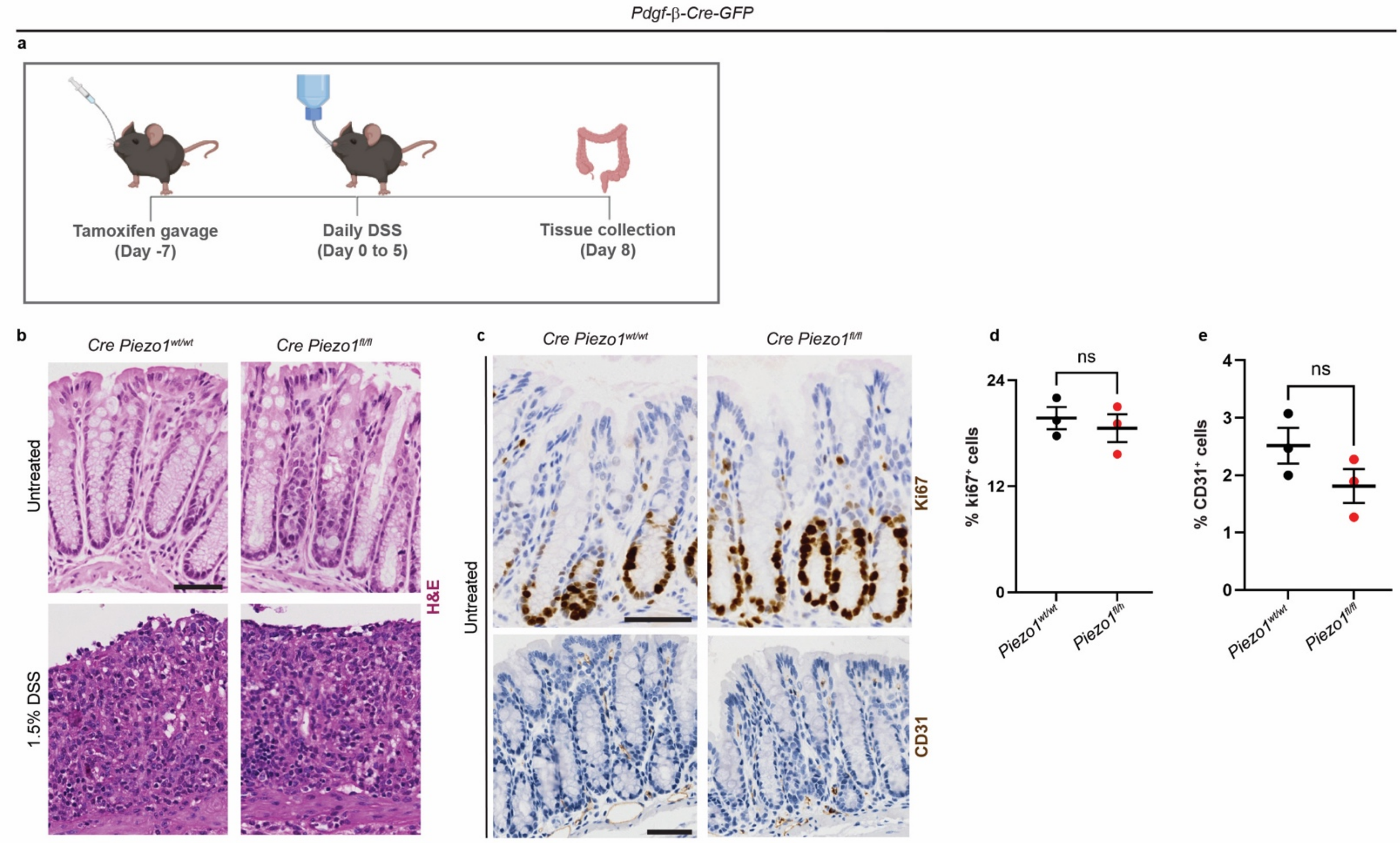

**Supplementary Figure 6. The role of vascular *Piezo1* is conserved in a DSS damage model in the mammalian colon. a.** Experimental design for gene induction and DSS treatment in mice. *Created in BioRender. Cordero, J. (2026)* https://BioRender.com/7jxf72k. **b**. H&E staining of colon samples from healthy and DSS treated *Pdgf-beta-CreER Piezo1*$^{wt/wt}$ and *Piezo1*$^{fl/fl}$ mice. Scale bar, 20 mm. **c**. Ki67 and CD31 staining of colon samples from healthy *Pdgf-beta-CreER Piezo1*$^{wt/wt}$ and *Piezo1*$^{fl/fl}$ mice. Scale bar, 20 mm. **d, e**. Quantification of percentage of Ki67 (**d**) and CD31 (**e**) cells in healthy colon tissue as in c (n=3,3) (n=3,3). Data represent the mean ± SEM. n=numbers of guts/mice analysed. Statistics: Two-tailed unpaired Student's t-test.

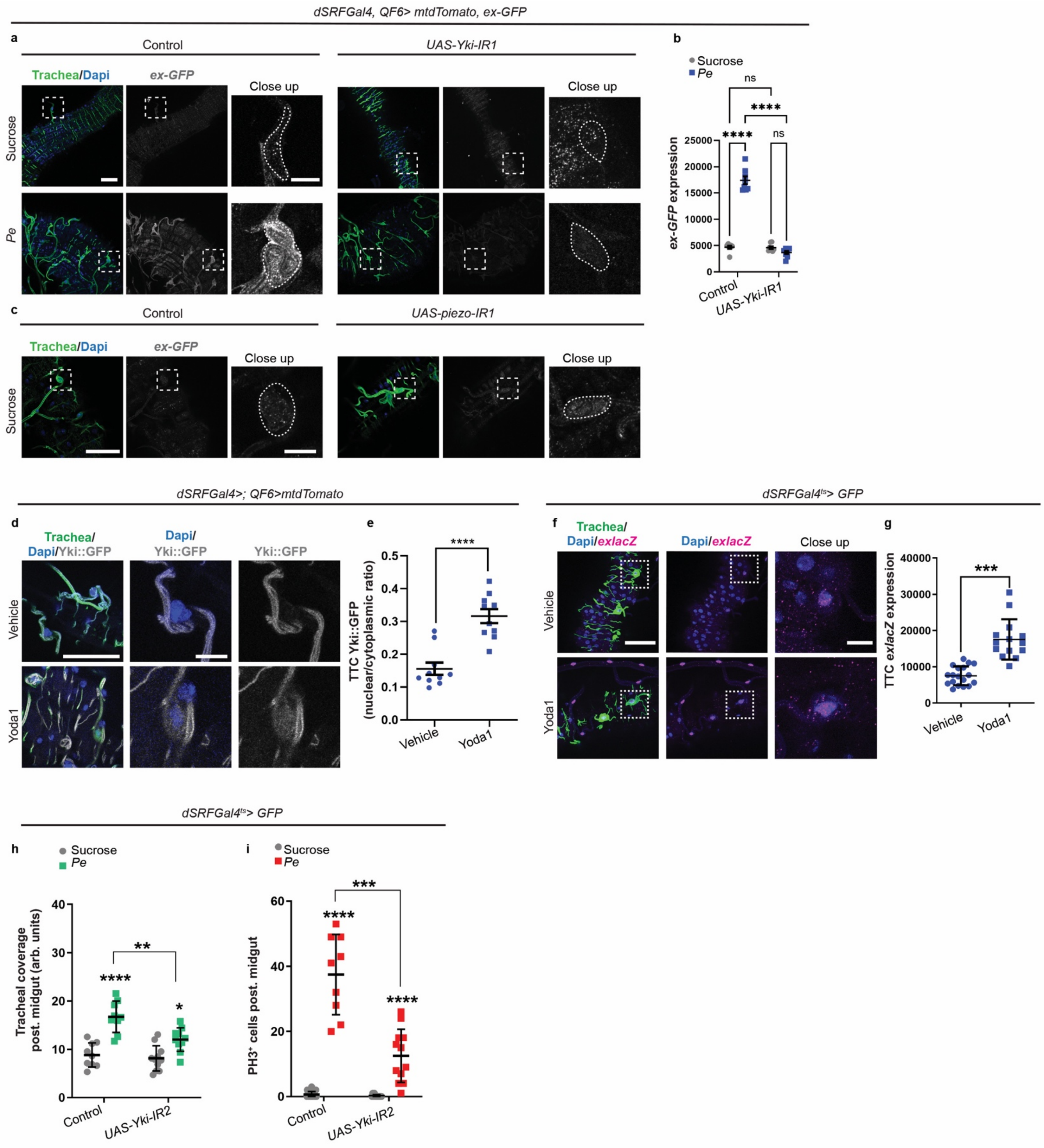

dSRFGal4, QF6> mtdTomato, ex-GFP
a
Control
UAS-Yki-IR1
Trachea/Dapi
ex-GFP
Close up
Sucrose
Pe
b
Sucrose
Pe
ns
****
****
ns
ex-GFP expression
25000
20000
15000
10000
5000
0
Control
UAS-Yki-IR1
c
Control
UAS-piezo-IR1
Trachea/Dapi
ex-GFP
Close up
Sucrose
dSRFGal4>; QF6>mtdTomato
d
Trachea/Dapi/Yki::GFP
Dapi/Yki::GFP
Yki::GFP
Vehicle
Yoda1
e
****
TTC Yki::GFP (nuclear/cytoplasmic ratio)
0.5
0.4
0.3
0.2
0.1
0.0
Vehicle
Yoda1
dSRFGal4ts> GFP
f
Trachea/Dapi/exlacZ
Dapi/exlacZ
Close up
Vehicle
Yoda1
g
***
TTC exlacZ expression
40000
30000
20000
10000
0
Vehicle
Yoda1
dSRFGal4ts> GFP
h
Sucrose
Pe
**
****
*
Tracheal coverage post. midgut (arb. units)
40
30
20
10
0
Control
UAS-Yki-IR2
i
Sucrose
Pe
***
****
****
PH3+ cells post. midgut
60
40
20
0
Control
UAS-Yki-IR2

**Supplementary Figure 7. Yki is activated downstream of Piezo in intestinal trachea.** **a.** Confocal images of Ex-GFP (grey) in midgut trachea (green) following sucrose and *Pe* feeding in control or tracheal *yki* knockdown (*yki-IR1*) midguts. Scale bar, 50 mm, 10 mm. **b**. Quantification of tracheal Ex-GFP expression as in a (n=8,8,8,8). **c**. Confocal images of Ex-GFP (grey) in midgut trachea (green) following sucrose feeding in control or tracheal *piezo* knockdown (*piezo-IR1*) midguts. Scale bar, 50 mm, 10 mm. **d**. Representative confocal images of midgut associated tracheal tissue (green) and Yki::GFP (grey) in midguts from vehicle or Yoda1 fed animals. Scale bar, 50 mm, 10 mm. **e**. Quantification of Yki nuclear to cytoplasmic ratio in midguts as in d (n=10,10). **f**. *Expanded* expression (*ex-lacZ*; pink) in midgut trachea (green) from vehicle or Yoda1 fed animals. Scale bar, 50 mm, 10 mm. **g**. Quantification of tracheal *ex-lacZ* expression in midguts as in f (n=18,14). **h, i**. Quantification of tracheal remodelling (h) (n=9,9,11,10) and ISC proliferation (i) (n=21,9, 12,12) upon sucrose or *Pe* infection in wildtype and tracheal *yki* knockdown (*UAS-yki-IR2*) midguts. Dashed lined squares or circles highlight individual tracheal cells. Data represent the mean ± SEM. n=numbers of guts analysed. Statistics: **e, g**. Two-tailed unpaired Student's t-test. **b, h, i**. Two-way ANOVA followed by Tukey's multiple comparisons test.

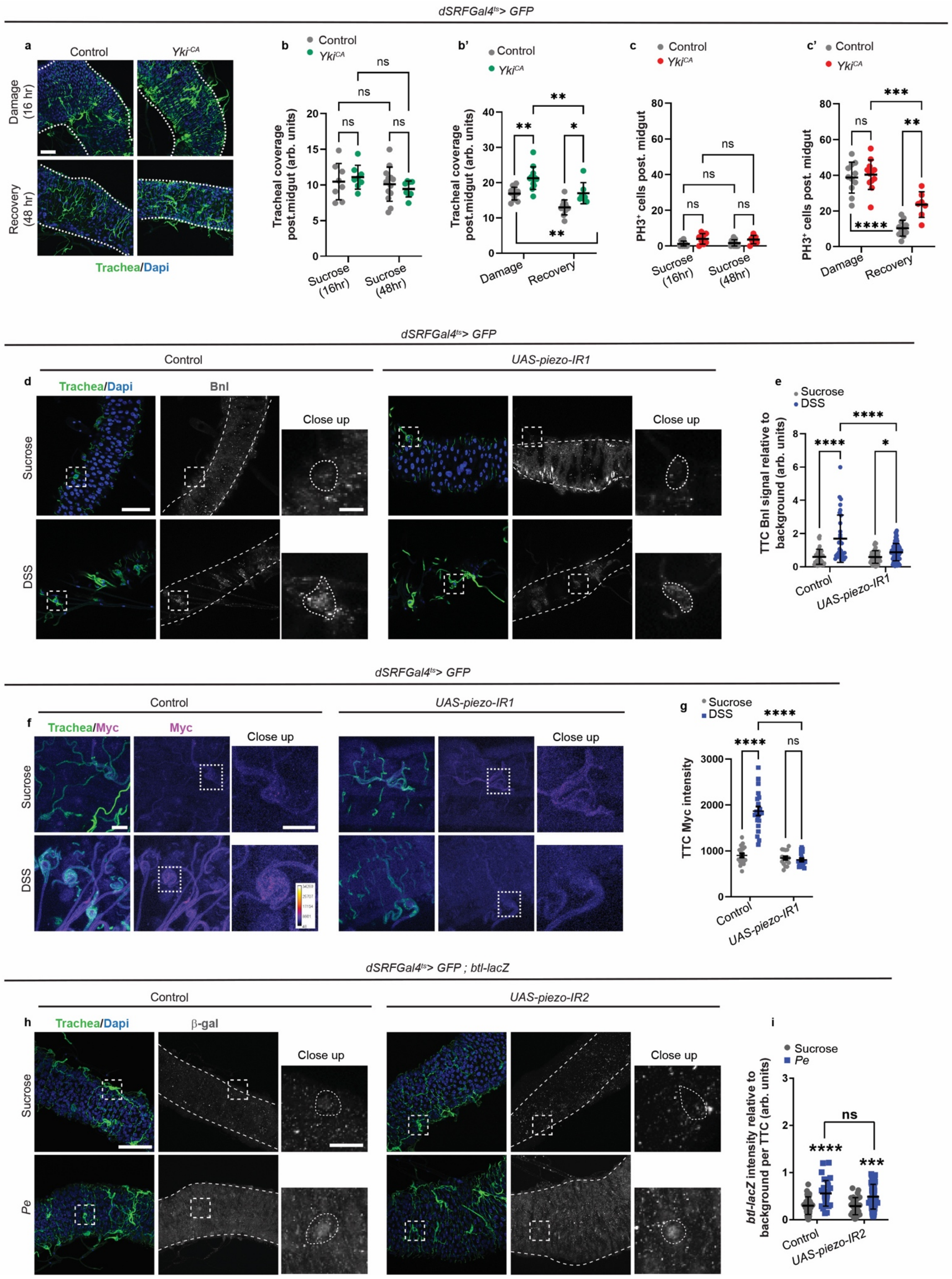
dSRFGal4ts> GFP
a
Control
Yki-CA
Damage (16 hr)
Recovery (48 hr)
Trachea/Dapi
b
Control
YkiCA
ns
ns
ns
ns
Tracheal coverage post.midgut (arb. units)
Sucrose (16hr)
Sucrose (48hr)
b'
Control
YkiCA
**
**
*
**
Tracheal coverage post.midgut (arb. units)
Damage
Recovery
c
Control
YkiCA
ns
ns
ns
ns
PH3+ cells post. midgut
Sucrose (16hr)
Sucrose (48hr)
c'
Control
YkiCA
***
ns
**
****
PH3+ cells post. midgut
Damage
Recovery
dSRFGal4ts> GFP
Control
UAS-piezo-IR1
d
Trachea/Dapi
Bnl
Close up
Sucrose
DSS
e
Sucrose
DSS
****
****
*
TTC Bnl signal relative to background (arb. units)
Control
UAS-piezo-IR1
dSRFGal4ts> GFP
Control
UAS-piezo-IR1
f
Trachea/Myc
Myc
Close up
Sucrose
DSS
g
Sucrose
DSS
****
****
ns
TTC Myc intensity
Control
UAS-piezo-IR1
dSRFGal4ts> GFP ; btl-lacZ
Control
UAS-piezo-IR2
h
Trachea/Dapi
β-gal
Close up
Sucrose
Pe
i
Sucrose
Pe
ns
****
***
btl-lacZ intensity relative to background per TTC (arb. units)
Control
UAS-piezo-IR2

**Supplementary Figure 8. Piezo regulates tissue remodelling pathways in the gut associated trachea. a.** Confocal images of midgut associated trachea (green) without (control) or upon overexpression of constitutively active *Yki* (*Yki-CA*) during 16h *Pe* infection or 48hs after *Pe* removal (recovery). Scale bar, 50 mm. **b, b'.** Quantification of tracheal coverage in sucrose (**b**) or infected and recovery (**b'**) midguts in conditions as in a (n=8,8,13,8) (n=11,12,11,7). **c, c'.** Quantification of ISC proliferation in sucrose (**c**) or infected and recovery (**c'**) midguts as in a (n=8,8,13,8) (n=11,12,11,7). **d**. Confocal images of midgut associated trachea (green) and *bnl* expression (grey) upon sucrose or DSS feeding in control midguts or following tracheal *piezo* knockdown (*piezo-IR1*). Scale bar, 50 mm. **e**. Quantification of *bnl* expression per TTC in midguts as in d (n=44,31,45,87); n=tracheal cells. **f**. Confocal images of midgut trachea (green) and Myc expression (pink) upon sucrose or DSS feeding in control midguts or following tracheal *piezo* knockdown (*piezo-IR1*). Scale bar, 20 mm, 10 mm. **g**. Quantification of Myc expression per TTC in midguts as in f (n=20,20,17,17). **h**. Confocal images of gut trachea (green) and *btl-lacZ* (grey) upon sucrose or *Pe* feeding in control midguts or following tracheal *piezo* knockdown (*piezo-IR1*). Scale bar, 50 mm, 10 mm. **i**. Quantification of average *btl-lacZ* expression per TTC in midguts as in h (n=43,31,34,60); n=tracheal cells. Dashed lined squares or circles highlight individual tracheal cells. Data represent the mean ± SEM. Unless noted, n=numbers of guts analysed. Statistics: Two-way ANOVA followed by Tukey's multiple comparisons test.

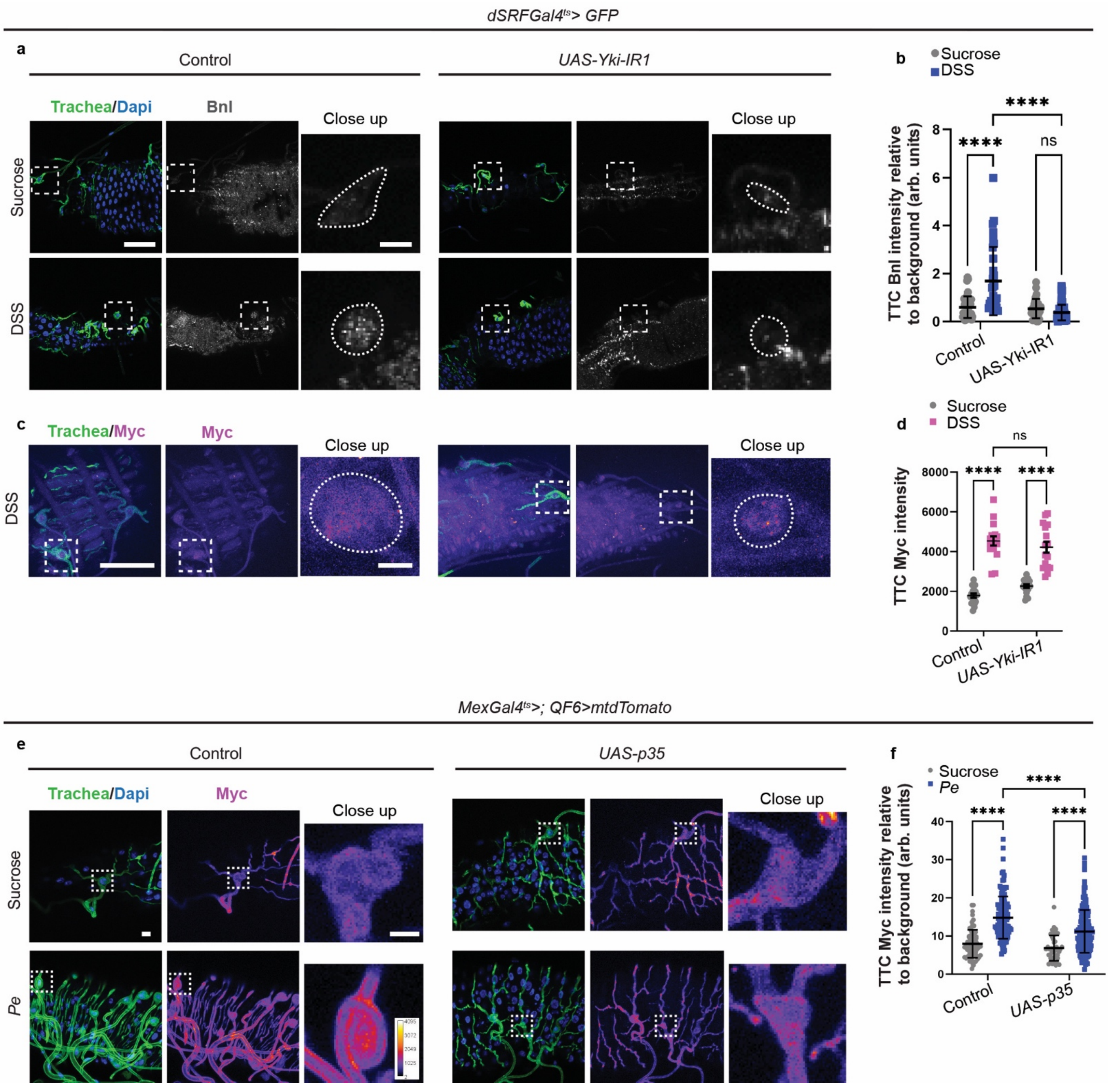

dSRFGal4ts> GFP
a
Control
UAS-Yki-IR1
Trachea/Dapi
Bnl
Close up
Sucrose
DSS
b
Sucrose
DSS
TTC Bnl intensity relative to background (arb. units)
****
****
ns
Control
UAS-Yki-IR1
c
Trachea/Myc
Myc
Close up
DSS
d
Sucrose
DSS
ns
****
****
TTC Myc intensity
Control
UAS-Yki-IR1
MexGal4ts>; QF6>mtdTomato
e
Control
UAS-p35
Trachea/Dapi
Myc
Close up
Sucrose
Pe
f
Sucrose
Pe
****
****
****
TTC Myc intensity relative to background (arb. units)
Control
UAS-p35

**Supplementary Figure 9. Yki regulates tissue remodelling pathways in the gut associated trachea. a.** Confocal images of midgut associated trachea (green) and anti-Bnl expression (grey) upon sucrose or DSS feeding in control midguts or following tracheal *yki* knockdown (*yki-IR1*). Scale bar, 50 mm, 10 mm. **b**. Quantification of Bnl expression per TTC in midguts as in a (n=44,31,28,75); n=tracheal cells. **c**. Confocal images of midgut associated trachea (green) and anti-Myc expression (pink) upon DSS feeding in control midguts or following tracheal *yki* knockdown (*yki-IR1*). Scale bar, 50 mm, 10 mm. **d**. Quantification of Myc expression per TTC in midguts as in c (n=17,16,18,16). **e**. Confocal images of tracheal Myc expression (magenta) in wildtype flies or following *p35* overexpression in enterocytes (*$Mex^{ts}$>p35*) upon sucrose and *Pe* feeding. Scale bar, 10 mm, 10 mm. **f**. Quantification of tracheal Myc expression as in e (n=70,126,42,164); n=tracheal cells. Dashed lined squares or circles highlight individual tracheal cells. Data represent the mean ± SEM. Unless otherwise noted, n=numbers of guts analysed. Statistics: Two-way ANOVA followed by Tukey's multiple comparisons test.

| | | |
|---|---|---|
| **Fig.1** | **a** | *dSRF-Gal4, UAS-mCD8GFP/+; tub-Gal80ts/+* |
| | **b** | *yw, hsFLP; dSRF-Gal4, UAS-mCD8GFP/+; FRT82B, tub-Gal80/FRT82B, LacZ* |
| | **c-h** | *dSRF-Gal4, UAS-mCD8GFP/+; tub-Gal80ts/+* |
| | **i-j** | *dSRF-Gal4, UAS-mCD8GFP/lgl::mcherry; tub-Gal80ts/+* |
| | **k-n'** | *dSRF-Gal4, UAS-mCD8GFP/+; tub-Gal80ts/+* |
| | **o-q** | *dSRF-Gal4, UAS-mCD8GFP/lgl::mcherry; tub-Gal80ts/+* |
| **Fig.2** | **a** | *dSRF-Gal4, UAS-mCD8GFP/+; tub-Gal80ts/+* |
| | **b-f** | *dSRF-Gal4, UAS-mCD8GFP/lgl::mcherry; tub-Gal80ts/+* |
| | **g-j** | *dSRF-Gal4, UAS-mCD8GFP/+; tub-Gal80ts/+* |
| | **k-m** | *Mex-Gal4/Btl-LexAHG,lexAopmCherry; UAS-GFP1-10, lexAop-GFP11/tub-Gal80ts* |
| | **n-r** | *Mex-Gal4, tub-Gal80ts/+; QF6,QUAS-mtdTomato/+* |
| | | *Mex-Gal4, tub-Gal80ts/UAS-MyoII-IR; QF6, QUAS-mtdTomato/+* |
| **Fig.3** | **a,b** | *dSRF-Gal4, UAS-mCD8GFP/+; tub-Gal80ts/+* |
| | **c,d** | *Mex-Gal4, tub-Gal80ts/+; QF6,QUAS mtdTomato/+* |
| | | *Mex-Gal4, tub-Gal80ts/UAS-Bax, tub-Gal80ts; QF6,QUAS mtdTomato/MKRS* |
| | **e-g** | *Mex-Gal4, tub-Gal80ts/+; +/+* |
| | | *Mex-Gal4, tub-Gal80ts/UAS-Bax, tub-Gal80ts; +/+* |
| | **h-i** | *Mex-Gal4/lgl::mcherry; UAS-GC3Ai/MKRS* |
| | **j-m** | *; ;UAS-p35/+* |
| | | *Mex-Gal4,tub-Gal80ts/+; +/+* |
| | | *Mex-Gal4,tub-Gal80ts/+; UAS-p35/+* |
| | **n-p** | *Mex-Gal4, tub-Gal80ts/+; QF6,QUAS mtdTomato/+* |
| | | *Mex-Gal4,tub-Gal80ts/+; QF6,QUAS mtdTomato/UAS-p35* |
| **Fig.4** | **a-i'** | *Pdgf-beta Cre-GFP Piezowt/wt (C57BL/6)* |
| | **j, k** | *Piezo-Gal4/+; UAS-mCD8GFP/QF6,QUAS mtdTomato* |
| | **l, m** | *Piezo-Gal4/+; UAS-mCD8GFP/+* |
| | **n, o** | *Piezo-Gal4/+; UAS-mCD8GFP/QF6,QUAS mtdTomato* |
| | **p-q** | *Mex-Gal4, tub-Gal80ts/piezo::mimic; QF6,QUAS mtdTomato/+* |
| | | *Mex-Gal4, tub-Gal80ts/piezo::mimic; QF6,QUAS mtdTomato/UAS-p35* |
| **Fig.5** | **a-d** | *Piezo-Gal4/+; UAS-mCD8GFP/QF6,QUAS mtdTomato* |
| | **e-g** | *QF6,QUAS mtdTomato/QUAS-GCamp6m* |

| | | |
|---|---|---|
| | **h, i** | *dSRF-Gal4, UAS-mtdTomato/UAS-IVS-jGCaMP7s; +/+* |
| | | *dSRF-Gal4, UAS-mtdTomato/UAS-IVS-jGCaMP7s; UAS-piezo-IR1/+* |
| | **j-n** | *dSRF-Gal4, UAS-mCD8GFP/+; tub-Gal80ts/+* |
| | | *dSRF-gal4, UAS-mCD8GFP/+; tub-Gal80ts/UAS-piezo-IR1* |
| | | *dSRF-gal4, UAS-mCD8GFP/UAS-piezo-IR2; tub-Gal80ts/+* |
| **Fig.6** | **a-p** | *Pdgf-beta Cre-GFP Piezowt/wt (C57BL/6)* |
| | | *Pdgf-beta Cre-GFP Piezofl/fl (C57BL/6)* |
| **Fig.7** | **a-d** | *Piezo-Gal4/+; UAS-mCD8GFP/QF6,QUAS mtdTomato* |
| | **e-g** | *dSRF-Gal4/ Yki::GFP; QF6,QUAS mtdTomato/+* |
| | | *dSRF-Gal4/ Yki::GFP; QF6,QUAS mtdTomato/UAS-piezo-IR1* |
| | **h, i** | *dSRF-Gal4, UAS-mCD8GFP/Ex::LacZ; tub-Gal80ts/+* |
| | **j-k** | *dSRF-Gal4/Ex::GFP; +/QF6,QUAS mtdTomato* |
| | | *dSRF-Gal4/Ex::GFP; UAS-piezo-IR1/QF6,QUAS mtdTomato* |
| | **l-n** | *dSRF-Gal4/piezo::GFP*$^{mimic}$*; +/QF6,QUAS mtdTomato* |
| | | *dSRF-Gal4/piezo::GFP*$^{mimic}$*; UAS-Yki-IR-1/QF6,QUAS mtdTomato* |
| **Fig.8** | **a-c** | *dSRF-Gal4, UAS-mCD8GFP/+; tub-Gal80ts/+* |
| | | *dSRF-Gal4, UAS-mCD8GFP/+; tub-Gal80ts/UAS-Yki-IR1* |
| | **d-f** | *dSRF-Gal4, UAS-mCD8GFP/+; tub-Gal80ts/+* |
| | | *dSRF-Gal4, UAS-mCD8GFP/UAS-piezo-IR2; tub-Gal80ts/+* |
| | | *dSRF-Gal4, UAS-mCD8GFP/+; tub-Gal80ts/UAS-Yki*$^{CA}$ |
| | | *dSRF-Gal4, UAS-mCD8GFP/UAS-piezo-IR2; tub-Gal80ts/UAS-Yki*$^{CA}$ |
| | **g, h** | *dSRF-Gal4, UAS-CD8GFP/+; tub-Gal80ts/bnl-lacZ* |
| | | *dSRF-Gal4, UAS-mCD8GFP/UAS-piezo-IR2; tub-Gal80ts/bnl-lacZ* |
| | **i, j** | *dSRF-Gal4, UAS-mCD8GFP/+; tub-Gal80ts/+* |
| | | *dSRF-Gal4, UAS-mCD8GFP/+; tub-Gal80ts/UAS-piezo-IR1* |
| **Fig.9** | **a-d** | *dSRF-Gal4, UAS-mCD8GFP/+; tub-Gal80ts/+* |
| | | *dSRF-Gal4, UAS-mCD8GFP/+; tub-Gal80ts/UAS-Yki-IR1* |
| **Supplementary Fig.1** | **a-f** | *;+; QF6,QUAS mtdTomato/+* |
| | **g-p** | *dSRF-Gal4, UAS-mCD8GFP/+; tub-Gal80ts/+* |
| **Supplementary Fig.2** | **a-h & j** | *Mex-Gal4/+; tub-Gal80ts/+* |
| | **i** | *Mex-Gal4/+; UAS-GFP1-10, lexAop-GFP11/tub-Gal80ts* |

| | | |
|---|---|---|
| | | *Btl-LexAHG,lexAopmCherry/+; UAS-GFP1-10, lexAop-GFP11/tub-Gal80ts* |
| **Supplementary Fig.3** | **a** | *; ; UAS-p35/+* |
| | | *Mex-Gal4,tub-Gal80ts/+; ;* |
| | | *Mex-Gal4,tub-Gal80ts/+; /UAS-p35/+* |
| | **b-e** | *Mex-Gal4,tub-Gal80ts/+; ;* |
| | | *Mex-Gal4,tub-Gal80ts/+; /UAS-p35/+* |
| | **f-i** | *Pdgf-beta Cre-GFP Piezowt/wt (C57BL/6)* |
| **Supplementary Fig.4** | **a** | *Piezo-Gal4/+; UAS-mCD8GFP/+* |
| | **b, c** | *piezo::GFPmimic/+; QF6,QUAS mtdTomato/+* |
| | **d-h** | *Piezo-Gal4/+; UAS-mCD8GFP/QF6,QUAS mtdTomato* |
| | **i** | *Piezo-Gal4/+; +/+* |
| | | *Piezo-Gal4/+; +/UAS-piezo-IR1* |
| | | *Piezo-Gal4/UAS-piezo-IR2; +/+* |
| | **j-m** | *dSRF-Gal4, UAS-mCD8GFP/+; tub-Gal80ts/+* |
| | | *dSRF-Gal4, UAS-mCD8GFP/+; tub-Gal80ts/UAS-piezo-IR1* |
| | **o-q** | *dSRF-Gal4, UAS-mCD8GFP/+; tub-Gal80ts/+* |
| | | *dSRF-Gal4, UAS-mCD8GFP/UAS-piezo; tub-Gal80ts/+* |
| **Supplementary Fig.5** | **b-i** | *Pdgf-beta Cre-GFP Piezowt/wt (C57BL/6)* |
| | | *Pdgf-beta Cre-GFP Piezofl/fl (C57BL/6)* |
| **Supplementary Fig.6** | **b-e** | *Pdgf-beta Cre-GFP Piezowt/wt (C57BL/6)* |
| | | *Pdgf-beta Cre-GFP Piezofl/fl (C57BL/6)* |
| **Supplementary Fig.7** | **a, b** | *dSRF-Gal4/Ex::GFP; +/QF6,QUAS mtdTomato* |
| | | *dSRF-Gal4/Ex::GFP; UAS-Yki-IR1/QF6,QUAS mtdTomato* |
| | **c** | *dSRF-Gal4/Ex::GFP; +/QF6,QUAS mtdTomato* |
| | | *dSRF-Gal4/Ex::GFP; UAS-piezo-IR1/QF6,QUAS mtdTomato* |
| | **d, e** | *dSRF-Gal4/ Yki::GFP; QF6,QUAS mtdTomato/+* |
| | **f-g** | *dSRF-Gal4, UAS-mCD8GFP/+; Ex::LacZ/tub-Gal80ts* |
| | **h-i** | *dSRF-Gal4, UAS-mCD8GFP/+; tub-Gal80ts/+* |
| | | *dSRF-Gal4, UAS-mCD8GFP/+; tub-Gal80ts/UAS-Yki-IR2* |
| **Supplementary Fig.8** | **a-c'** | *dSRF-gal4, UAS-mCD8GFP/+; tub-Gal80ts/+* |
| | | *dSRF-Gal4, UAS-mCD8GFP/+; tub-Gal80ts/UAS-*$Yki^{CA}$ |
| | **d-g** | *dSRF-Gal4, UAS-mCD8GFP/+; tub-Gal80ts/+* |
| | | *dSRF-Gal4, UAS-mCD8GFP/+; tub-Gal80ts/UAS-piezo-IR1* |

| | | |
|---|---|---|
| | h, i | *dSRF-Gal4, UAS-mCD8GFP/+; tub-gal80ts/btl-lacZ* |
| | | *dSRF-gal4, UAS-mCD8GFP/UAS-piezo-IR2; tub-gal80ts/btl-lacZ* |
| **Supplementary Fig.9** | a-d | *dSRF-Gal4, UAS-mCD8GFP/+; tub-Gal80ts/+* |
| | | *dSRF-Gal4, UAS-mCD8GFP/+; tub-Gal80ts/UAS-Yki-IR1* |
| | e, f | *Mex-Gal4, tub-Gal80ts/+; QF6,QUAS mtdTomato/+* |
| | | *Mex-Gal4, tub-Gal80ts/+; QF6,QUAS mtdTomato/UAS-p35* |

**Supplementary Table 1: Full genotype of animals used in this study.**

| REAGENT or RESOURCE | SOURCE | IDENTIFIER |
|---|---|---|
| Primary antibodies for fly work | | |
| Rat anti-Bnl | S.Hirabayashi | [1] |
| Chicken anti-βeta-Galactosidase | Abcam | ab-9361 |
| Chicken anti-GFP | Abcam | ab-13970 |
| Rabbit anti-DsRed | Clontech | 632496 |
| Guinea pig anti-dMyc | Gift from G.Morata | N/A |
| Rabbit anti-cleaved Dcp1 | Cell Signaling | cat No.9578s |
| Mouse anti-Dlg | DSHB | 528203 |
| Mouse anti-SRF | M. Krasnow | N/A |
| Rabbit anti-PH3 S10 | Cell Signaling | NEB-9701 |
| Mouse anti-PH3 S10 | Cell Signaling | 9706S |
| Primary antibodies for mouse work | | |
| Rabbit anti-Chromogranin A | Abcam | ab15160 |
| Rabbit anti-c-Myc | Abcam | ab32072 |
| Rabbit anti-Caspase 3 cleaved | Cell signalling | 9661L |
| Rabbit anti-Sox-9 | Millipore | AB5535 |
| Rabbit anti-Ki67 | Cell Signalling Technology | 12202 |
| Rat anti-Pecam1 | BD Pharmingen | 557355/MEC13.3 1 |
| Rabbit anti-Ecadherin, 1:500 | Cell Signalling | 24E10 |
| Rat anti-CD31 | BD Pharmingen | 557355 |
| RNAscope *Mm-Piezo1-O1* | ACDBio | 500518 |
| RNAscope *Mm-Pecam1-C2* | ACDBo | 316728-C2 |
| Secondary antibodies | | |
| Goat Alexa Fluor anti-chicken IgY 488 | Invitrogen | A11039 |
| Goat Alexa Fluor anti-chicken IgY 594 | Invitrogen | A11042 |
| Goat Alexa Fluor anti-mouse IgG 488 | Invitrogen | A11029 |
| Goat Alexa Fluor anti-mouse IgG 594 | Invitrogen | A11032 |
| Goat Alexa Fluor anti-mouse IgG 647 | Invitrogen | A21235 |
| Goat Alexa Fluor anti-Rabbit IgG 488 | Invitrogen | A11008 |
| Goat Alexa Fluor anti-Rabbit IgG 594 | Invitrogen | A11034 |
| Donkey Alexa Fluor anti-Rat IgG 594 | Invitrogen | A21209 |
| Opal 520 | Akoya Biosciences | FP1487001KT |
| Opal 570 | Akoya Biosciences | FP1488001KT |
| Chemicals and Reagents | | |
| NAC | Sigma | A7250 |
| Sucrose | Fisher Chemical | S/8600/60 |
| Yoda1 | Tocris | 5586 |
| Methylcellulose | Sigma | 274429 |
| DSS | Sigma | 42867-5G |
| Formaldehyde | FA, EM grade, | 30525-89-4 |
| Tamoxifen | Sigma | T5648 |
| Triton X100 | Sigma | X-100-1L |
| Normal Goat serum | Abcam | ab7481 |
| Vectashield with DAPI | Vector Laboratories | H-1200 |
| Bovine Serum Albumin | Sigma | A3294-50G |
| Schnieder's Medium | Sigma | 50146 |
| Fly Extract | VDRC | |
| Pen-Strep | Invitrogen | 15140 |
| Whatman GF/B Glass microfiber filters 21mm | Cytiva | 1821-021 |
| Corn Oil | Sigma | C8267 |

| | | |
|---|---|---|
| QuikSil | World Precision Instruments | |
| DHE | Invitrogen | D23107 |
| DMSO | ThermoFisher | J66650.AK |
| Cell-Tak | Sigma | CLS354240 |
| Low gelling temperature agarose | Sigma | A4018 |
| FBS | Sigma | F4135 |
| Ecomount | Bio-Techne | 320409 |
| Trizol | Thermo Fischer Scientific | Cat No. 15596018. |
| Chloroform | Merck Life Sciences | Cat No. 372978. |
| Iso-propanol | Merck Life Sciences | Cat No. I9516. |
| DNase I | Merck Life Sciences | Cat No. AMPD1-1KT |
| high-capacity cDNA reverse transcription kit | Thermo Fisher Scientific | Cat No. 4833950 |
| Perfecta SYBR Fastmix | VWR | Cat No.733-1390 |
| Sodium Azide | Sigma | 26628-22-8 |
| Oligonucleotides | | |
| *rpl32 Forward: AGGCCCAAGATCGTGAAGAA* | This study | N/A |
| *pl32 Reverse: TGTGCACCAGGAACTTCTTGAA* | This study | N/A |
| *Piezo Forward primer CTCACGCTGCCCATCTACAA* | This Study | N/A |
| *Piezo Reverse primer GTGGCGAAAACCAACACCTC* | This Study | N/A |
| Experimental Models: mouse strains | | |
| *Pdgfb-CreER* | Marcus Fruttiger | [2] |
| *Piezo1* $^{fl/fl}$ | Jackson Laboratories | 029213 |
| *C57Bl6J* | Biological Service Unit CRUK SI | N/A |
| Experimental Models: fly strains | | |
| *Wild type strain (w* $^{1118}$*)* | Gift from Ross Cagan | N/A |
| *dSRF-gal4, UAS-CD8GFP* ; tub-gal80ts | Gift from Irene Miguel-Aliaga | [3] |

| | | |
|---|---|---|
| *piezo-gal4* | Gift from Norbert Perrimon | [4] |
| *QF6, QUAS mtdTomato* | Gift from Chrysoula Pitsouli | [5] |
| *Mex-gal4, tub-gal80* $^{ts}$ | Gift from Irene Miguel-Aliaga | N/A |
| *UAS-piezo-IR1* | Vienna Drosophila Resource Centre | 2796 |
| *UAS-piezo-IR2* | Vienna Drosophila Resource Centre | 105132 |
| *btl-lacZ* | Bloomington Drosophila Stock Center | 12086 |
| *bnl-lacZ* | Bloomington Drosophila Stock Center | 11704 |
| *Yki::GFP* | Gift from Barry Thompson | [6] |
| *UAS-GC3Ai* | Gift from Andrew Davidson | N/A |
| *UAS-Yki-IR1* | Bloomington *Drosophila* Stock Center | 31965 |
| *UAS-Yki-IR2* | Bloomington *Drosophila* Stock Center | 34067 |
| *lgl::mcherry* | Gift from Eurico Morais-de-Sá | N/A |
| *btl-LexA* | Gift from Thomas Kornberg | [7] |
| *UAS-GFP1-10, lexAop-GFP11* | Gift from Thomas Kornberg | [7] |
| *lexAop-Cherry-CAAX* | Gift from Thomas Kornberg | [7] |
| *QUAS-GCamp6m* | Bloomington *Drosophila* Stock Center | 91988 |
| *Piezo::GFP mimic* | Bloomington *Drosophila* Stock Center | 60209 |
| *UAS-Piezo-EGFP* | Bloomington *Drosophila* Stock Center | 58772 |
| *UAS-bax, tubgal80* $^{ts}$ | Gift from Cedric Polesello | N/A |
| *ExLacZ* | Gift from Nic Tapon | N/A |

| | | |
|---|---|---|
| *ExGFP* | Gift from Nic Tapon | N/A |
| *UAS-MyoII-IR* | Vienna Drosophila Resource Centre | 107177 |
| *UAS-Yki $^{CA}$ (III)* | Gift from Norbert Perrimon | N/A |
| *UAS-p35* | Bloomington *Drosophila* Stock Center | 5073 |
| *UAS-mCD8GFP* | Bloomington *Drosophila* Stock Center | 5137 |
| *UAS-IVS-jGCaMP7s* | Bloomington *Drosophila* Stock Center | 80905 |
| Software | | |
| Biorender | https://www.biorender.com/product/leo-ai-figure-builder | |
| Graph Pad Prism 6.07 | www.graphpad.com | |
| ImageJ 1.54r | http://imagej.nih.gov/ij | |
| Carl Zeiss Zen 2.0 (blue edition) Version 2.0.0.0 | https://www.zeiss.com | |
| Carl Zeiss Zen 3.0 Version 16.0.1.306 | https://www.zeiss.com | |
| Bayesian Force Inference | https://github.com/IshiharaLab/BayesianForceInference | |
| Deproj | https://github.com/Image-Analysis-Hub/DeProj | |
| LocalZprojector | https://github.com/Image-Analysis-Hub/LocalZProjector | |
| Cellpose | https://github.com/mouseland/cellpose | |
| Deproj | https://github.com/Image-Analysis-Hub/DeProj | |
| Nanoprepare | https://github.com/massimovassalli/NanoPrepare | |
| 7500 Real-Time PCR Software | Applied Biosystems | |
| Softmech | https://github.com/CellMechLab/softmech | |
| QUPath | https://qupath.github.io | |

**Supplementary Table 2: Full list of resources used in this study.**

### Supplementary References

1. Newton H, Wang Y-F, Camplese L, Mokochinski JB, Kramer HB, Brown AEX, et al. Systemic muscle wasting and coordinated tumour response drive tumourigenesis. Nature Communications. 2020 2020/09/16;11(1):4653.

2. Claxton S, Kostourou V, Jadeja S, Chambon P, Hodivala-Dilke K, Fruttiger M. Efficient, inducible Cre-recombinase activation in vascular endothelium. genesis. 2008;46(2):74-80.

3. Gervais L, Casanova J. The Drosophila homologue of SRF acts as a boosting mechanism to sustain FGF-induced terminal branching in the tracheal system. Development. 2011 Apr;138(7):1269-74. PubMed PMID: 21385762. Epub 2011/03/10. eng.

4. He L, Si G, Huang J, Samuel ADT, Perrimon N. Mechanical regulation of stem-cell differentiation by the stretch-activated Piezo channel. Nature. 2018;555(7694):103-6.

5. Tamamouna V, Rahman MM, Petersson M, Charalambous I, Kux K, Mainor H, et al. Remodelling of oxygen-transporting tracheoles drives intestinal regeneration and tumorigenesis in Drosophila. Nature Cell Biology. 2021;23(5):497-510.

6. Fletcher GC, Diaz-de-la-Loza MD, Borreguero-Muñoz N, Holder M, Aguilar-Aragon M, Thompson BJ. Mechanical strain regulates the Hippo pathway in Drosophila. Development. 2018 Mar 8;145(5). PubMed PMID: 29440303. PMCID: PMC5868995. Epub 2018/02/15. eng.

7. Roy S, Huang H, Liu S, Kornberg TB. Cytoneme-mediated contact-dependent transport of the Drosophila decapentaplegic signaling protein. Science. 2014 Feb 21;343(6173):1244624. PubMed PMID: 24385607. PMCID: PMC4336149. Epub 2014/01/05.
